# A universal cannabinoid CB1 and CB2 receptor TR-FRET kinetic ligand binding assay

**DOI:** 10.1101/2024.07.16.603654

**Authors:** Leire Borrega-Roman, Bradley L. Hoare, Miroslav Kosar, Roman C. Sarott, Kacper J. Patej, Jara Bouma, Morgan Scott-Dennis, Eline J. Koers, Thais Gazzi, Leonard Mach, Sergio Barrondo, Joan Sallés, Wolfgang Guba, Eric Kusznir, Marc Nazaré, Arne C. Rufer, Uwe Grether, Laura H. Heitman, Erick M. Carreira, David A. Sykes, Dmitry B. Veprintsev

## Abstract

**INTRODUCTION:** The kinetics of ligand binding to G protein-coupled receptors (GPCRs) is an important determining factor in the preclinical evaluation of a molecule. Therefore, efforts should be made to measure this property as part of any drug development plan. The original assays used to assess ligand binding kinetics were developed using radioligands. However, these types of assays are very labor-intensive, limiting their application to the later phases of the drug discovery process. Recently, fluorescence-based ligand binding assays have been developed for multiple GPCRs, demonstrating their superiority through a homogeneous format and continuous data acquisition capabilities. The overriding aim of this study was to develop a fluorescence-based homogeneous ligand binding assay to profile the kinetics of compounds binding to human cannabinoid type 1 and 2 receptors (CB1R and CB2R).

**METHODS:** We designed and synthesized D77, a novel universal tracer based on the lower affinity non-selective naturally occurring psychoactive cannabinoid, Δ^8^-THC. Using the TR-FRET (time-resolved Förster resonance energy transfer) technique to develop an assay to study the kinetics of ligand binding to CB1R and CB2R at physiological temperature. To establish a CB1R construct suitable for this assay, it was necessary to truncate the first 90 amino acids of the flexible CB1R N-terminal domain, in order to reduce the FRET distance between the terbium cryptate (donor) and the fluorescent ligand (acceptor), while the full length CB2R construct remained functional due to its shorter N-terminus. We then used the Motulsky-Mahan competition binding model to study the binding kinetics of non-fluorescent ligands.

**RESULTS:** D77 tracer displayed affinity for the truncated human CB1R (CB1R_91-472_) and full length CB2R (CB2R_1-360_) in the nanomolar range, and competitive binding behavior with orthosteric ligands. Crucially, D77 displayed fast dissociation kinetics from both CB1R and CB2R, comparable to those of the most rapidly dissociating reference compounds tested. This unique property of D77 proved pivotal to accurately determining the on- and off-rates of the fastest dissociating compounds. Using D77, we successfully determined the kinetic binding properties of a series of CB1R and CB2R agonists and antagonists at 37°C, including rimonabant, which was marketed for the treatment of obesity but later withdrawn due to serious neurological side effects.

**DISCUSSION:** The *k*_on_ values of molecules binding CB1R showed a difference of three orders of magnitude from the slowest associating compound, HU308 to the most rapid, rimonabant. Interestingly, we found a strong correlation between *k*_on_ and affinity for compounds binding to CB1R, suggesting that the association rate is the main parameter determining the affinity of compounds binding to CB1R. For compounds binding to CB2R, both *k*_on_ and *k*_off_ parameters contributed as affinity determinants. However, in contrast to CB1R, a stronger correlation was found between the dissociation constant rate parameter and the affinity of these molecules, suggesting that a combination of *k*_on_ and *k*_off_ dictates the overall affinity of compounds binding to CB2R. Ultimately, exploring the kinetic parameters of potential cannabinoid drug candidates could help future drug development programs targeting these receptors.

## Introduction

Cannabinoid type 1 and 2 receptors (CB1R and CB2R) are essential signalling elements of the endocannabinoid system. They belong to the G protein-coupled receptor (GPCR) family and show different distribution patterns in the human body. CB1R is predominantly expressed in the central nervous system (CNS) (Katona, Sperlágh et al. 1999), but it is also found in peripheral organs including adipose tissue, liver, and the pancreas, where it is thought to regulate metabolic functions(Pacher, Bátkai et al. 2006). The CB2R is primarily expressed in peripheral tissues related to the immune system (Khurana, Mackie et al. 2017), such as the spleen and thymus (Munro, Thomas et al. 1993, Galiègue, Mary et al. 2005), but it is also found in the CNS, although expressed at much lower levels compared to CB1R. CB1R are most commonly targeted for their neuromodulatory action and effects on metabolic regulation, finding use in the treatment of obesity, whilst the CB2R is typically targeted for its ability to modulate inflammatory processes (Ashton and Glass 2007, Aso and Ferrer 2016, Kaur, R. Ambwani et al. 2016, Vuic, Milos et al. 2022), with further potential to treat or ameliorate certain neurodegenerative disorders (Benito, Núñez et al. 2003, Gómez-Gálvez, Palomo-Garo et al. 2016, Navarro, Morales et al. 2016).

Phytocannabinoids such as extracts of the plant *C. sativa* have been used in traditional medicine for millennia (Zuardi 2006, Zuardi 2008). Even though the cannabinoid system has been considered a promising intervention point for a plethora of diseases (Di Marzo 2018, Maccarrone, Di Marzo et al. 2023), very few drugs targeting CB1R or CB2R have reached the clinic to date. Currently, dronabinol and nabilone are the only two FDA approved synthetic cannabinoids, which are Δ^9^-THC or close derivates. These are non-selective compounds used for treating nausea and anorexia derived from cancer chemotherapy treatments. One example of a selective CB1R ligand is the antagonist rimonabant, which was marketed as an effective anti-obesity drug. However, it was subsequently withdrawn from the market due to its adverse neuropsychiatric effects (Moreira and Crippa 2009).

Difficulties targeting CB1R and CB2R for therapeutic benefit can be partly explained by their wide distribution in the body, and their role as modulators of multiple processes that are primarily driven by other signalling cascades. The challenge is to design selective drug candidates, that possess appropriate pharmacological properties, which may not readily reveal themselves through mere affinity measures. In this regard there is significant potential to improve the pharmacological properties of compounds by optimising their residency times. The study of drug receptor-binding kinetics can ultimately lead to a more complete understanding of drug action, both *in vitro* and *in vivo*, helping to rationalise improvements in clinical performance, by relating them directly to measures of drug efficacy, rebinding, or off target toxicity based on improvements in kinetic selectivity (Fuchs, Breithaupt-Grögler et al. 2000, Casarosa, Bouyssou et al. 2009, Tresadern, Bartolome et al. 2011, Fleck, Hoare et al. 2012, Sykes, Dowling et al. 2012, Sykes, Moore et al. 2017, Sykes, Jimenez-Roses et al. 2022). Thus it is important to implement these screens early on in drug discovery programs as this information becomes key to rational drug design (Guo, Heitman et al. 2016). Ligand-receptor binding kinetic studies enable improved drug design by providing accurate predictions of drug-receptor target coverage *in vivo*, which in turn lead to improved therapeutic outcomes. Even though our understanding of cannabinoid receptor pharmacology has dramatically increased during the last few decades, the binding kinetics of compounds acting at either of these receptor subtypes under physiologically relevant conditions remain largely unexplored.

Traditionally, drug-receptor binding properties have been investigated by means of radiolabeled ligands, with excellent progress in developing selective ligands for CB2R (Dowling and Charlton 2009, Sykes, Dowling et al. 2010, Ramsey, Attkins et al. 2011, Martella, Sijben et al. 2017, Soethoudt, Hoorens et al. 2018). However, these methods are very labor-intensive and present many limitations such as lower throughput, the risk associated with the handling of hazardous radioactive material and the increased time and costs that these experimental procedures incur, and an inability to study fast binding events (Martella, Sijben et al. 2017, Xia, de Vries et al. 2017, Xia, de Vries et al. 2018, Georgi, Dubrovskiy et al. 2019, Sykes, Jain et al. 2019). Another disadvantage of radioligands is the higher levels of non-specific binding exhibited by certain probes, even when they possess high affinity for the primary target. The use of fluorescence-based methods and resonance energy transfer in particular, has helped to revolutionize the study of ligands binding to GPCRs, its main advantage being the lower levels of non-specific binding due to the requirement for proximity between the donor and acceptor in the generation of the specific binding signal (Sykes, Stoddart et al. 2019, Soave, Briddon et al. 2020). However, despite increases in the number of fluorescent probes targeting CB1 and CB2R, there is currently no fluorescent ligand binding assay described for CB1R and CB2R suitable for profiling the kinetic properties of low affinity rapidly dissociating compounds.

Herein, we report a novel homogeneous time-resolved Förster resonance energy transfer (TR-FRET) competition association binding assay using the novel fluoroprobe D77, a nitrobenzoxadiazole (NBD) labelled tracer, based on the pharmacophore embedded in Δ^8^-THC. The use of this tracer allowed us to characterize CB1R and CB2R ligand kinetic parameters (association rate constant *k*_on_ and dissociation rate constant *k*_off_) at 37°C, using the kinetic model of drug-receptor competition binding proposed by Motulsky and Mahan (Motulsky and Mahan 1984). Using a set of reference compounds for CB1R and CB2R, we present a robust method to perform high-throughput *in vitro* screening of cannabinoid compounds to assess both their affinity and ligand binding kinetics, and to explore the basis of selectivity between these two main cannabinoid receptors.

## Materials and methods

### Materials

T-Rex^TM^-293 cells (Invitrogen) cells were obtained from ThermoFisher Scientific. Culture flasks T75 and T175 cm2, were purchased from ThermoFisher Scientific. DMEM (high glucose), Dulbecco’s Phosphate Buffered Saline (DPBS), no calcium, no magnesium (D8537) was purchased from Sigma-Aldrich. CellStripper™ was purchased from Corning. Hanksʹ Balanced Salt solution (H8264), HEPES (4-(2-hydroxyethyl)-1-piperazineethanesulfonic acid), EDTA (ethylenediamine tetraacetic acid), bovine serum albumin (BSA) heat shock fraction, protease-free, fatty acid-free, essentially globulin free (A7030), poly-D-lysine, tetracycline and Pluronic F127 were purchased from Sigma-Aldrich. The transfection reagent PEI linear, MW 25000, transfection grade (PEI 25K) was obtained from Polysciences (23966-1). The selection reagents blasticidin, geneticin (G418) and zeocin were obtained from Invitrogen. A bicinchoninic acid (BCA) protein assay kit, used to determine the total protein content of membranes was obtained from ThermoFisher Scientific. 2-AG, AEA, rimonabant, SR 144528, HU-308, and HU-210 were obtained from Tocris Bioscience (UK). CP 55,940 was obtained from Sigma-Aldrich. All ligands were dissolved in 100% DMSO and stored as aliquots at −20°C until required. Dimethyl sulfoxide (DMSO, 276855) was purchased from Sigma-Aldrich. OptiPlate-384 (White Opaque 384-well Microplate), were purchased from PerkinElmer (Beaconsfield, UK).

### Cell culture and membrane preparation

T-Rex^TM^-293 cells were used to stably express human CB1R and CB2R with a SNAP-tag genetically encoded at the N-terminus of the receptor. The sequence of the human CB1R was modified, introducing a truncation of the first 90 amino acids of the N-terminus (SNAP-CB1R_91-472_). The SNAP-tagged CB2 receptor was expressed without modifications in the sequence. To generate the stable cell lines, pcDNA4 plasmids were used to transfect the cells and blasticidin (10 μg/ml) and zeocin (20 μg/ml) antibiotics were used for selection for 14 days.

T-Rex^TM^-293 cells stably expressing either SNAP-CB1R_91-472_ or SNAP-CB2 were cultured in T175 flasks using DMEM supplemented with 10% fetal bovine serum. Tetracycline (1 μg/ml) was added to the culture 48h prior to labeling to induce receptor expression. When the cells reached 90-100% confluency, medium was removed and 10 ml of Tag-lite labeling medium (LABMED, Cisbio) containing 100 nM of SNAP-Lumi4-Tb was added and incubated for 1h at 37°C under 5% CO_2_. After a single wash with LABMED, PBS was used to wash the cells twice and cells were dissociated using non-enzymatic dissociation buffer. Cells were centrifuged for 5 min (350 × *g*) and pellets were kept at −80 °C until membranes were prepared.

For membrane preparation, all steps were conducted at 4 °C to avoid tissue degradation. Cells pellets were thawed and resuspended using ice-cold buffer containing 10 mM HEPES and 10 mM EDTA, pH 7.4. The suspension was homogenized using an electrical homogenizer Ultra-Turrax (Ika-Werk GmbH, Germany) and subsequently centrifuged at 1200 *× g* for 5 min. The pellet obtained then, containing cell nucleus and other heavy organelles was discarded and supernatant was centrifuged for 30 min at 48,000 *× g* at 4 °C (Beckman Avanti J-251 Ultra-centrifuge; Beckman Coulter). The supernatant was discarded, and the pellet was resuspended using the same buffer (10 mM HEPES and 10 mM EDTA, pH 7.4) and centrifuged a second time for 30 min as described above. Finally, the supernatant was discarded, and the pellet resuspended using ice-cold 10 mM HEPES and 0.1 mM EDTA, pH 7.4. Protein concentration determination was carried out using the bicinchoninic acid assay kit (Sigma-Aldrich) and using BSA as a standard. The final membrane suspension was aliquoted and maintained at −80°C until required for the assays.

### Common procedures used in TR-FRET experiments

Experiments were performed using T-Rex^TM^-293 cell membranes expressing SNAP-tagged human CB1R and CB2R (SNAP-CB1R_91-472_ and SNAP-CB2). All the assays were conducted at either 25 or 37°C in white 384-well Optiplate plates (PerkinElmer) using the PHERAstar FSX microplate reader (BMG Labtech). Assay buffer comprised HBSS (Hanks Balance Salt Solution) containing 5mM HEPES, 0.5% BSA and 0.02% pluronic F-127, PH 7.4. Membranes were preincubated with 400 μM of GppNHp prior to addition to the plate and automatic injectors were used to ensure early time points could be measured with accuracy. Non-specific binding (NSB) at CB1R and CB2R was determined in the presence of saturating concentrations of rimonabant (3 μM) and SR 144528 (1 μM) respectively.

### Saturation binding studies

Fluorescent ligand binding to the CB1R and CB2R receptor was assessed by HTRF (homogeneous time-resolved FRET) detection allowing the construction of saturation binding curves. For fluorescent ligand characterization, six concentrations of D77 were chosen ranging from 31.25 to 1000 nM. D77 was added simultaneously (at t = 0) to cell membranes containing the human CB1R and CB2R (1 μg per well) in a total volume of 40 μL containing 2% DMSO, at 25 or 37°C, with gentle agitation. NSB was determined in the presence of saturating concentrations of either rimonabant (3 μM) or SR 144528 (1 μM). The resulting data were fitted to the one-site model equation (Equation 1) to derive a single best-fit estimate for *K*_D_, as described under Data analysis.

### Determination of affinity constants (K_i_)

To obtain affinity estimates of unlabelled ligand, D77 competition experiments were performed at equilibrium. D77 was used at a concentration of 600 nM and 900 nM in binding assays, for CB1R and CB2R respectively. D77 was incubated in the presence of the indicated concentration of unlabelled ligand and CB1 and CB2R cell membranes (1 μg/well) in a total volume of 40 μL containing 2% DMSO at 25 or 37°C, with gentle agitation. NSB was determined in the presence of saturating concentrations of either rimonabant (3 μM) or SR 144528 (1 μM). Steady state competition curves were obtained after 15 min incubation and data was fitted using GraphPad Prism 9.2 to the one site competition binding model (Equation 2) to calculate IC_50_ values, which were converted to *K*_i_ values by applying the Cheng-Prusoff correction as described under Data analysis.

### Determination of the association rate (k_on_) and dissociation rate (k_off_) of fluorescent ligands

Fluorescent ligand binding to the CB1R and CB2R receptor was assessed by HTRF (homogeneous time-resolved FRET) detection allowing the construction of association binding curves. For fluorescent ligand characterization, six increasing concentrations of fluorescent ligands were added (at t = 0) to cell membranes containing the human CB1R and CB2R (1 μg/well) in a total volume of 40 μL containing 2% DMSO at 25 or 37°C, with gentle agitation. NSB was determined in the presence of saturating concentrations of either rimonabant (3 μM) or SR 144528 (1 μM). The resulting data were globally fitted to the association kinetic model (Equation 3) to derive a single best-fit estimate for *K*_D_, *k*_on_ and *k*_off_ as described under Data analysis.

### Competition binding kinetics

Competitive kinetic association experiments were carried out using the D77 fluorescent ligand as a tracer. D77 was added simultaneously with the unlabelled compound (at t = 0) to cell membranes containing the human CB1R and CB2R (1 μg/well) in a total volume of 40 μL containing 2% DMSO at 25 or 37°C, with gentle agitation. The concentration of tracer used in the experiments was 600 nM and 900 nM for CB1R and CB2R respectively, concentrations which avoid ligand depletion in this assay volume. The degree of D77 bound to the receptor was assessed at several time points from 0 to 15min. NSB was determined in the presence of saturating concentrations of either rimonabant (3 μM) or SR 144528 (1 μM) and was subtracted from each time point. The association of the D77 was monitored using TR-FRET in the competition with 10 different concentrations of each cannabinoid ligand. The resulting data were globally fitted in GraphPad Prism 9.2 to the “kinetics of competitive binding” model (Equation 4) to derive a single best-fit estimate of *K*_D_, *k*_on_ and *k*_off_ for the different cannabinoid compounds as described under Data analysis.

### Data analysis

Data are reported in the text and tables are mean ± SEM for the indicated number of experiments. All experiments were analyzed by non-linear regression using Prism 9.2 (GraphPad Software, San Diego, U.S.A.).

### Saturation binding

D77 total and non-specific binding data were analyzed by non-linear regression according to a one-site binding equations, and individual estimates for total receptor number (*B*_max_) and ligand dissociation constant (*K*_D_) were calculated. The following one-site model equations were used, where [A] is the concentration of the ligand:

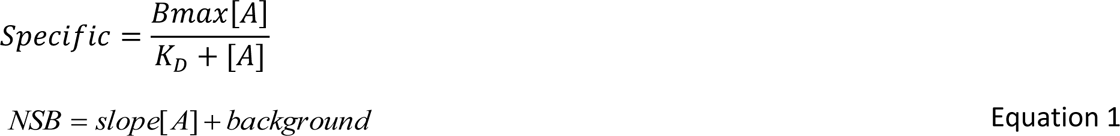

### Competition binding

Steady state competition displacement binding data were fitted to sigmoidal (variable slope) curves using a “four parameter logistic equation”:

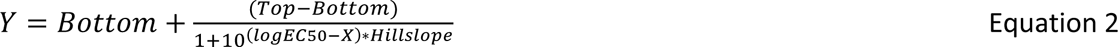

Where, Bottom and Top are the plateaus of the agonist and inverse agonist curves. LogEC_50_ is the concentration of agonist/inverse agonist that gives a half-maximal effect, and the Hillslope is the unitless slope factor or Hillslope. IC_50_ values obtained from the inhibition curves were converted to *K*_i_ values using the method of Cheng and Prusoff (Cheng and Prusoff 1973).

### Association binding

D77 association data was fitted as follows to a global fitting model using GraphPad Prism 9.0 to simultaneously calculate *k*_on_ and *k*_off_ using the following equation where *k*_ob_ equals the observed rate of association and L is the concentration of D77:

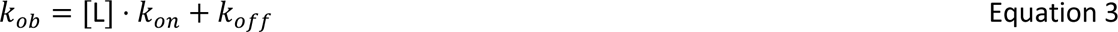

### Competition kinetic binding

Association rate *k*_on_ and dissociation rate *k*_off_ were calculated for the different cannabinoid compounds were obtained by global fitting of competition binding data using the model “kinetics of competitive binding” in GraphPad Prism 9.2:

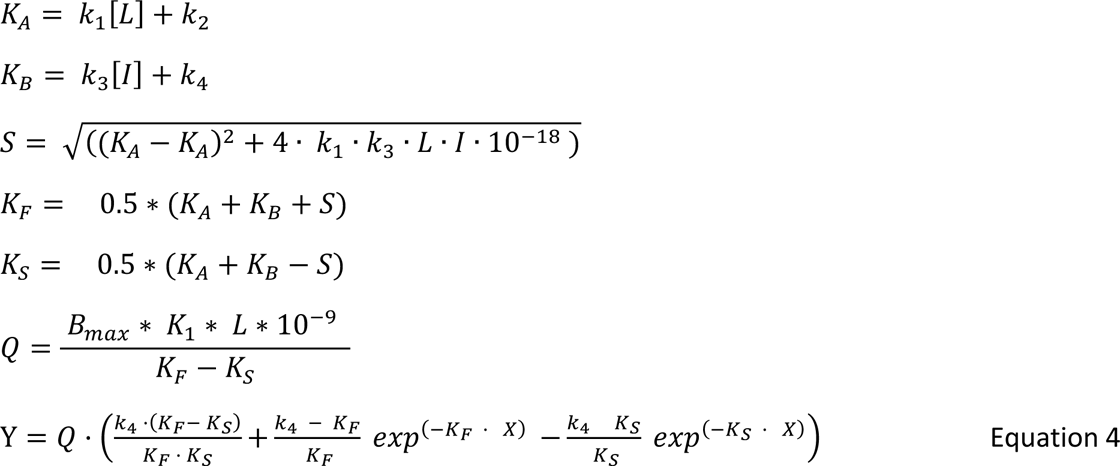

where X is time (min), Y is specific binding (HTRF units 520nm/620nm*10,000), *k*_1_ is *k*_on_ of the tracer D77, *k*_2_ is *k*_off_ of the tracer D77, *L* is the concentration of D77 used (nM) and *I* is the concentration of unlabeled agonist (nM). Fixing the above parameters allowed the following to be simultaneously calculated: B_max_ is total binding (HTRF units 520nm/620nm*10,000), *k*_3_ is association rate of unlabeled ligand (M^−1^ min^−1^) or *k*_on_, and *k*_4_ is the dissociation rate of unlabeled ligand (min^−1^) or *k*_off_.

### Linear correlations

The correlation between datasets was determined by calculating a Pearson correlation coefficient (presented as the r^2^ coefficient of determination, which shows percentage variation in y which is explained by all the x variables together) in GraphPad Prism 9.2.

### Synthesis of fluorescent probes

The details of synthetic procedures as well as compound characterizations can be found in the Supplementary Information.

## Results

### Developing a TR-FRET binding assay for SNAP-CB1 required truncation of its N-terminus

We have previously developed a TR-FRET based binding assay for human CB2R using a genetically encoded SNAP-tag at the N-terminus of the full length CB2R along with complementary CB2 fluorescent probes (Sarott, Westphal et al. 2020, Gazzi, Brennecke et al. 2022, Kosar, Sykes et al. 2023). While these compounds showed good selectivity for CB2, as measured in radio-ligand binding assays, they also bound CB1R albeit with reduced affinity. However, we found that we could not obtain a TR-FRET signal for CB1R.

We hypothesized that the 111 amino acid residue long N-terminus of CB1R, as opposed to the 33-residue short N-terminus of CB2R, places the SNAP-Lumi4 donor too far from the fluorescent ligand bound in the orthosteric binding site (Figure 1A) for efficient FRET, a distance of approximately 60-70 Å (or 6-7 nm). We therefore truncated the CB1 N-terminus to place the SNAP tag closer to the binding pocket. The truncation site was rationally chosen based on knowledge of reported CB1R splicing variants, which have modified N-terminal domains (NTD) and retain functionality (Xiao, Jewell et al. 2008). Two truncated CB1R variants were produced and named based on the residues remaining after truncation-CB1R_55-472_ and CB1R_91-472_ (Figure 1B), both also containing a N-terminal SNAP tag for terbium cryptate labelling. In TR-FRET experiments which tested for specific binding of 1 µM NBD-691, only the most truncated receptor, CB1R_91-472_, displayed a specific TR-FRET signal indicative of ligand binding (Figure 1C).

**Figure 1.**
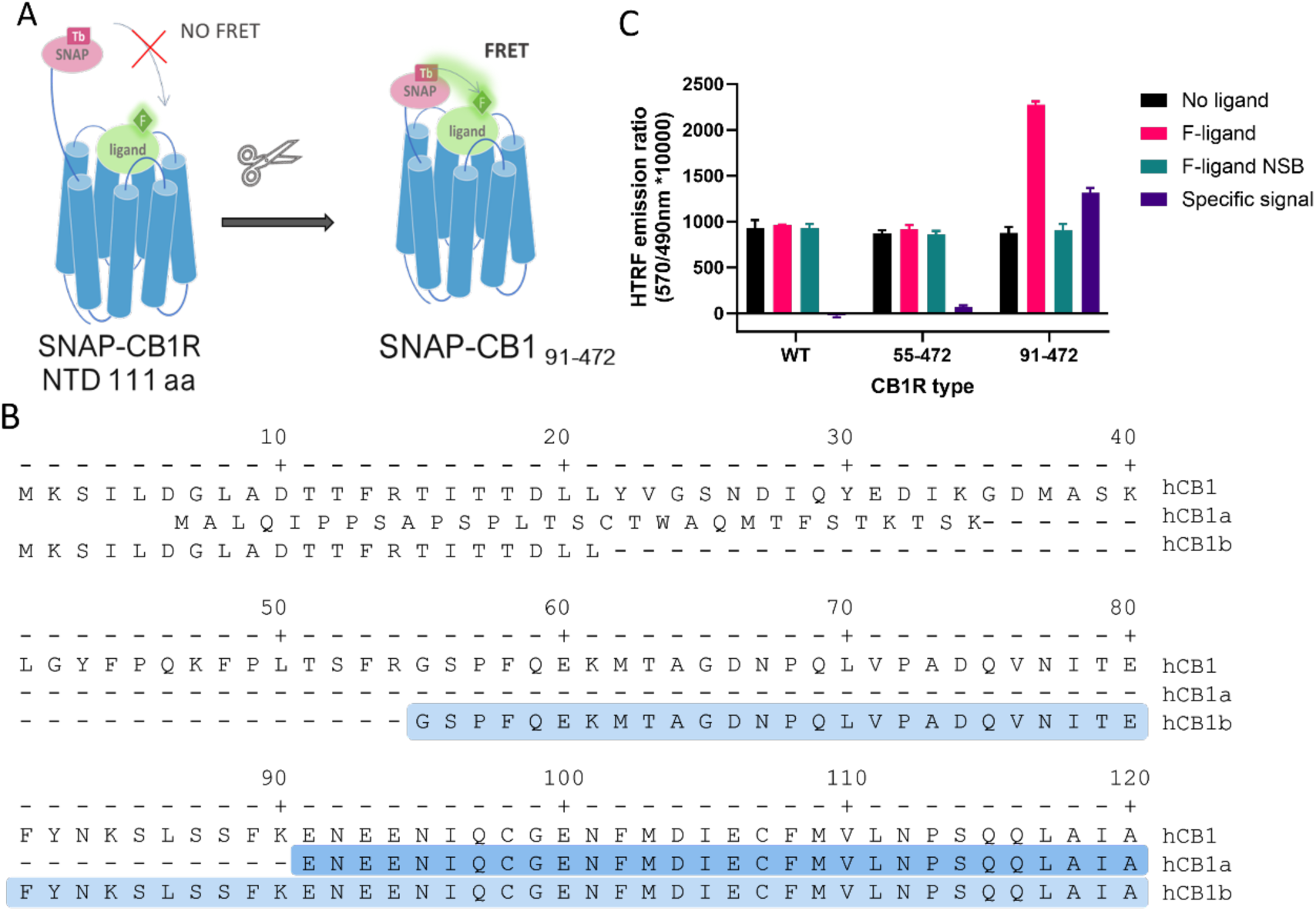
Truncation of human cannabinoid receptor type 1 (CB1R) allows specific TR-FRET binding signal from fluorescent tracers. **(A)** Diagram showing how the long N-terminus of CB1R may preclude FRET between terbium cryptate donor and fluorescent tracer, and rationale for truncating to shorten the distance. **(B)** Amino acid sequence of the N-terminal of human CB1R and its splicing variants hCB1a and hCB1b. The highlighted amino acids represent the N-terminal sequence of the truncated versions used (91-472 in dark blue, 55-472 in light blue, modified from (Ryberg, Vu et al. 2005). **(C)** TR-FRET binding signal at full length and truncated CB1R obtained using NBD-691 (1 µM).

### Truncated receptor retains intact cell-surface localization and functionality

Earlier reports suggested the N-terminal domain of the CB1R may be playing a role in receptor trafficking and stability (Hebert-Chatelain, Desprez et al. 2016, Fletcher-Jones, Hildick et al. 2020). In our case, we observed no difference between the sub-cellular localization of SNAP-CB1R constructs. Most likely, the signal peptide derived from the 5HT_3A_ receptor, and the SNAP tag (a large extracellular folded protein domain) ensured trafficking of the engineered receptor to the cell surface (see Supplementary Figure 1).

A full pharmacological comparison of the truncated and full-length CB1R was carried out to understand the effects of CB1R NTD truncation (i.e., the removal of the first 90 amino acids, CB1R_91-472_). Radioligand binding studies with the tritiated synthetic cannabinoid CP 55,940 strongly suggest that CB1R truncation had no discernable effects on synthetic cannabinoid binding (see Supplementary Figure 2 and Supplementary Table 1). In addition, the functional effects of the synthetic cannabinoid, HU-210 and the endocannabinoids, 2-AG and AEA were indistinguishable between truncated and native receptors when tested in intact cells expressing a Gi-CASE activation biosensor (Schihada, Shekhani et al. 2021, Scott-Dennis, Rafani et al. 2023). This suggests that neither the efficacy (intrinsic activity) nor potency of the synthetic cannabinoids nor endocannabinoids was affected by truncation of the CB1R (see Supplementary Figure 3 and Supplementary Table 2).

### Previously developed fluorescent tracers are too slow for measurements of ligand binding kinetics

The binding kinetic profile (the association rate *k*_on_ and the dissociation rate *k*_off_) of the tracer molecule will significantly influence the performance of the Motulsky & Mahan model (Georgi, Dubrovskiy et al. 2019, Sykes, Jain et al. 2019). A previously developed tracer based on HU-308, MKA-115, exhibited a slow rate of association to both CB1R and CB2R. Since fast association is desired to observe competition with the cold compounds at the very earliest time points, MKA-115 was not suitable for use (see Figure 2). In addition, MKA-115 failed to reach equilibrium in the 15 min data collection period meaning that reliable kinetic parameters could not be obtained for this tracer. In total, **10** tracers were designed, synthesized, and tested (see Supplementary Figure 4 and 5 and Supplementary Table 3). From this set only one, **D77**, showed suitable kinetic profiles for our desired application of a universal tracer (see Supplementary Figure 6).

**Figure 2.**
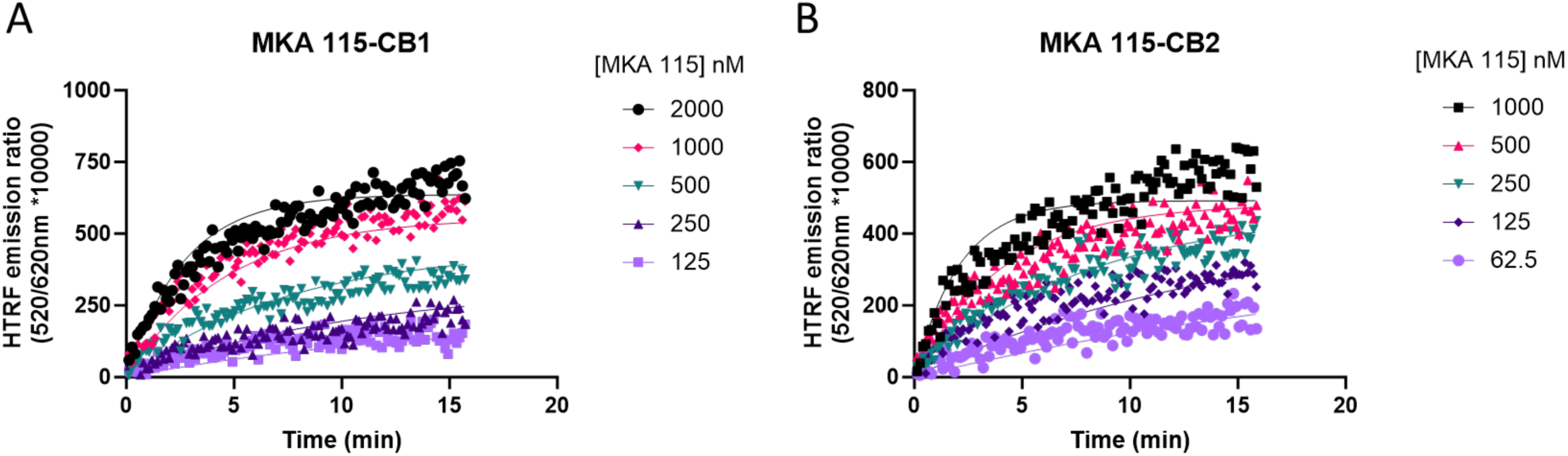
Example kinetic profile of a non-ideal nonselective cannabinoid probe. Kinetic association binding curves of the fluorescent cannabinoid ligand MKA 115 binding to **(A)** CB1R and **(B)** CB2R expressing membranes. Graphs are representatives of 3 independent experiments and show specific binding expressed as HTRF emission ratio.

### Synthesis of Tracer D77 (NBD-Δ^8^-THC)

Our goal of designing a universal fluorescent probe to determine the pharmacology of CBR specific ligands necessitated the identification of a pharmacophore with balanced affinity for both CB1R and CB2R. In addition, the determination of kinetic parameters of lower affinity CBR specific ligands, such as the endocannabinoids, necessitates the use of a rapidly dissociating tracer. These criteria contrast our previously reported, highly CB2R-selective fluorescent probes (Sarott, Westphal et al. 2020). Δ^8^-THC fulfills this criterion and exhibits good affinity for both CB1R and CB2R, providing an ideal starting point for our studies (Soethoudt, Grether et al. 2017). Phenol modification is well-known to sharply reduce the affinity of THC derivatives for CB1R and was thus unsuitable for linker attachment for our purposes (Compton, Prescott et al. 2002). In contrast, chemical probes based on the THC scaffold, as well as available SAR data indicate THC carbon 11 (C(11)) as a suitable locus for attachment of linkers extending into the extracellular matrix, and providing a sound basis for the incorporation of fluorophores (Thakur, Duclos et al. 2005, Papahatjis, Nahmias et al. 2006, Muppidi, Lee et al. 2018). We have previously shown the requirement for minimally six carbon atoms to reach the extracellular space as well as the pharmacological superiority of C(11) amide-over ester-linkages in cannabinoid-fluorophore conjugates; both considerations being accounted for in our design of the FRET tracer D77 (or compound 1) (Westphal, Sarott et al. 2020). Additionally, D77 harbors a C(3) n-heptyl chain, which recently has been shown to enhance affinity for both CB1 and CB2R when compared to the n-pentyl side chain in THC (Citti, Linciano et al. 2019). As a fluorescent moiety we have chosen nitrobenzoxadiazole (NBD). Although NBD has lower extinction coefficient (ca 20000 cm^−1^M^−1^) and quantum yield (ca 0.4) (Uchiyama, Takehira et al. 2003) compared to dyes like fluorescein (65000 cm^−1^M^−1^, 0.98), it has proven to be a good acceptor for the terbium cryptate Lumi4 donor in our hands (Kosar, Sykes et al. 2023).

The modular synthetic strategy and structure of our universal cannabinoid receptor probe D77 is shown in Figure 3 with synthetic methodology described in Supplementary Scheme S1.Once D77 was identified as a suitable tracer, a novel convergent synthetic approach was designed and optimized, starting from commercially available spherophorol, to access D77 **(1)** in 3 steps with 34% overall yield (Supplementary Scheme S2). This dramatically simplified the synthesis of D77 (3 vs 14 steps) and increased the yield of the synthesis from (<1% to 34%). The new streamlined and very efficient synthesis pathway enables ready supply of D77 tracer for screening purposes.

**Figure 3.**
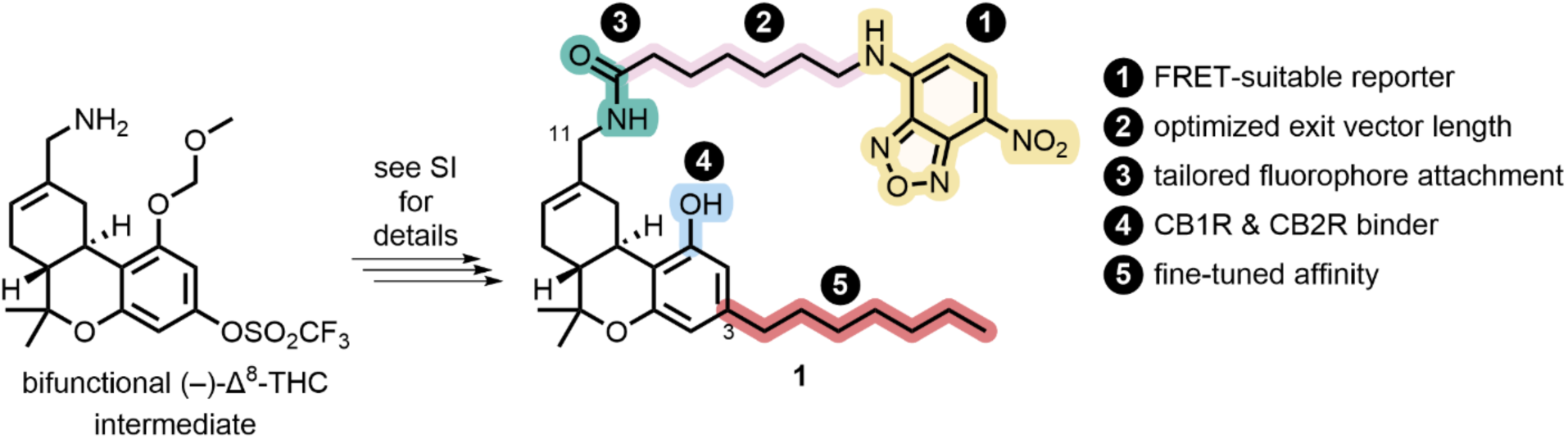
Synthesis and design of D77, a non-selective fluorescent ligand targeting CB1R and CB2R.

### Characterization of saturation binding of D77 to CB1R and CB2R

Saturation binding experiments with increasing concentration of D77 were carried out in order to determine the affinity of D77 for the two cannabinoid receptor subtypes. Measurements at steady-state taken after 5 min following ligand addition were used as an equilibrium endpoint to obtain an equilibrium affinity measurement, or *K*_D_ value at each receptor (see Figure 4). Notably, the level of non-specific binding signal was low at both receptors, representing less than 25% of the total binding signal in both cases. Consequently, D77 achieves a relatively high level of specific binding making it a useful tracer for competition studies and a practical alternative to the high-affinity, agonist radioligands used in the past, and which are routinely employed in the absence of guanine nucleotides to preserve higher levels of specific binding. Equilibrium binding affinity values for D77 binding to the two CBR subtypes are reported in Table 1.

**Figure 4.**
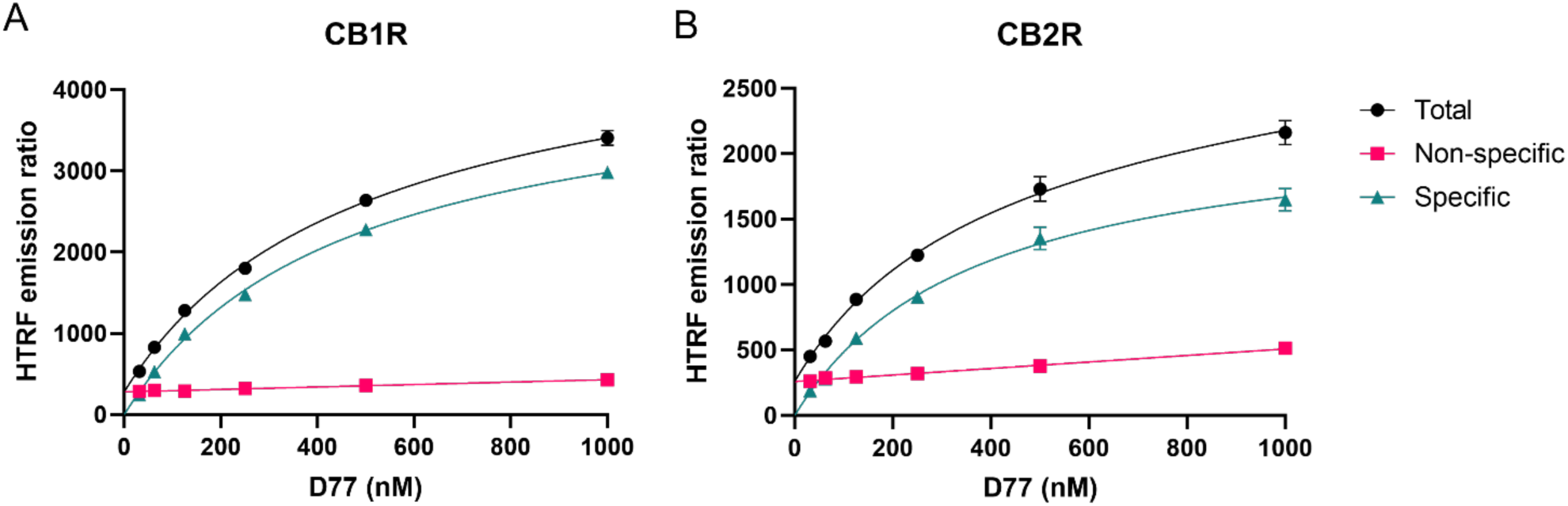
Saturation binding curves at 5 min for D77 ligand binding to CB1R and CB2R. Total binding, non-specific binding, and the specific signal saturation curve are shown for D77 ligand binding to CB1R (A) and CB2R (B) expressed as the HTRF emission ratio. Non-specific binding was defined in presence of 3 μM of rimonabant and 1 μM of SR144,528 for CB1R and CB2R, respectively. Graphs are representatives of 3 independent experiments performed in duplicate and are displayed as mean ± SEM.

**Table 1.** Kinetic parameters *k*_on_ and *k*_off_ and affinity values of D77 fluorescent tracer at 37°C Values are calculated from kinetic association experiments and from saturation experiments performed at equilibrium. Data shown are mean ± SEM of 4 experiments conducted independently.

| | $k_{on}$ ($M^{-1} \text{ min}^{-1}$) | $k_{off}$ ($\text{min}^{-1}$) |
| --- | --- | --- |
| **CB1R** | $(4.3 \pm 0.2) \times 10^6$ | $1.9 \pm 0.1$ |
| **CB2R** | $(3.5 \pm 0.2) \times 10^6$ | $1.1 \pm 0.1$ |
| | Kinetic $K_d$ (nM) | Equilibrium $K_d$ (nM) |
| **CB1R** | $437 \pm 22$ | $427 \pm 19$ |
| **CB2R** | $370 \pm 16$ | $380 \pm 18$ |

### Equilibrium competition experiments using D77

Equilibrium competition experiments allowed the calculation of the equilibrium dissociation constant (or p*K*_i_ values) of our test set of cannabinoid compounds under equilibrium conditions. To achieve this aim binding of D77 to CB1R and CB2R was monitored using TR-FRET in the presence of increasing concentrations of cannabinoid ligands and IC_50_ parameters were obtained from the derived curves, as shown in Figure 5. The Cheng-Prusoff conversion equation was used to calculate *K*_i_ values from the IC_50_ values derived from these inhibitory curves and these values expressed as negative logarithms can be found in Table 2. The p*K*_i_ values obtained using D77 were in good agreement with the literature (Govaerts, Hermans et al. 2004, Khajehali, Malone et al. 2015, Martella, Sijben et al. 2017, Soethoudt, Grether et al. 2017).

**Figure 5:**
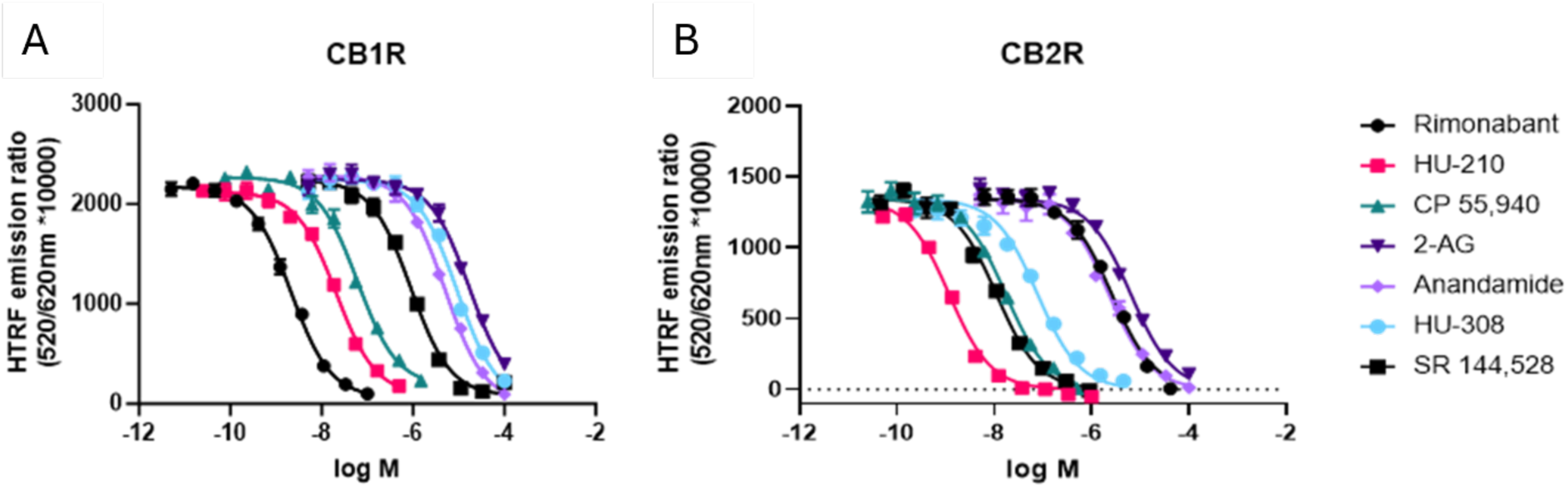
Steady-state competition binding curves of cannabinoid ligands competing with the tracer D77 for CB1R and CB2R. Experiments were conducted using a fixed concentration of the tracer molecule D77 (600 nM) and increasing concentrations of unlabeled ligands using membranes expressing (A) CB1R and (B) CB2R. Competition binding curves for the cannabinoid compounds are shown for both receptors at 37°C and following a 15 min incubation. Graphs are representatives of 3 independent experiments and represent mean ± SEM of 6 technical replicates.

**Table 2.** Equilibrium dissociation constant of the cannabinoid ligands tested expressed as p*K*_i_ at 37°C. The data shown are mean ± SEM from 4 experiments conducted independently in singlet.

| | $pK_i$ | |
| --- | --- | --- |
|  | CB1R | CB2R |
| Rimonabant | 9.0 ± 0.1 | 5.9 ± 0.1 |
| HU-210 | 8.0 ± 0.0 | 9.5 ± 0.1 |
| CP 55,940 | 7.5 ± 0.0 | 8.2 ± 0.1 |
| 2-AG | 5.1 ± 0.3 | 5.7 ± 0.2 |
| Anandamide | 5.6 ± 0.0 | 6.0 ± 0.1 |
| HU-308 | 5.4 ± 0.0 | 7.5 ± 0.1 |
| SR 144528 | 6.4 ± 0.0 | 8.5 ± 0.1 |

### Kinetics of Association of D77 binding CB1R and CB2R

We measured the real-time association of the fluorescent tracer D77 to both cannabinoid receptors subtypes at 37°C over a period of 5 minutes using TR-FRET. D77 showed a rapid association profile to both CB1R and CB2R, reaching equilibrium within the first two minutes (see Figure 6).

**Figure 6.**
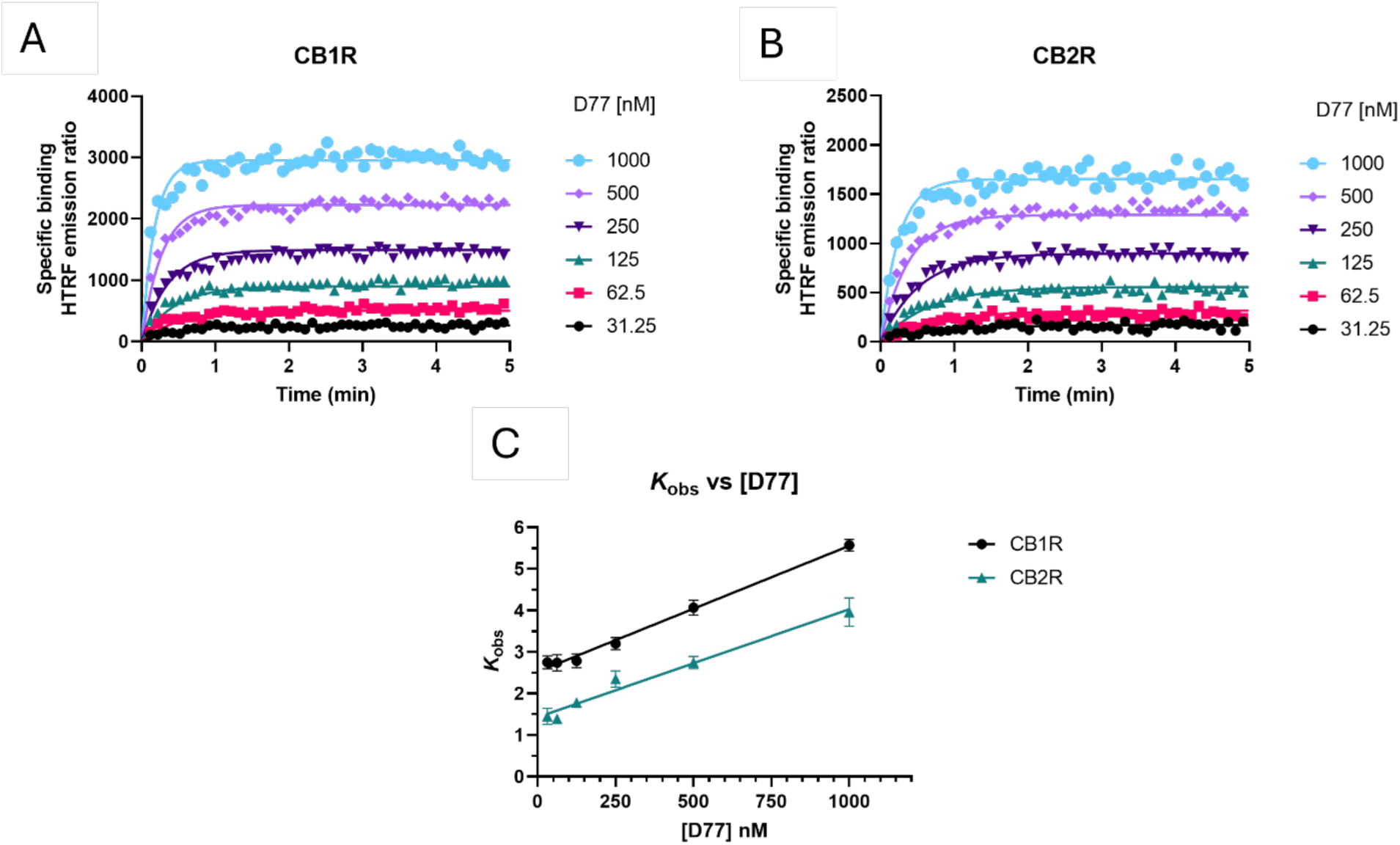
Determination of D77 kinetic binding parameters. Association binding curves obtained from six concentrations of the fluorescent ligand D77 are shown for (A) CB1R and (B) CB2R. Kinetic experiments derive a *K*_d_ for D77 ligand of 437 ± 22 nM for CB1R and 370 ± 16 nM for CB2R. Observed association rate constant (*k*_obs_) obtained for each concentration of fluorescent tracer D77 fitted to a linear regression model (C). Binding followed a simple law of mass action model, k_obs_ increasing in a linear manner with fluorescent ligand concentration. Data presented as representative graphs (A and B) or mean ± SEM (C) from 4 experiments conducted independently.

As shown in Figure 6A and B D77 shows optimal binding characteristics at CB1R and CB2R, making it an ideal tracer for the Motulsky and Mahan approach used to determine the kinetics of unlabeled compounds binding these receptors. D77 shows a relatively fast observed association profile (*k*_obs_ = (*k*_on_ × L) + *k*_off_), yet still allows the accurate determination of the associating phase prior to equilibrium, necessary for the application of the competitive association binding model (see Figure 6A and B). The association rate constants measured for D77 were relatively slow at both receptors (*k*_on_-CB1= (4.3 ± 0.2)×10^6^ M^−1^ min ^−1^, *k*_on_-CB2 = (3.5 ± 0.2)×10^6^ M^−1^ min^−1^). However, D77 exhibits fast dissociation rate constants at both cannabinoid receptors, with *k*_off_ values of 1.87 ± 0.05 min^−1^ and 1.13 ± 0.06 min ^−1^ being obtained at CB1 and CB2R respectively, meaning equilibrium between receptor and tracer is achieved rapidly at both receptor subtypes.

Moreover, the specific binding signal of D77 remains constant over time and we do not observe any decay in the signal due to signal bleaching within the time frame of the data acquisition. The affinity values obtained from saturation equilibrium experiments were in good agreement with the kinetic *K*_D_ values determined using the kinetic approach (where, *K*_D_ = *k*_off_/*k*_on_) (Table 1), demonstrating the reliability of the tracer kinetic model fitting.

As shown in Figure 6C, the observed associated rate (*k*_obs_) increases linearly with fluorescent ligand concentration, demonstrating the expected relationship for a reversible bimolecular binding interaction (Motulsky and Christopoulos 2004). The extrapolation of the fitted line to Y=0 yields to an estimation of *k*_off_ of 2.53 min^−1^ and 1.43 min^−1^ at CB1R and CB2R, respectively, and the slope indicates a *k*_on_ at CB1R of 3.03 x 10^6^ M^−1^ min ^−1^ and at CB2R of 2.60 x 10^6^ M^−1^ min ^−1^. These values are in good agreement with those obtained from global fitting of the D77 association binding curves (see Table 1).

### Competition association binding

Having characterized the kinetics of D77 binding to CB1R and CB2R, we set out to evaluate its use as a tracer for the determination of the kinetics of unlabelled cannabinoid receptor ligands. Competition between the unlabelled compounds and D77 resulted in a concentration-dependent inhibition of the D77 tracer binding in both the case of CB1R (see Figure 7) and CB2R (see Figure 8). The shape of the HU-210 competition curves exhibits the characteristic “overshoot” of the tracer, which can be seen to rapidly bind the receptor before equilibrium with the unlabelled compound is reached. This reveals the slow dissociation profile of the competitor compound, HU-210 relative to the tracer. This effect was more prominent in the case of CB2R due to HU-210’s much slower rate of dissociation at this receptor subtype relative to the tracer (see Figure 7B and 8B). The majority of the competition association curves show a gradual increase in D77 binding, indicating faster dissociation of the competing compounds, apart from SR 144528 binding the CB2R. In Tables 3 and 4 we report the kinetic association and dissociation rate constants (*k*_on_ and *k*_off_) and residence times (Rt = 1/*k*_off_) of the unlabelled agonists and antagonists binding to both CB1R and CB2R.

**Figure 7.**
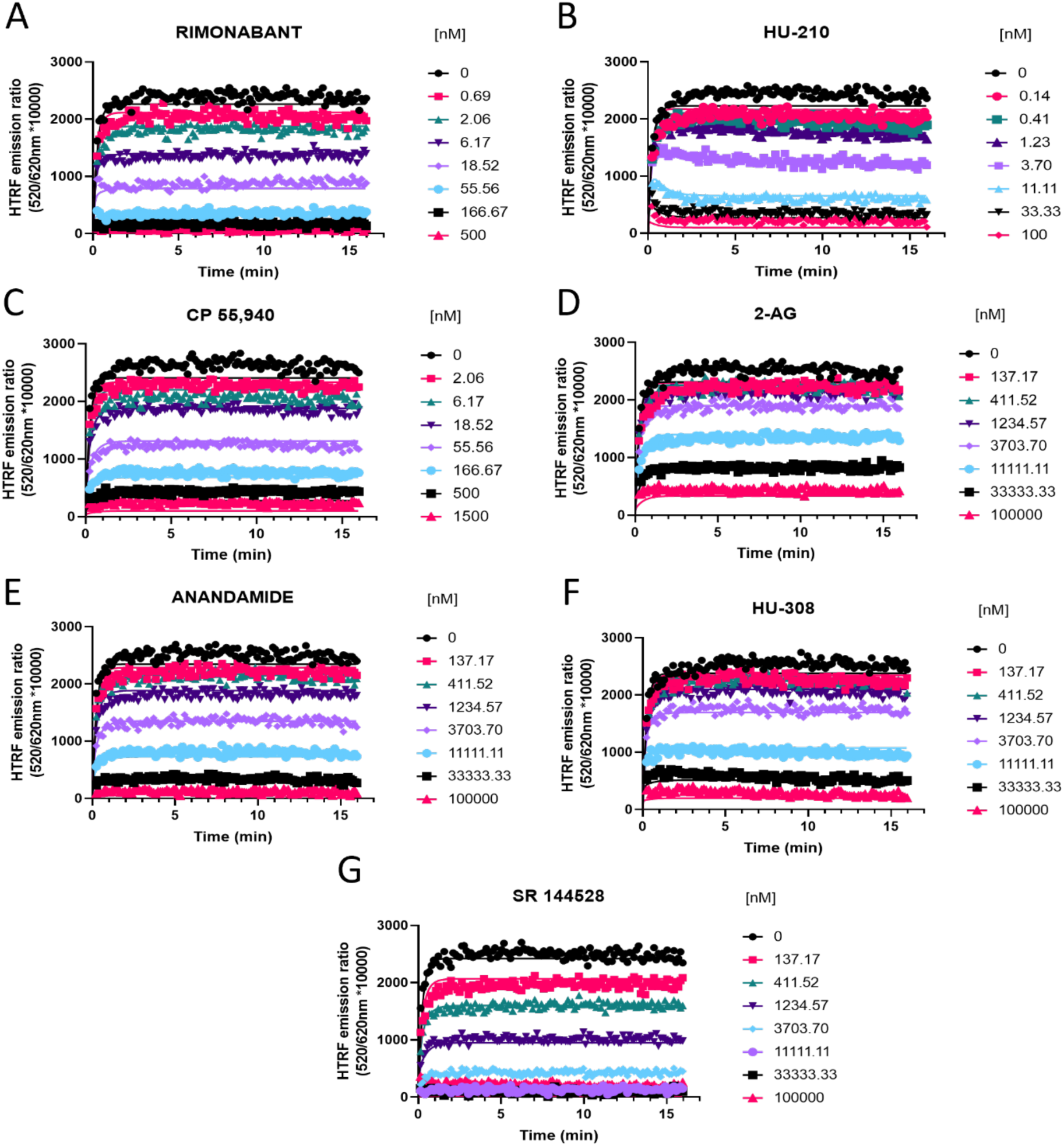
Competition kinetic binding curves of cannabinoid ligands competing with D77 for the at CB1R. Experiments were conducted using a fixed concentration of the tracer molecule D77 (600 nM) and increasing concentrations of unlabeled ligands (A) Rimonabant, (B) HU-210, (C) CP 55,940, (D) 2-AG, (E) anandamide (AEA), (F) HU-308 and (G) SR 144528. Data were globally fitted to the competition association model using GraphPad Prism 9.2 to simultaneously calculate *k*_on_ and *k*_off_ of the unlabeled competitors. Graphs show competition association curves from a single experiment representative of ≥ 3 experiments conducted independently.

**Figure 8.**
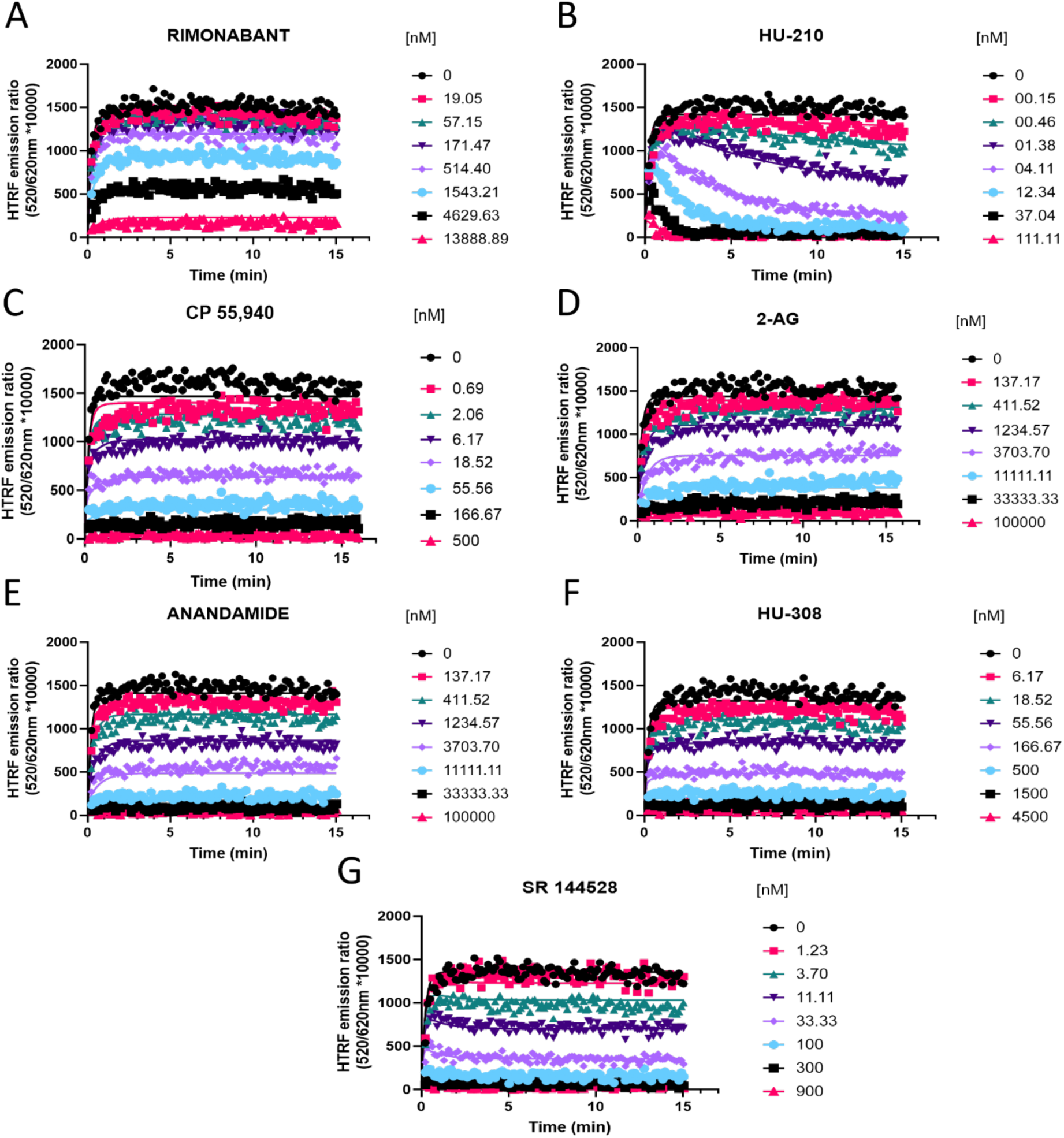
Competition kinetic binding curves of cannabinoid ligands competing with D77 for the CB2R. Experiments were conducted using a fixed concentration of the tracer molecule D-77 (900 nM) and increasing concentrations of unlabeled ligands (A) Rimonabant, (B) HU-210, (C) CP 55,940, (D) 2-AG, (E) Anandamide (AEA), (F) HU-308 and (G) SR 144528. Data were globally fitted to the competition association model using GraphPad Prism 9.2 to simultaneously calculate *k*_on_ and *k*_off_. of the unlabeled competitors. Graphs show representative curves from 4 experiments conducted independently.

**Table 3.**
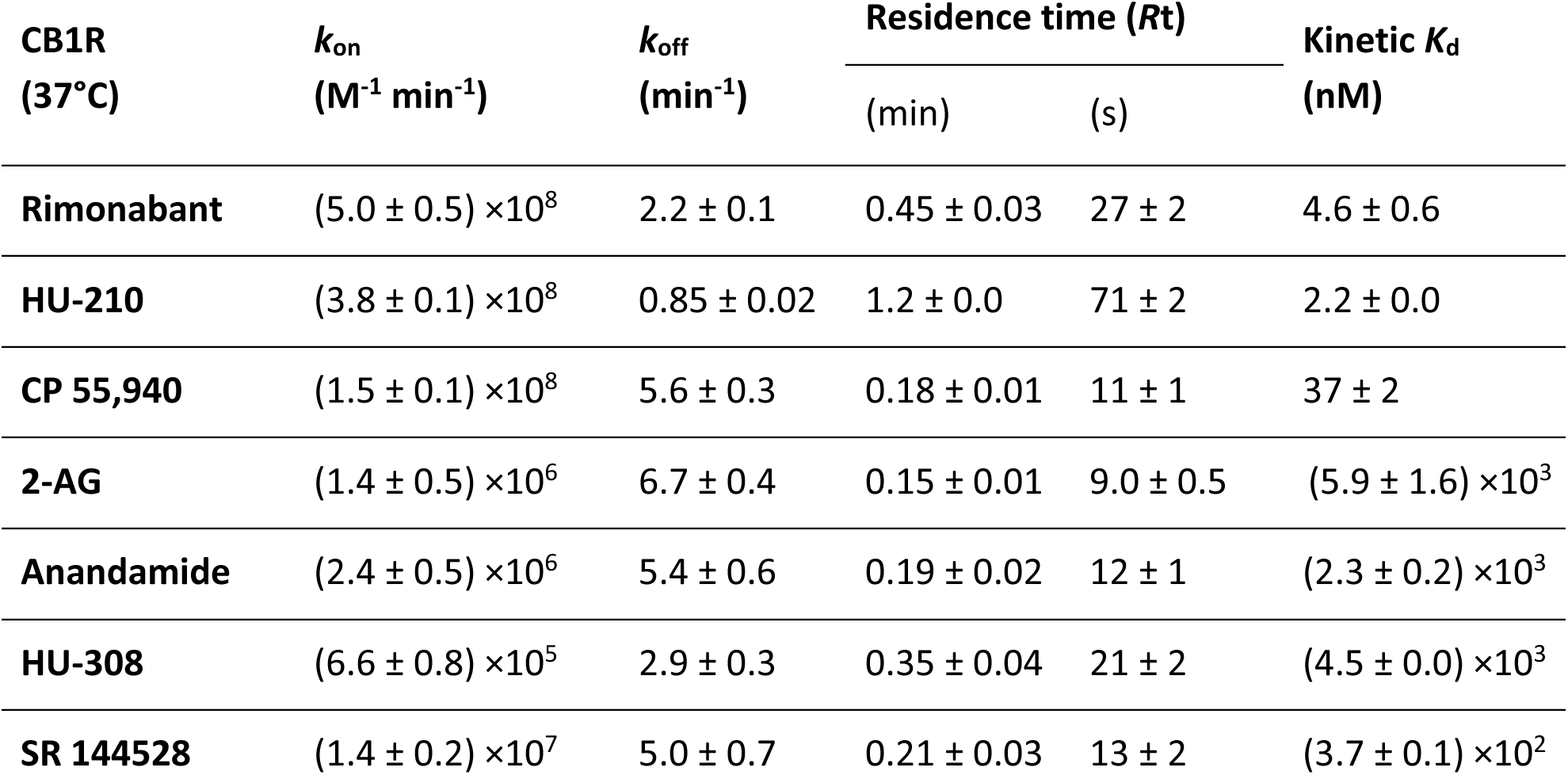
Kinetic parameters calculated from the Motulsky and Mahan experimental approach for the cannabinoid compounds tested at CB1R at 37°C. The data shown are mean ± SEM from 4 experiments conducted independently (except for 2-AG, where N=3)

**Table 4.** Kinetic parameters calculated from the Motulsky and Mahan experimental approach for the cannabinoid compounds tested for CB2R at 37°C. Data are expressed as mean ± SEM of 4 experiments conducted independently.

| CB2R<br>(37°C) | $k_{on}$<br>($M^{-1} min^{-1}$) | $k_{off}$<br>($min^{-1}$) | Residence time ($Rt$) | | $K_d$<br>(nM) |
| --- | --- | --- | --- | --- | --- |
|  |  |  | (min) | (s) |  |
| Rimonabant | $(9.2 \pm 1.2) \times 10^6$ | $6.4 \pm 0.7$ | $0.16 \pm 0.02$ | $10 \pm 1$ | $701 \pm 52$ |
| HU-210 | $(2.6 \pm 0.2) \times 10^8$ | $0.051 \pm 0.002$ | $20 \pm 1$ | $1190 \pm 67$ | $0.20 \pm 0.02$ |
| CP 55,940 | $(4.4 \pm 0.5) \times 10^8$ | $1.9 \pm 0.1$ | $0.55 \pm 0.04$ | $33 \pm 3$ | $4.27 \pm 0.24$ |
| 2-AG | $(1.2 \pm 0.2) \times 10^7$ | $14 \pm 2$ | $(7.4 \pm 1.2) \times 10^{-2}$ | $4.4 \pm 0.7$ | $1213 \pm 109$ |
| Anandamide | $(9.7 \pm 0.4) \times 10^6$ | $6.1 \pm 0.7$ | $0.17 \pm 0.02$ | $10 \pm 1$ | $632 \pm 52$ |
| HU-308 | $(5.6 \pm 0.3) \times 10^7$ | $1.5 \pm 0.1$ | $0.66 \pm 0.04$ | $40 \pm 2$ | $27.5 \pm 1.5$ |
| SR 144528 | $(2.5 \pm 0.2) \times 10^8$ | $0.69 \pm 0.03$ | $1.46 \pm 0.07$ | $87 \pm 4$ | $2.82 \pm 0.16$ |

For CB1R, the faster associating compounds were rimonabant, HU-210 and CP 55,940, displaying *k*_on_ values of 5, 3.8, and 1.5×10^8^ M^−1^ min^−1^, respectively. The two endocannabinoid compounds 2-AG and AEA showed similar slower association rate constants (*k*_on_ = 1.4 and 2.4×10^6^ M^−1^ min^−1^, respectively). The slowest associating compound was a CB2R selective binder, HU-308, with a *k*_on_ of 6.6×10^5^ M^−1^ min^−1^, consistent with its lower affinity for CB1R.

Regarding the dissociation profile of the compounds at CB1R, the faster dissociating compounds were CP 55,940, 2-AG, AEA and SR 144528, therefore displaying the shorter residence times at this receptor subtype of ∼10 s. HU-210 exhibited the longest residence time (slowest *k*_off_) of 71 s, followed by the inverse agonist rimonabant (Rt= 27 s) and HU-308 (Rt= 21 s).

The competition binding approach has revealed a remarkably small difference between the kinetic dissociation parameters of these ligands with only the agonist HU-210 displaying a relatively slow off-rate from the CB1R compared to the rapidly dissociating tracer D77 and the endogenous agonists AEA and 2-AG.

For CB2R, the faster associating compounds were HU-210, CP 55,940 and SR 144528, displaying *k*_on_ values of 2.6, 4.4, and 2.5×10^8^ M^−1^ min^−1^, respectively, followed by HU-308, which displayed a *k*_on_ of 5.6×10^7^ M^−1^ min^−1^. The two endocannabinoid compounds 2-AG and AEA showed similar slower association rate constants (*k*_on_ 1.2 x10^7^ and 9.7×10^6^ M^−1^ min^−1^ respectively), similar to rimonabant (*k*_on_ of 9.2×10^6^ M^−1^ min^−1^).

Regarding the dissociation profile of the compounds, the fastest dissociating compounds were 2-AG, rimonabant and AEA, therefore displaying the shorter CB2R residence times of ∼4 s (2-AG) and ∼10s (both rimonabant and AEA). CP 55,940, HU-308 and SR 144528 exhibited slower dissociation rates, with residence times of 33 s, 40 s and 87 s. By far the slowest dissociating compounds was HU-210, with a residence time of >20 min.

### Kinetic parameters correlate differently for CB1R and CB2R

We performed a correlation analysis using the kinetic parameters (*k*_on_ and *k*_off_) versus affinity values obtained for our test set of compounds, in order to explore the role of association and dissociation constant rates in dictating the affinity of the ligands for CB1R and CB2R.

The affinities and association rates of compounds show a strong correlation at both CB1R and CB2R when performing a correlation analysis of their logarithmic transformations (p*K*_D_ vs log *k*_on_), whereas the dissociation rates, expressed as negative logarithmic transformations, are only significantly correlated with ligand affinity values for CB2R (see Figure 9). These results suggest that the *k*_on_, the association rate constant rather than *k*_off_, the dissociation rate constant is the biggest determinant of receptor affinity for CB1R, whereas both *k*_on_ and *k*_off_ parameters dictate the affinity for the CB2R binding. The correlation between the affinity and dissociation rates for the CB2R, found with the selected compounds in this study, contrasts with the results published previously with a different set of CB2R binding compounds (Martella, Sijben et al. 2017) where no significant correlation was reported between affinity and *k*_off_ values.

**Figure 9.**
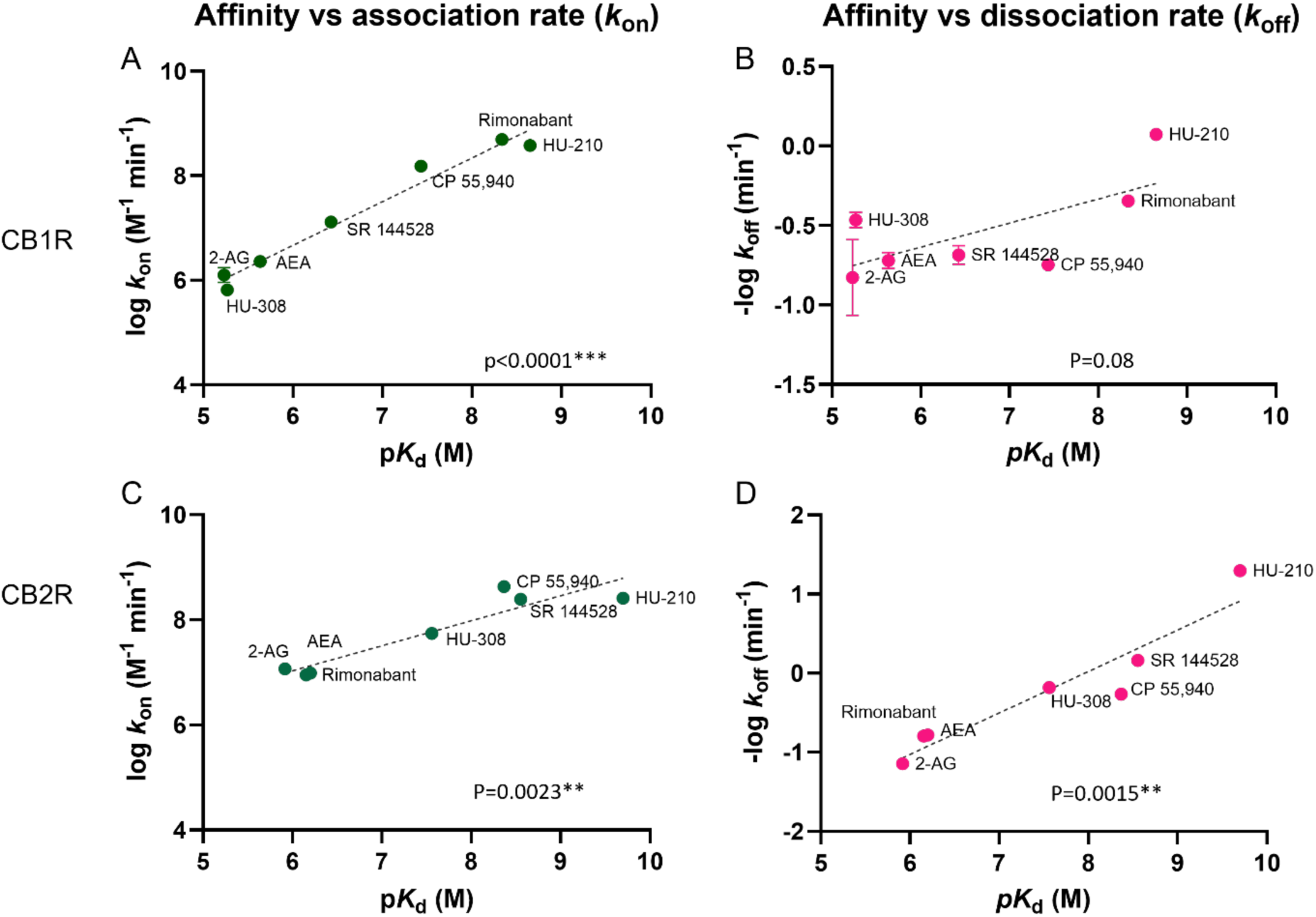
Correlation plots of equilibrium and kinetic parameters of cannabinoid compounds at CB1R and CB2R. Correlation between negative logarithmic transformation of affinities (-log *K*_d_) and logarithmic (A) association rate (log *k*_on_) and (B) dissociation rate (log *k*_off_) for CB1R ligands. Correlation between negative logarithmic transformation of affinities (-log *K*_d_) and logarithmic (C) association rates (log *k*_on_) and (D) dissociation rates (log *k*_off_) for CB2R ligands. Correlation analysis was carried out using a Pearson correlation analysis (two-tails). Data shown are the mean and SEM of 4 independent experiments.

The equilibrium dissociation constants for the tested cannabinoid compounds were calculated from the kinetic association and dissociation rates from kinetic experiments (kinetic *K*_D_; *K*_D_=*k*_off_/*k*_on_) and from the equilibrium displacement data (*K*_i_). Both values were compared for the binding of the compounds to both CB1R and CB2R. As shown in Figure 10, the kinetic *K*_D_ affinity values generated showed a strong correlation with the *K*_i_ values obtained from the equilibrium displacement binding, a Pearson correlation coefficient (r) of 0.97 (P=0.0004) and 0.98 (P<0.0001) was obtained at CB1R and CB2R respectively. These results show that the association and dissociation rates obtained from the association competition experiments are consistent with the affinity values obtained from equilibrium competition data, validating our approach using the tracer D77.

**Figure 10.**
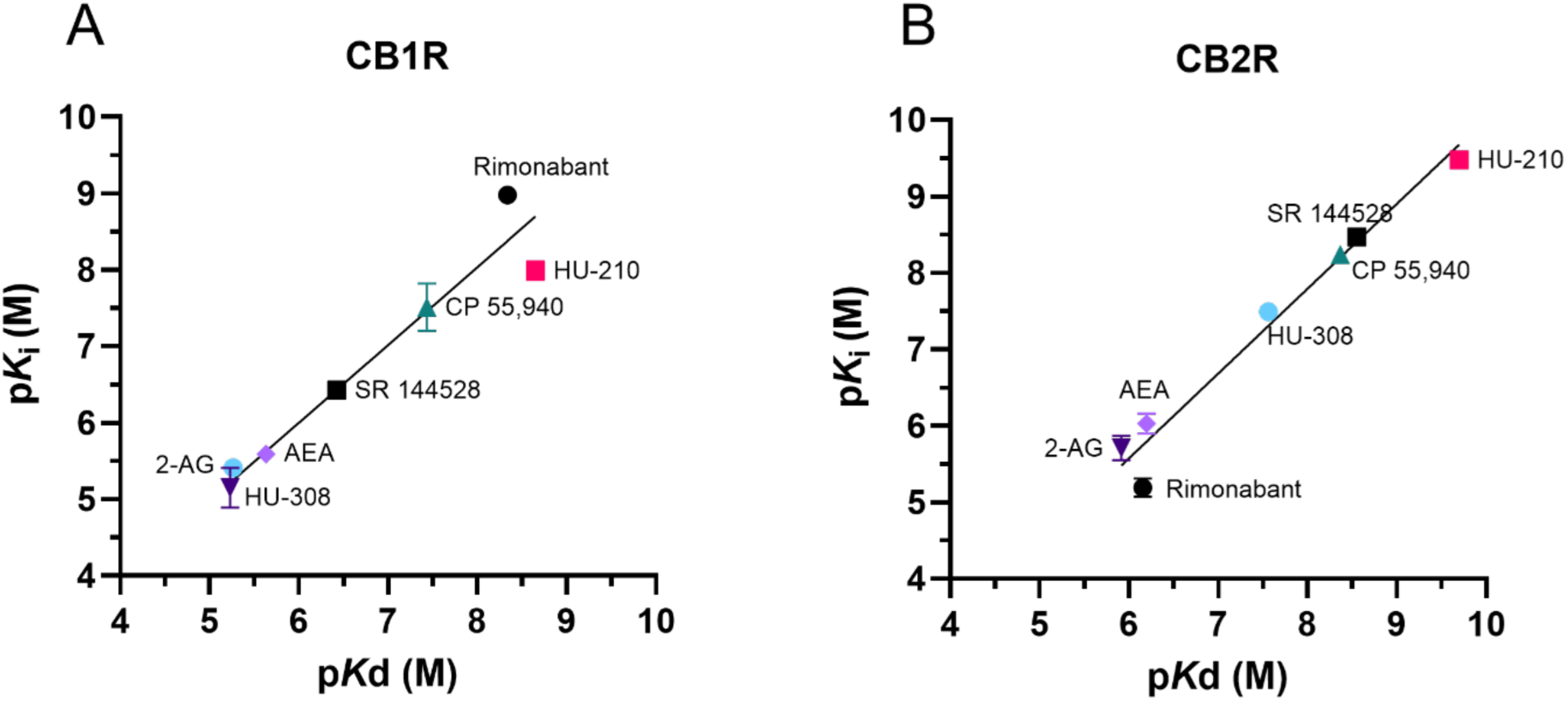
Kinetic versus equilibrium affinity estimates. Correlation between affinity values obtained from equilibrium displacement (p*K*_i_) and kinetic binding experiments (p*K*_D_) for the seven test compounds at **(A)** CB1R and **(B)** CB2R. p*K*_i_ values were taken from D77 competition binding experiments at equilibrium (see Figure 6). The values comprising the kinetically derived *K*_D_ (*k*_off_/*k*_on_) were taken from the experiments shown in Figure 7 and Figure 8.

## Discussion

In this study, we set out to develop a fluorescent ligand binding assay for profiling the kinetic parameters of compounds binding to the orthosteric binding site of CB1R and CB2R. The motivation for this was to advance on the existing radioligand binding techniques currently available for this purpose. An important consideration for such filtration-based radioligand binding assays is that the tracer itself must possess a relatively slow-off rate from the receptor of interest, so that the effective separation of bound and unbound radioligand is achievable during the washing stage. However, in order to quantify the kinetic paraments of typical CB2R selective ligands, that display fast dissociation kinetics, the tracer itself must also display comparably fast dissociation kinetics (Sykes, Jain et al. 2019). Fluorescence-based TR-FRET assays are ideal for these purposes as they are homogenous, and do not require the separation of bound and unbound ligand.

### Development of TR-FRET ligand binding assay for cannabinoid receptors

Previously, we reported a TR-FRET ligand binding assay for CB2R (Sarott, Westphal et al. 2020, Gazzi, Brennecke et al. 2022). Here, we developed the same assay format for CB1R. Firstly, we used genetic engineering to introduce a SNAP-tag, enabling the incorporation of the donor, terbium cryptate, so that our assay reported binding to only the overexpressed subtype of the receptor and unaffected by tracer specificity (unlike in the radioligand binding assay). TR-FRET is a well-established approach, but it is often underappreciated that it provides ultimate assay specificity irrespective of the selectivity of the tracer employed. Secondly, TR-FRET only reports a signal based on proximity. This improves the specific/non-specific signal ratio compared to traditional radio-ligand binding assays, where all ligand remaining after filtration of the membranes contributes to the total signal, including the ligand which remains bound to unspecific sites of the cell membrane, or the filters themselves.

However, for CB1R the truncation of the N-terminus was necessary to shorten the donor-acceptor distance and to facilitate the FRET process. While there is always a consideration that any receptor modification may affect its biogenesis, ligand binding or signaling properties, our imaging, binding, and functional data strongly suggest that the distal N-terminal residues are not in any way impacting the trafficking or binding/activation capability of the receptor (see Supplementary Figure 1, 2 and 3). This same truncation strategy could be applied to develop fluorescent ligand binding assays for other receptors which have an exceptionally long N-terminus.

We recently reported the development of high affinity selective fluorescent probes for CRB1R (Mach, Omran et al. 2024) where we used truncated CB1R for ligand binding characterization. Fluorescent ligand binding to the human CB1R assay has also been described previously using CELT-335 ligand, TR-FRET and a SNAP-tagged full-length CB1R (Raïch, Rivas-Santisteban et al. 2021, Navarro, Sotelo et al. 2023). However, only equilibrium binding data were reported. It is unclear if kinetic assays with the reported tracer would be feasible, as its binding kinetic parameters were not reported.

### D77 is an excellent fluorescent tracer for equilibrium ligand binding to CB1 and CB2

Our secondary aim was to develop a kinetically fast fluorescent ligand that would serve as an ideal tracer to profile other compounds binding to CB1R and CB2R. In order to achieve this aim, we synthesized and developed D77, a fluorescent version of Δ^8^-THC, as an optimal fluorescent tracer for cannabinoid receptors. Δ^8^-THC is a slightly less potent version of Δ^9^-THC (commonly known as THC). D77 performed extremely well in terms of its ability to accurately measure the affinity of the non-fluorescent competitor compounds. Given its ease of use and our ability to scale up the testing throughput, through use of a commonly available plate reader with injectors and TR-FRET capability, this assay employing D77, effectively replaces radioligand binding assays previously employed to screen for ligands binding to cannabinoid receptors.

### Fluorescent tracer D77 applicability to kinetic studies

Crucially D77, based on Δ^8^-THC, has a balanced affinity for both CBR subtypes, and exhibits a more rapid *k*_off_ over the previously described CB2R tracers (Sarott, Westphal et al. 2020, Gazzi, Brennecke et al. 2022, Kosar, Sykes et al. 2023). This improves the assay performance and allows more accurate and precise estimates of the kinetic parameters of more rapidly dissociating compounds (Georgi, Dubrovskiy et al. 2019, Sykes, Stoddart et al. 2019). The kinetic parameters for a number of compounds including HU-308, CP 55,940, 2-AG, AEA and SR 144528 obtained with D77 are similar to those obtained earlier using a radioligand kinetic binding assay at 25°C (Martella, Sijben et al. 2017). Of note, D77 served our interest of developing an assay capable of determining the kinetic parameters of unlabeled ligands at physiological temperature. Moreover, the use of injectors overcomes the challenge of working at 37°C, which necessitates the collection of early time points, due to the extremely fast association and dissociation rates observed for both the tracer D77 and the tested ligands.

### Measuring kinetic parameters of cannabinoid ligands at physiological temperature creates opportunities for improving compound efficacy at limited diffusion conditions

We successfully determined the kinetic binding parameters of seven different reference compounds binding to CB1R and CB2R. The kinetic parameters determined at 25°C (see Supplementary Tables 4 to 7) and 37°C (see Table 3 and 4) reveal important differences in the residence time of the compounds tested at CB1R and CB2R, where for example, HU-210 displayed a much longer residence time at CB2R (over 30 min longer). These findings strongly support the development of new approaches that enable kinetic characterization of ligand-receptor binding at physiological temperatures, allowing more accurate and detailed preclinical prediction models of drug action to be formulated. For example if a particular receptor is mainly expressed in the brain, prediction of drug concentration in this compartment over time, through implementation of a rebinding model, will be more useful to identify optimal drug dosing (Sykes, Moore et al. 2017). In view of HU-210s rapid association and slow off rate it is intriguing that it has been shown to exhibit a noticeably longer duration of action in preclinical animal studies (Hruba and McMahon 2014) and thus may possess unique beneficial qualities when it comes to behavioral and neurobiological alterations, compared to THC and other cannabinoids (Farinha-Ferreira, Rei et al. 2022). In contrast, the endogenous cannabinoids exhibit slow association and a much more rapid dissociation profile from both receptor subtypes. The potential for rebinding to affect the pharmacodynamic properties of ligands binding to CB1R in the brain and periphery has been documented in previous publications (Vauquelin 2010; Terry et al., 2009) and its effect on apparent receptor reversal of the endogenous agonist 2-AG and the synthetic agonist HU-210, is illustrated in Figure 11. In conditions of limited diffusion as occurs in the synapses of the brain we can expect that HU-210, which possess a relatively rapid association rate, should occupy receptors for much longer than is the case in the periphery where diffusion occurs more freely. In contrast under identical conditions the endogenous ligand 2-AG, which has a much slower association rate and more rapid dissociation rate, would be expected to reverse more rapidly with very little influence of diffusion on this process.

**Figure 11.**
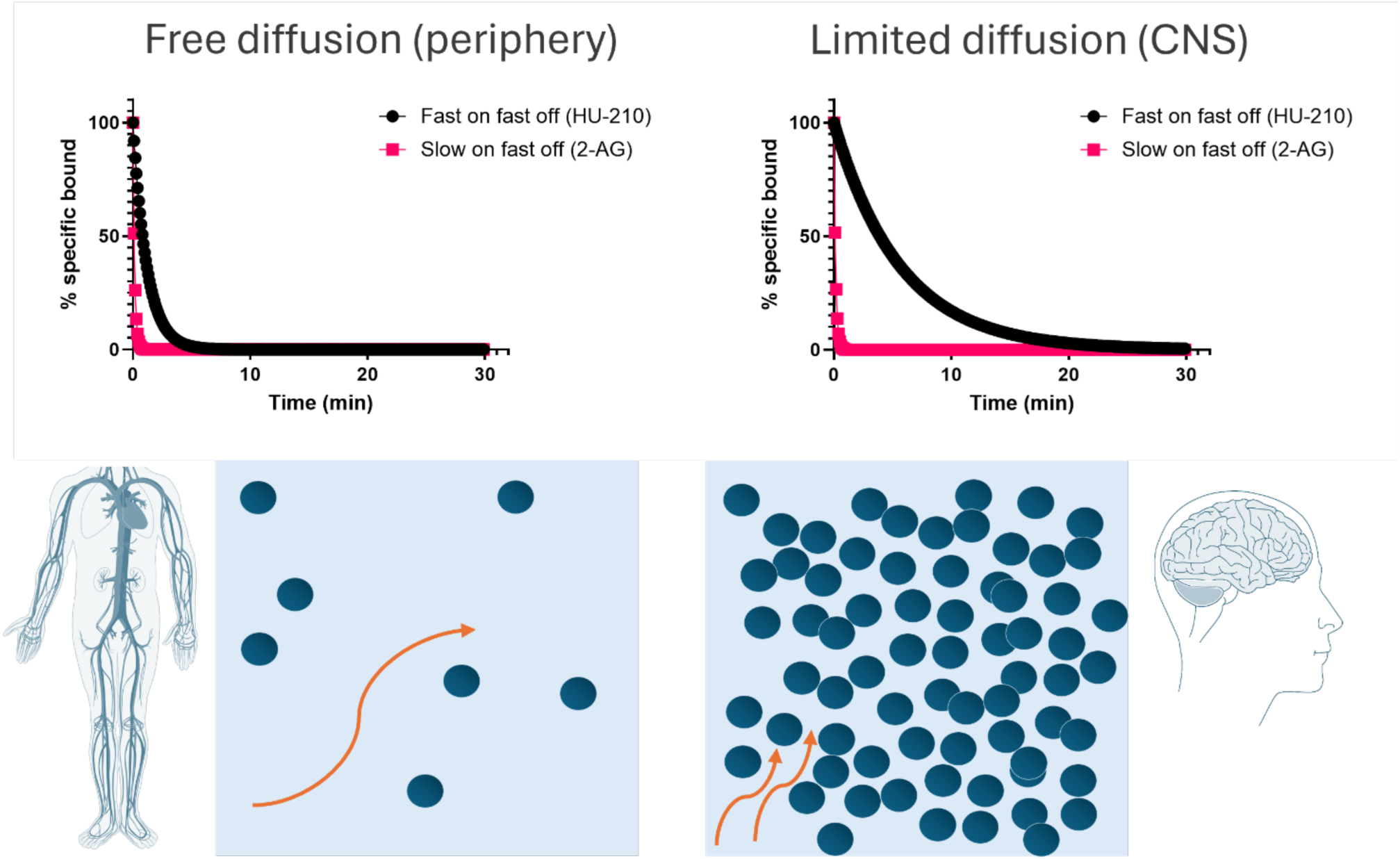
Consequences of rebinding on the apparent reversal of CB1R occupancy in the brain and periphery. Simulated target reversal rates were derived under conditions of limited diffusion based on the association (*k*_on_) and dissociation (*k*_off_) rates determined in competition kinetic binding experiments. All kinetic parameters used to these plots are taken from Table 3. For simulation purposes the reversal rate *k*_r_ was based on the model of an immunological synapse. *k*_r_ values calculated using the following equation *k*_r_ = *k_off_*/(1+ *k_on_* *R/k-) where: *k*_off_ = dissociation rate from the receptor, *k*_on_ = association rate onto the receptor, [R] = surface receptor density fixed at 1×10^11^ cm^−2^ for CB_1_R calculations, and k-= the diffusion rate out of the synaptic compartment into bulk aqueous, fixed at 1.2×10^−5^ cm/sec.

### Future applications

The TR-FRET -based equilibrium and kinetic ligand binding assay developed for CB1R and CB2R can be used to characterize and develop novel compounds with improved sub-type selectivity for these receptors. This novel assay can also be used to better understand the receptor target occupancy of different cannabinoid ligands for these two key subtypes and potentially to screen for allosteric modulators, something which has only been possible with radioligands up until now (Price, Baillie et al. 2005, Bouma, Broekhuis et al. 2023).

### Limitations of the study

Previous TR-FRET binding assays have been successfully developed to profile the kinetics of ligands binding GPCRs (Schiele, Ayaz et al. 2015, Sykes, Moore et al. 2017). One limitation of this new CB1R assay format is that the receptor needs to be truncated to incorporate the SNAP-tag at the N-terminus close to the orthosteric binding site. Indeed fluorescent ligand binding to the human CB1R has been described previously using TR-FRET and a SNAP-tagged full-length CB1R (Raïch, Rivas-Santisteban et al. 2021), however it is unclear if kinetic assays with the reported tracer would be feasible, as its binding kinetic parameters are not reported.

## Conclusions

The novel TR-FRET-based method we outline utilizing the probe D77 (or fluorescent Δ^8^-THC) constitutes a simple and superior alternative to radioligand binding methodologies, to determine equilibrium and kinetic binding of compounds for cannabinoid receptors at physiological temperature. Investigating the kinetic parameters of prospective cannabinoid drug candidates could help us identify essential factors for refining their design and lead to the discovery of more effective medicines to target these receptors.

## Acknowledgements

LBR is a recipient of a Predoctoral grant awarded by the Department of Education of the Basque Government, a Short-Term Fellowship awarded by European Molecular Biology Organization (EMBO) and a Short-Term Scientific Mission grant awarded by European Research Network on Signal Transduction (ERNEST, Cost Action 18133). DAS and DBV gratefully acknowledge funding by F. Hoffmann-La Roche Ltd., Basel, Switzerland [Roche Postdoctoral Fellowship RPF-551]. DBV gratefully acknowledges funding by the Swiss National Science Foundation grants 135754 and 159748 and the Novartis Foundation FreeNovation grant. EJK, DAS and DBV gratefully acknowledge funding by the Medical Research Council [grant number MR/Y003667/1]. JB contribution was supported by the Dutch Research Council (NWO, Vidi #16573). LBR, BLH, JB, MSD, EJK, LHH, DAS, and DBV are members of COST Action CA18133 “ERNEST”.

## Author contributions

LBR and BLH designed and performed experiments, analyzed data, wrote the first drafts of the manuscript, and contributed to the writing of the manuscript. MK, RCS and KJP carried out the design and synthesis of fluorescent compounds and contributed to the writing of the manuscript under the supervision of EMC. TG and LM carried out design and synthesis of fluorescent compounds under the supervision of MN. WG, EK, ACR and UG participated in modelling, design, and characterization of compounds. JB performed and analyzed radioligand binding assays under the supervision of LHH. MSD validate the receptor constructs using Gi-CASE assays. EJK performed microscopy experiments. DBV, DAS, SB and JS conceptualized the project. DAS and DBV designed and supervised the research and contributed to the writing of the manuscript. All authors commented and approved the final version of the manuscript.

