## Supplementary data and methods for "A universal cannabinoid CB1 and CB2 receptor TR-FRET kinetic ligand binding assay"

### Supplementary Methods

#### Confocal microscopy

T-RexTM-293 cells were seeded into 6-well plates containing a poly-D-lysine coated glass coverslip. 24h later cells were transfected with 1 ug/well of pcDNA4/TO plasmids encoding either full length human CB1R or truncated ($CB1R_{91-472}$) receptors containing an N-terminal SNAP tag, using PEI transfection and a 1:6 ratio of DNA:PEI. 24 hours post transfection, cells were washed in PBS and labeled with SNAP-Surface® Alexa Fluor® 647 (100 nM; NEBiolabs) for 30 minutes in growth medium, washed 3 times with PBS to remove unreacted dye, and fixed using 3% paraformaldehyde for 20 min, and washed twice with PBS. Coverslips were then mounted onto glass slides for confocal imaging. Images were recorded using a Zeiss LSM 710 laser scanning confocal microscope fitted with a Zeiss Plan-Apochromat 63x/1.40 NA oil immersion objective.

#### Radioligand binding assays

All radioligand binding assays were carried out at 25°C in a 100 µL reaction volume containing assay buffer (50 mM Tris-HCl (pH 7.4), 5 mM $MgCl_2$, 0.1% (w/v) bovine serum albumin (BSA)) and 1.5 µg membrane protein from T-RexTM-293 cells expressing CB1R or $CB1R_{91-472}$ in the presence of 1.5 nM [$^3$H]CP 55,940 (Specific activity 101 Ci/mmol, Revvity, Waltham, MA, USA). Nonspecific binding was determined using 10 µM SR141716A (rimonabant) and DMSO concentrations were < 1%. In all cases, total radioligand binding (TB) did not exceed 10% of the total radioligand added to avoid ligand depletion.

Displacement assays were carried out on membrane aliquots using radioligand and increasing concentrations of competing ligands CP 55,940 or SR141716A. The reaction mixture was incubated for 120 minutes after which receptor-bound radioactivity was measured. For association assays, T-RexTM-293 CB1R WT or T-RexTM-293 $CB1R_{91-472}$ membranes were added to the reaction mixture at eleven different time points over a total incubation time of 120 minutes. In dissociation assays, membranes were first incubated with radioligand for 120 minutes to reach equilibrium. Dissociation was initiated by addition of 5 µL 10 µM SR141716A (final concentration) at ten different time points. Final DMSO concentrations were increased to 1.5% to improve solubility.

For all assays, incubations were terminated by rapid vacuum filtration over 96-well Whatman GF/C filter plates using a PerkinElmer Filtermate harvester (Revvity, Waltham, MA, USA). Subsequently, filters were washed 20 times using ice-cold wash buffer (50 mM Tris-HCl (pH 7.4), 5 mM MgCl2, 0.1% (w/v) BSA) to reduce non-specific binding. Filter-bound radioactivity was determined in a 2450 Microbeta2 plate counter after addition of 25 µL MicroScint scintillation cocktail per well (Revvity).

All radioligand binding data were analyzed using Prism 9.0 (GraphPad Software, San Diego, California, USA). The $pIC_{50}$ values were obtained from nonlinear regression analysis of competition displacement assays. Apparent association rate constants ($k_{obs}$) were determined by fitting the association data to a one-phase exponential association function. Dissociation rate constants ($k_{off}$) were determined by fitting the data to the statistically preferred biphasic ($k_{off,\ fast}$ and $k_{off,\ slow}$) exponential decay function. Data are shown as mean ± SEM of three individual experiments performed in duplicate. To compare differences in $pIC_{50}$ or kinetic parameters, an unpaired, two-tailed Student's t-test was performed ($p < 0.05$ *).

**Gi-CASE cellular Nano-BRET assay**

Receptor activation of CB1R and truncated $CB1R_{91-472}$-expressing T-Rex$^{TM}$-293 cells with the Gi-CASE biosensor was achieved as follows. Cells were plated in a white, 96-well clear-bottomed previously coated in poly-D-lysine (5 μg/mL in PBS). After 48-72-hour incubation with tetracycline to induce CB1R expression, cell culture media was aspirated and the cells were then washed (100 µl/well) with assay buffer, Hank's balanced salt solution (HBSS) containing 0.5% BSA, 5 mM HEPES. Assay buffer (90 µl/well) containing the NanoLuciferase substrate furimazine (10 µM) was dispensed into the wells and the plate was incubated at 37$^{o}$C for 15 minutes. The assay plate was transferred to the PHERAstar FSX and three BRET cycles were collected every minute, before adding 10 µl of a 10x stock of compound. Dilutions of the synthetic CBR agonist HU-210 and endocannabinoids 2-AG and AEA were prepared as follows. Compounds were initially serially diluted in DMSO, and then diluted 1/10 in assay buffer before the addition to the assay plate. The final DMSO concentration in the cell plate was 1% after compound addition.  Data from compound and vehicle treated cells were collected with 1 min intervals during 40 cycles after compound addition using the BRET1 plus module (535-30LP/475-30BP). Normalized data was fit to a four-parameter log(agonist) vs. response model in GraphPad Prism.

### Supplementary Results

#### Confocal imaging to determine cell surface localisation

Removing part of the CB1R at its N-terminus meant removing amino acid residues which are likely to be glycosylated. As NT glycosylation is implicated in receptor maturation and trafficking for many receptors (Schjoldager, Narimatsu et al. 2020), it was important to check the localisation of the truncated receptor to ensure appropriate expression, folding, and cell surface trafficking. In transiently transfected HEK293T cells, there was clear evidence of cell surface localisation of both the SNAP-tagged full length CB1R and CB1R$_{91-472}$ receptors when the SNAP-tag was labelled with Alexa Fluor 647 and visualised by confocal microscopy (see Supplementary Figure 1).

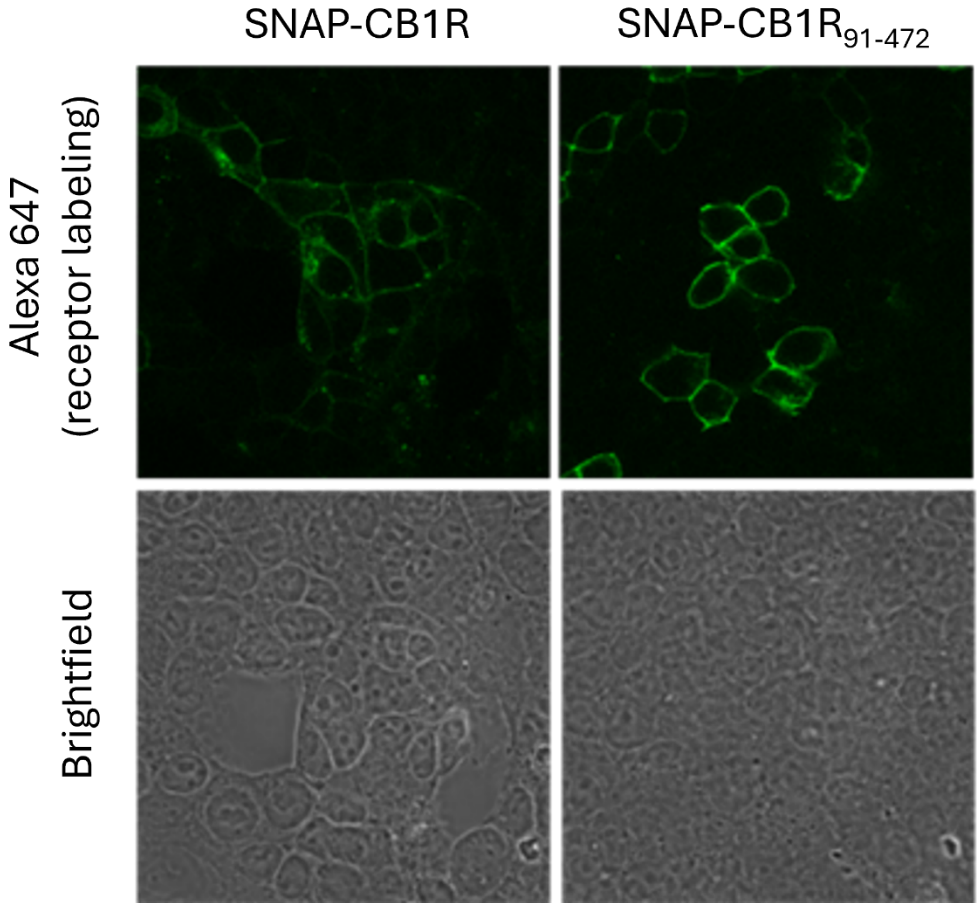


**Supplementary Figure 1.** Cell surface localization of SNAP-tagged full-length CB1R and truncated CB1R$_{91-472}$ receptors by confocal microscopy.

**Detailed binding and functional comparison of the full length CB1R WT and truncated CB1R$_{91\text{-}472}$**

Truncation of CB1R is unlikely to affect the binding of orthosteric ligands which bind deep in the transmembrane helices of the receptor. It has however been suggested that splice variants of CB1R with similarly truncated NT regions might have disrupted the binding or signalling potency of the endogenous agonists such as anandamide (AEA) (Ryberg, Vu et al. 2005). However, a more recent study contradicted these findings, suggesting no effect on the pharmacology of recombinantly expressed CB1R splice variants (Xiao, Jewell et al. 2008). It was therefore important to determine any potential deleterious effect of CB1R NT truncation in the CB1R$_{91\text{-}472}$ containing cell line.

**Radioligand binding**

The binding affinity of CP 55,940 and SR141716A was measured using equilibrium radioligand displacement experiments in order to compare the full length CB1R WT and the truncated CB1R$_{91\text{-}472}$ receptor (see Supplementary Figure 2A and B). CP 55,940's affinity for CB1R$_{91\text{-}472}$ was increased by 0.3 log units compared to WT. Although statistically significant, this is small change from the pharmacological perspective. The affinity of SR141716A was unaffected (see Supplementary Table ). It is possible that the kinetics (i.e., $k_{off}$ and $k_{on}$) of ligand binding may be different even when equilibrium measures of affinity appear the same. Hence, we studied the kinetics of [$^3$H]CP 55,940 binding to both the CB1 WT and the truncated CB1$_{91\text{-}472}$. The association and dissociation rates of [$^3$H]CP 55,940 binding to the CB1R WT and truncated CB1R$_{91\text{-}472}$ were very similar (Figure 2C and D) and their observed association rate ($k_{obs}$) and constant dissociation rate showed no statistically significant differences (see Supplementary Supplementary Table ).

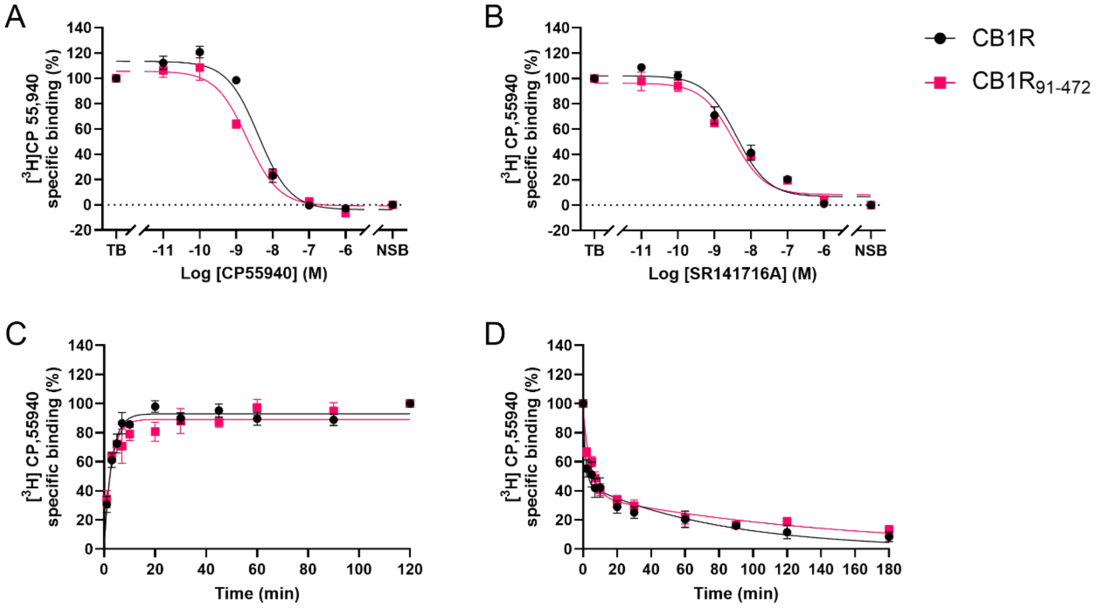


**Supplementary Figure 2.** Equilibrium binding and binding kinetics on CB1R and $CB1_{91-472}$ membranes Displacement of [$^3$H]CP 55,940 from CB1R and $CB1R_{91-472}$ in the presence of increasing concentration of CP 55,940 **(A)** and SR141716A **(B)**. Association **(C)** and dissociation **(D)** of [$^3$H]CP 55,940 from CB1R WT and $CB1R_{91-472}$. Results are presented as mean ± SEM of three independent experiments performed in duplicate.

**Supplementary Table 1.** Affinity and binding kinetic parameters on CB1R and $CB1R_{91-472}$ membranes determined in [$^3$H]CP 55,940 binding assays.

| | **p$IC_{50}$ ± SEM ($IC_{50}$ (nM))** CP55940 | **p$IC_{50}$ ± SEM ($IC_{50}$ (nM))** SR141716A | ***$k_{obs}$* ($min^{-1}$)** | ***$k_{off, fast}$* ($min^{-1}$)** | ***$k_{off, slow}$* ($min^{-1}$)** | **% fast** |
|---|---|---|---|---|---|---|
| **CB1R** | 8.4 ± 0.0 (4) | 8.4 ± 0.2 (4) | 0.37 ± 0.09 | 0.56 ± 0.19 | 0.012 ± 0.003 | 59 ± 5 |
| **$CB1R_{91-472}$** | 8.7 ± 0.1 (2)* | 8.5 ± 0.1 (3) | 0.36 ± 0.05 | 0.23 ± 0.01 | 0.007 ± 0.002 | 62 ± 5 |

Values represent mean ± SEM of three independent experiments performed in duplicate. Differences in p$IC_{50}$ and kinetic parameters values were analyzed using a two-tailed Student's t-test with Welch's correction. Differences compared to WT are denoted as follows: *$p < 0.05$.

**Gi-CASE cellular assay**

HU-210, a high affinity non-selective synthetic cannabinoid agonist, and the two endogenous cannabinoids 2-AG and AEA were tested for their ability to bind CB1R and induce an active receptor conformation. In the BRET-based Gi-CASE assay agonist stimulation of the receptors leads to the dissociation of the G$\alpha$ subunit (tagged with Nluc) from $\beta\gamma$ (the $\gamma$ subunit tagged with Venus) and a decrease in the BRET signal. There was no discernible difference in the agonist potencies comparing the full length CB1R (CB1R WT), CB1R and the truncated CB1R$_{91-472}$ (see Supplementary 3A and B, and Supplementary Table 2).

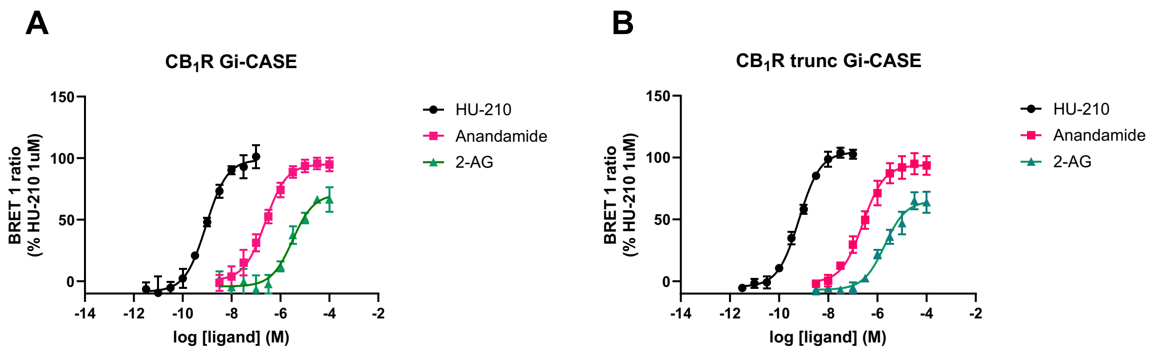


**Supplementary Figure 3.** Gi-CASE activation at the full-length or truncated CB1R$_{91-472}$ receptors when stimulated with HU-210 and the endocannabinoids 2-AG and anandamide. Concentration-response curves for HU-210, 2-AG and anandamide at the full-length receptor **(A)** or truncated receptor **(B)**. Data points and error bars represent the mean ± SEM of ≥ 3 independent experiments performed with duplicate technical replicates.

**Supplementary Table 2.** Gi-CASE cellular assay. Compound potency values expressed as pEC$_{50}$ for full length CB1R WT and truncated CB1R$_{91-472}$. Values were fitted from Supplementary Figure 3. Values represent mean ± SEM of ≥ 3 independent experiments.

| | **CB1R** | | **CB1R$_{91-472}$** | |
|---|---|---|---|---|
| | **p*EC*$_{50}$ (M)** | ***E*$_{max}$ (%)** | **p*EC*$_{50}$ (M)** | ***E*$_{max}$ (%)** |
| **HU-210** | 8.97 ± 0.08 | 98.2 ± 6.8 | 9.15 ± 0.09 | 104.1 ± 4.7 |
| **AEA** | 6.63 ± 0.12 | 97.0 ± 6.6 | 6.55 ± 0.13 | 95.3 ± 9.7 |
| **2-AG** | 5.54 ± 0.20 | 68.5 ± 8.3 | 5.51 ± 0.12 | 66.8 ± 9.5 |

Differences in pEC$_{50}$ and E$_{max}$ parameters values were analyzed using a paired t-test with differences compared to WT. No significant differences were observed between the p$EC_{50}$ or $E_{max}$ values $p > 0.05$.

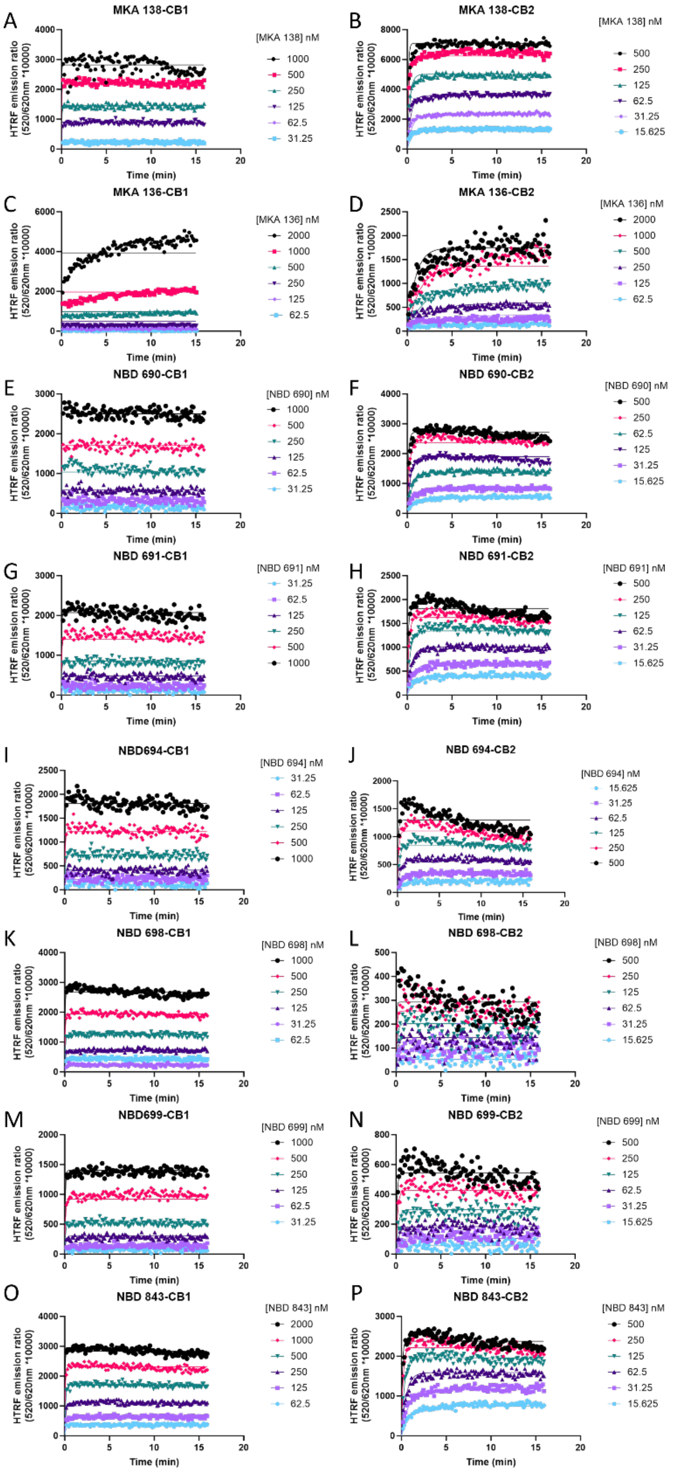


**Supplementary Figure 4. Determination of tracer kinetic binding parameters.** Kinetic association binding curves of fluorescent cannabinoid ligands CB1R (left column) and CB2R (right column) expressing membranes. Representative graphs show specific binding expressed as HTRF emission ratio for the tracer molecule MKA 138 **(A)** and **(B)**, MKA 136 **(C)** and **(D),** NBD 690 **(E)** and **(F),** NBD 691 **(G)** and **(H)**, NBD 694 **(I)** and **(J),** NBD 698 **(K)** and **(L),** NBD 699 **(M)** and **(N)**, and NBD 843 **(O)** and **(P)**, at CB1R and CB2R respectively.

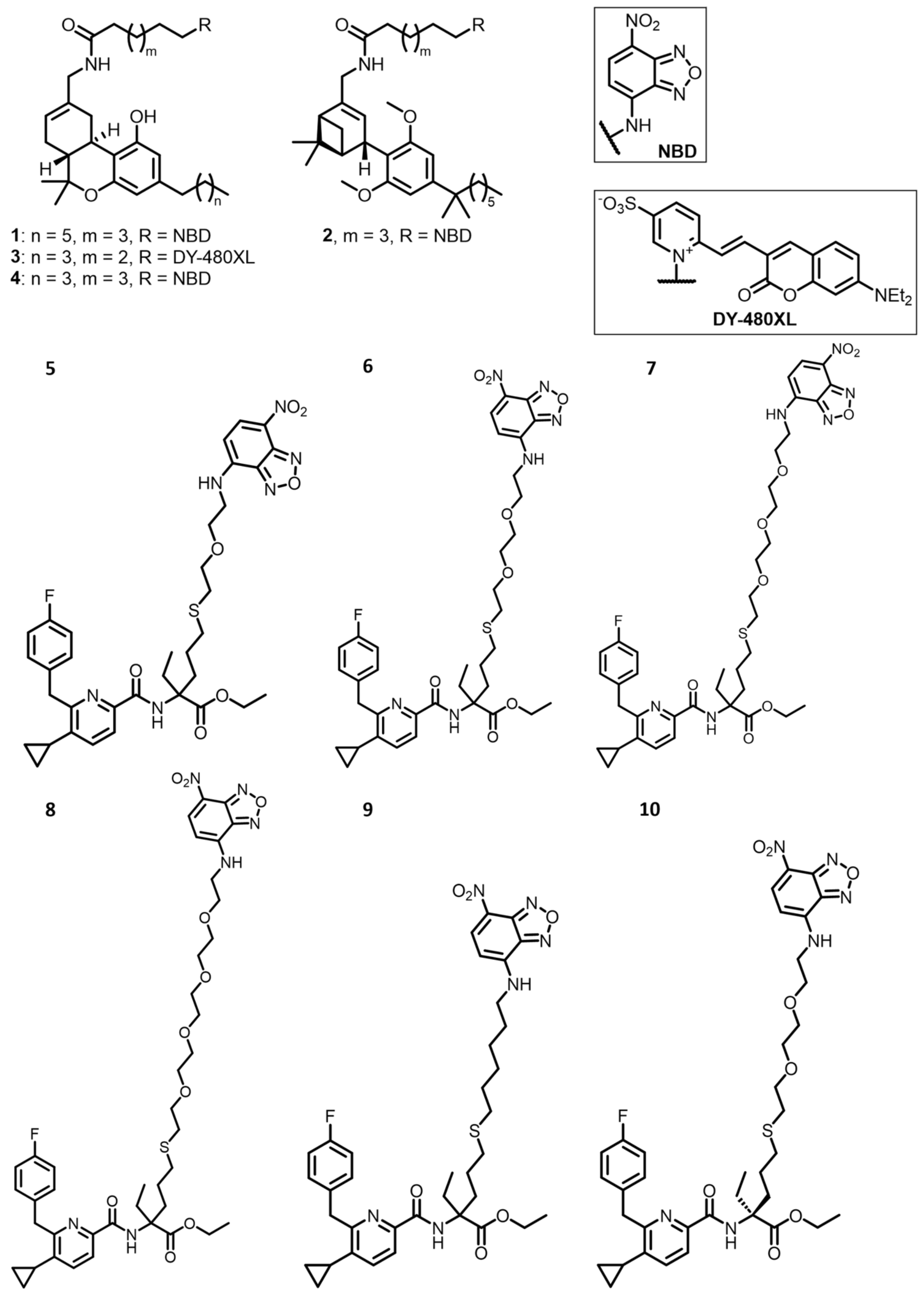


**Supplementary Figure 5.** Structures of TR-FRET tracers employed in the study. Numbers 1-10 in the figure correspond to the following compounds 1:D77; 2:MKA-115; 3:MKA-138; 4:MKA-136; 5:NBD-690; 6:NBD-691; 7:NBD-694; 8:NBD-698; 9:NBD-699; 10: NBD-843.

**Supplementary Table 3**. Affinity and binding kinetic parameters of fluorescent tracers binding to CB1R and CB2R. Data are expressed as mean ± SEM of ≥3 independent experiments conducted at 25°C. ND = Not determined.

| | **CB1R** | | | **CB2R** | | |
|---|---|---|---|---|---|---|
| | $k_{on}$ **($M^{-1}$ $min^{-1}$)** | $k_{off}$ **($min^{-1}$)** | $K_d$ **(M)** | $k_{on}$ **($M^{-1}$ $min^{-1}$)** | $k_{off}$ **($min^{-1}$)** | $K_d$ **(M)** |
| **MKA-138** | (1.9±0.5) $\times10^{7}$ | 9.0±1.8 | (4.9±0.6) $\times10^{-7}$ | (9±2) $\times10^{6}$ | 0.9±0.1 | (1.1±0.3) $\times10^{-7}$ |
| **MKA-136** | ND | ND | ND | (6.1±0.9) $\times10^{5}$ | 0.2±0.1 | (4±1) $\times10^{-7}$ |
| **NBD-690** | (1.1±0.2) $\times10^{7}$ | 7.7±0.4 | (7.7±1.4) $\times10^{-7}$ | (1.4±0.0) $\times10^{7}$ | 1.0±0.1 | (7.2±0.4) $\times10^{-8}$ |
| **NBD-691** | (1.3±0.2) $\times10^{7}$ | 12±1 | (9.2±0.2) $\times10^{-7}$ | (2.0±0.1) $x10^{7}$ | 1.2±0.06 | (6.3±0.1) $\times10^{-8}$ |
| **NBD-694** | (1.2±0.0) $\times10^{7}$ | 11.4±0.3 | (9.6±0.2) $\times10^{-7}$ | (2.1±0.2) $\times10^{7}$ | 2.3±0.2 | (1.1±0.0) $\times10^{-7}$ |
| **NBD-698** | (1.6±0.0) $\times10^{7}$ | 9.9±0.2 | (6.1±0.0) $\times10^{-7}$ | (2.4±0.4) $\times10^{7}$ | 2.8±0.4 | (1.2±0.1) $\times10^{-7}$ |
| **NBD-699** | (4.4±1.3) $\times10^{6}$ | 7.5±0.6 | (1.9±0.5) $\times10^{-6}$ | (1.5±0.2) $\times10^{7}$ | 1.7±0.5 | (1.2±0.4) $\times10^{-7}$ |
| **NBD-843** | (1.3±0.3) $\times10^{7}$ | 8.6±1.1 | (6.9±1.4) $\times10^{-7}$ | (1.5±0.1) $\times10^{7}$ | 1.2±0.6 | (8.6±3.1) $\times10^{-8}$ |

One of the key elements that will impact the fit of the Motulsky and Mahan model will be the binding kinetic characteristics of the tracer molecule, its association rate ($k_{on}$) and the dissociation rate ($k_{off}$). We performed a kinetic characterization of previously developed fluorescent tracers in order to characterise their kinetic binding properties towards CB1R and CB2R

The ligand MKA 136 showed a slow $k_{on}$ when binding to both CB1 and CB2, and equilibrium was not obtained in the 15 min data collection period, indicating that the time required for equilibrium was >15min. The slow association of MKA 136 makes it unsuitable for the use in the Motulsky and Mahan model, as a fast association is desired in order to observe significant competition with the unlabelled ligands at the very early time points. On the other hand, MKA 138, NBD 690, NBD 691, NBD 694 and NBD 698 show extremely fast association to CB1, although slightly slower to CB2 receptor. The fast $k_{on}$ observed at these receptors make it impractical to reliably measure the binding association phase, since probe binding equilibrium is reached almost instantly at the CB1R. Obtaining reliable data from the association phase of probe binding is crucial to apply the competition binding model, since the absence of early data points prior to reaching equilibrium leads to poorer kinetic parameters $k_{on}$ (K1, in the Motulsky Mahan model) and $k_{off}$, (K2) estimations thus increasing the errors associated with cold compound kinetic determinations $k_{on}$ (K3) and $k_{off}$, (K4). The kinetic profiles of MKA 138, NBD 690, NBD 691, NBD 694 and NBD 698 shown for their binding to CB2R could be considered suitable for the application of the Motulsky and Mahan modelling, as they exhibit moderately rapid association. However, a time-dependent decay of the HTRF ratio is observed for NBD 694 and NBD 698 binding to CB2R. And although less pronounced, the same signal decay can be observed for the NBD 690 and NBD 691 binding to CB2R.

**Chemical synthesis of D77**

Our synthesis of 1 (tracer D77) commenced with the acid-mediated FRIEDEL–CRAFTS allylation of phloroglucinol (**2**) with (*1R*)-(–)-myrtenol-derived allylic alcohol **3** according to a procedure previously reported by our group (Sarott, Viray et al. 2021). Upon exposure of **2** and **3** to *p*TsOH·$H_2O$, adduct **4** was obtained as a single diastereomer in 80% yield (see Supplementary Figure 5). Intermediate **5** was accessed by treatment of **4** with $FeCl_3$ to forge the central benzochromene motif of THC, followed by *para*-selective phenol triflation. Protection of the phenol as MOM ether and reductive cleavage of the pivalate ester yielded allylic alcohol **6**. To arrive at heterobifunctional linchpin **7**, compound **6** was converted to the corresponding primary amine in a sequence consisting of azide substitution under MITSUNOBU conditions and STAUDINGER reduction. The aryl triflate and primary amine in **7** served as orthogonal handles for late–stage functionalization with the affinity–controlling alkyl side chain as well as with the fluorophore of choice, respectively. NEGISHI cross-coupling of **7** with *n*-$C_7H_{15}ZnBr$ installed the $C_7$ alkyl side chain at C(3). The synthesis concluded with introduction of the nitrobenzoxadiazole (NBD) fluorophore by a PyBOP-mediated amide coupling with carboxylic acid **8**, followed by MOM ether deprotection to reveal TR-FRET tracer **1**.

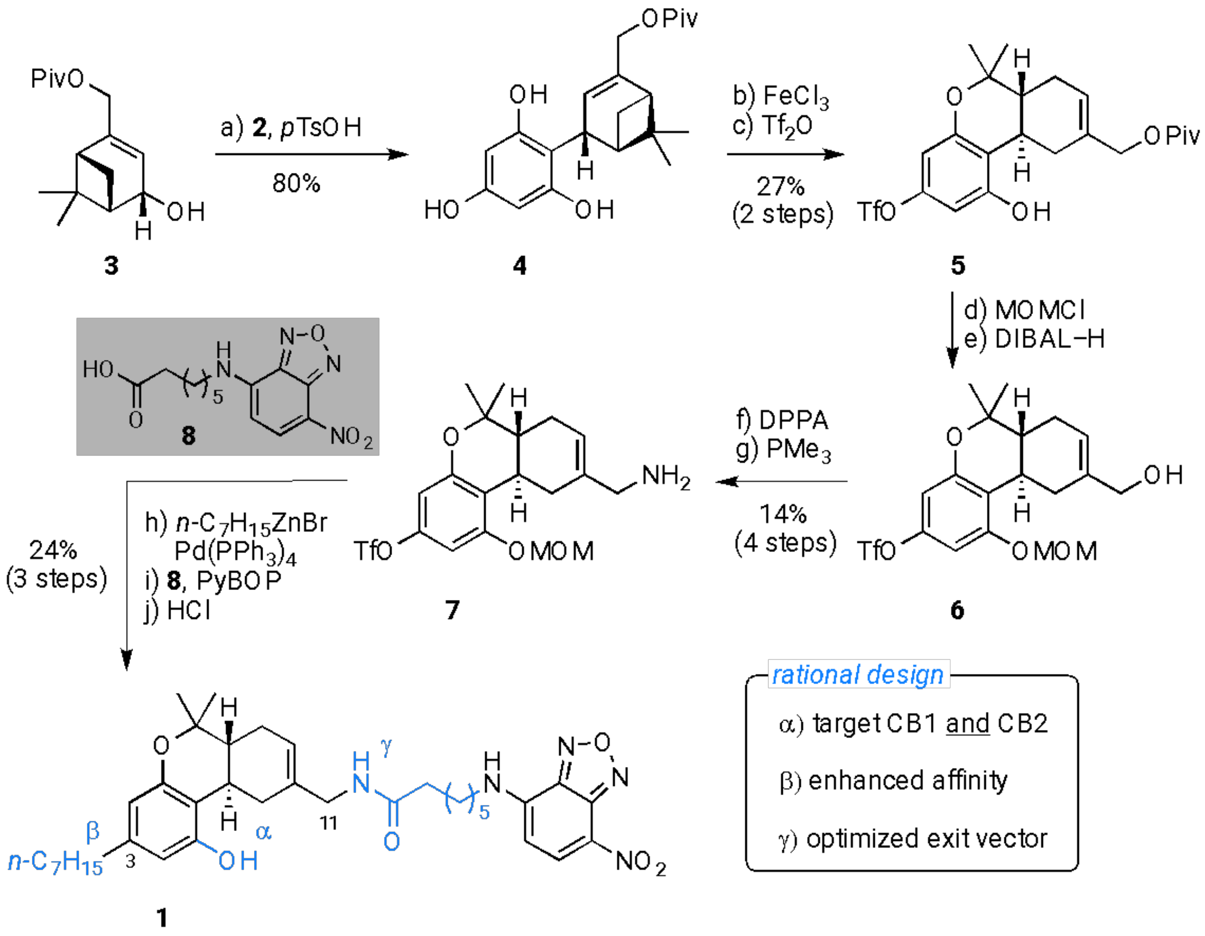


**Supplementary Figure 6.** Scheme of design and synthesis of THC-based fluorescent tracer D77 (1). Reagents and conditions: a) phloroglucinol (**2**), *p*TsOH·$H_2O$ (0.2 equiv), MeCN, rt, 80%; b) $FeCl_3$, $CH_2Cl_2$, 4 Å MS, 0 °C to rt; c) $Tf_2O$, 2,6-lutidine, $CH_2Cl_2$, –30 °C to rt, 27% over two steps; d) MOMCl, DIPEA, $CH_2Cl_2$, 0 °C; e) DIBAL–H, $CH_2Cl_2$, 0 °C; f) DIAD, $PPh_3$, $(PhO)_2P(O)N_3$, THF, 0 °C to rt; g) $PMe_3$, THF/$H_2O$ (10:1), rt, 14% over four steps; h) *n*-$C_7H_{15}$ZnBr, $Pd(PPh_3)_4$, THF, 70 °C, 39%; i) **8**, PyBOP, DIPEA, $CH_2Cl_2$, rt, 69%; j) HCl, MeOH, rt, 89%.

**Kinetic parameters of unlabelled cannabinoid ligands at CB1R and CB2R at room temperature**

The kinetic parameters of the cannabinoid compounds tested were also determined at room temperature (25 °C) following the Motulsky and Mahan approach and using the D77 fluorescent ligand as the tracer (600 nM and 900 nM for CB1R and CB2R respectively). The association of the tracer ligand was monitored using TR-FRET in competition with different concentrations of each cold cannabinoid ligand. Data were globally fitted using the model "kinetics of competitive binding" in GraphPad Prism and association rate-$k_{on}$ and dissociation rate-$k_{off}$ were calculated for the different cannabinoid compounds, which are found in Supplementary Table 3 and Table 4.

**Supplementary Table 4.** Kinetic parameters calculated from the Motulsky and Mahan experimental approach for the cannabinoid compounds tested for CB1R at room temperature (25°C). Data are expressed as mean ± SEM of 4 experiments (except HU-308, N=3) conducted independently.

| **CB1R** | $k_{on}$ | $k_{off}$ | **Residence time (Rt)** | |
|---|---|---|---|---|
| **(25°C)** | **($M^{-1}$ $min^{-1}$)** | **($min^{-1}$)** | **(min)** | **(s)** |
| **Rimonabant** | (1.2 ± 0.2) ×$10^8$ | 0.36 ± 0.03 | 2.9 ± 0.2 | 171 ± 14 |
| **HU-210** | (1.6 ± 0.1) ×$10^8$ | 0.38 ± 0.00 | 2.7± 0.0 | 160 ± 2 |
| **CP 55,940** | (1.0 ± 0.1) ×$10^8$ | 2.4 ± 0.2 | 0.42 ± 0.03 | 25 ± 2 |
| **2-AG** | (1.2 ± 0.2) ×$10^6$ | 4.8 ± 0.8 | 0.22 ± 0.03 | 13 ± 2 |
| **Anandamide** | (1.4 ± 0.2) ×$10^6$ | 3.8 ± 0.5 | 0.27 ± 0.03 | 17 ± 2 |
| **HU-308** | (4.0 ± 0.5) ×$10^5$ | 1.7 ± 0.2 | 0.60 ± 0.08 | 36 ± 5 |
| **SR 144528** | (1.2 ± 0.2) ×$10^7$ | 2.3 ± 0.2 | 0.44 ± 0.04 | 26 ± 3 |

**Supplementary Table 5**. Kinetic parameters calculated from the Motulsky and Mahan experimental approach for the cannabinoid compounds tested for CB2R at room temperature (25°C). Data are expressed as mean ± SEM of 4 experiments conducted independently.

| CB2R (25 °C) | $k_{on}$ ($M^{-1}$ $min^{-1}$) | $k_{off}$ ($min^{-1}$) | Residence time (Rt) | |
|---|---|---|---|---|
| | | | (min) | (s) |
| **Rimonabant** | (2.0 ± 0.3) ×$10^6$ | 2.0 ± 0.3 | 0.54 ± 0.08 | 32 ± 5 |
| **HU-210** | (6.42 ± 0.3) ×$10^7$ | 0.024 ± 0.005 | 50 ± 13 | 2971 ± 772 |
| **CP-55940** | (1.1 ± 0.1) ×$10^8$ | 0.33 ± 0.05 | 3.2 ± 0.4 | 191 ± 23 |
| **2-AG** | (6 ± 2) ×$10^6$ | 2.5 ± 0.6 | 0.5 ± 0.1 | 29 ± 7 |
| **Anandamide** | (1.4 ± 0.2) ×$10^6$ | 1.27 ± 0.2 | 0.8 ± 0.1 | 51 ± 8 |
| **HU-308** | (2.5 ± 0.4) ×$10^7$ | 0.42 ± 0.05 | 2.7 ± 0.2 | 164 ± 12 |
| **SR-144528** | (8.5 ± 0.4) ×$10^6$ | 0.16 ± 0.01 | 6.3 ± 0.4 | 379 ± 27 |

**Affinity values estimated from kinetic parameters**

The kinetic parameters obtained from the application of Motulsky and Mahan approach were used to calculate the kinetically derived affinity dissociation constant of the compounds tested by dividing the association and dissociation constants ($K_d$=$k_{off}$/$k_{on}$), shown Supplementary Tables 5 and 6.

**Supplementary Table 6.** Affinity values for the cannabinoid compounds tested at CB1R at 25 °C. The equilibrium dissociation constant $K_d$ was calculated from the kinetic parameters $k_{on}$ and $k_{off}$ obtained using the Motulsky and Mahan methodology. The data shown are mean ± SEM from 4 experiments (apart from HU-308, N=3) conducted independently.

| **CB1R (25 °C)** | **$K_d$ (nM)** | **p$K_d$ (M)** |
|---|---|---|
| **Rimonabant** | 3.2 ± 0.2 | 8.5 ± 0.03 |
| **HU-210** | 2.3 ± 0.1 | 8.6 ± 0.02 |
| **CP-55940** | 25 ± 2 | 7.61 ± 0.03 |
| **2-AG** | 4237 ± 728 | 5.40 ± 0.08 |
| **Anandamide** | 2710 ± 73 | 5.57 ± 0.01 |
| **HU-308** | 4308 ± 50 | 5.4 ± 0.0 |
| **SR-144528** | 190 ± 10 | 6.72 ±0.02 |

**Supplementary Table 7.** Affinity values for the cannabinoid compounds tested at CB2R at 25 °C. The equilibrium dissociation constant $K_d$ was calculated from the kinetic parameters $k_{on}$ and $k_{off}$ obtained using the Motulsky and Mahan methodology. The data shown are mean ± SEM from 4 experiments conducted independently.

| **CB2R (25 °C)** | **$K_d$ (nM)** | **p$K_d$ (M)** |
|---|---|---|
| **Rimonabant** | 1008 ± 95 | 6.00 ± 0.04 |
| **HU-210** | 0.38 ± 0.08 | 9.46 ± 0.12 |
| **CP-55940** | 2.8 ± 0.1 | 8.55 ± 0.01 |
| **2-AG** | 446 ± 57 | 6.36 ± 0.06 |
| **Anandamide** | 872 ± 31 | 6.06 ± 0.02 |
| **HU-308** | 18 ± 3 | 7.75 ± 0.08 |
| **SR-144528** | 1.9 ± 0.2 | 8.73 ± 0.05 |

### General synthetic and characterization methods for TR-FRET tracers.

#### Procedure

Unless otherwise noted, all reactions were carried out under nitrogen atmosphere. Compounds 5-10 shown in Supplementary Figure 5 were synthesized according to Gazzi, Brennecke et al. (2022)

#### Chemicals

All chemicals and solvents were purchased from commercial suppliers and were used without further purification. DMF, DMSO, THF, $Et_2O$, MeCN, and $CH_2Cl_2$ were dried using 4 Å molecular sieves or using an LC Technology Solutions solvent purification system (SP-1) under an atmosphere of dry nitrogen. *i*-$Pr_2NEt$ was distilled from KOH under an atmosphere of dry nitrogen.

#### Chromatography

Analytical thin-layer chromatography (TLC) was performed on Merck silica gel 60 $F_{254}$ TLC glass plates. Purification of reaction products was carried out by flash column chromatography (FCC) using Sigma Aldrich silica 230-400 mesh particle size, 60 Å under 0.3–0.5 bar overpressure or Büchi Pure Chromatography System C-805 Flash with FlashPure silica cartridges.

#### Nuclear Magnetic Resonance Spectroscopy

NMR spectra were acquired on Bruker AVIII HD 500 MHz and 400 MHz spectrometers operating at the denoted spectrometer frequency given in MHz for the specified nucleus. $^{1}H$ NMR spectra are reported with the solvent resonance as the reference ($CDCl_3$ at 7.26 ppm, $CD_3OD$ at 3.31 ppm, $CD_2Cl_2$ at 5.32 ppm, $(CD_3)_2SO$ at 2.50 ppm, $CD_3CN$ at 1.94 ppm). Peaks are reported as (s = singlet, bs = broad singlet, d = doublet, bt = broad triplet, t = triplet, q = quartet, m = multiplet or unresolved, coupling constant(s) in Hz, integral). $^{13}C$ NMR spectra were recorded with $^{1}H$-decoupling and are reported in ppm with the solvent resonance as the reference ($CDCl_3$ at 77.16 ppm, $CD_3OD$ at 49.00 ppm, $CD_2Cl_2$ at 54.00 ppm, $(CD_3)_2SO$ at 39.52 ppm, $CD_3CN$ at 1.32 ppm). Service measurements were performed by the NMR service team of the Laboratorium für Organische Chemie at ETH Zürich.

#### High-Resolution Mass Spectrometry

High-resolution mass spectrometric data were obtained at ETH Zürich mass spectrometry service on Bruker Daltonics maXis ESI-QTOF, Thermo Q Exactive EI-Trace 1310 Analyser or a Bruker Daltonics maXiS II ESI-QTOF spectrometers and are reported as (*m/z*).

#### Infrared Spectroscopy

Infrared (IR) spectra were measured neat on a Perkin-Elmer UATR Two FT-IR Spectrometer and the band maxima are reported in wavenumbers ($cm^{-1}$).

#### Optical Rotation

Optical rotations ($[\alpha]_D^T$) were determined using a Jasco P-2000 Polarimeter (10 cm, 1.5 mL cell).

### SYNTHETIC SCHEMES

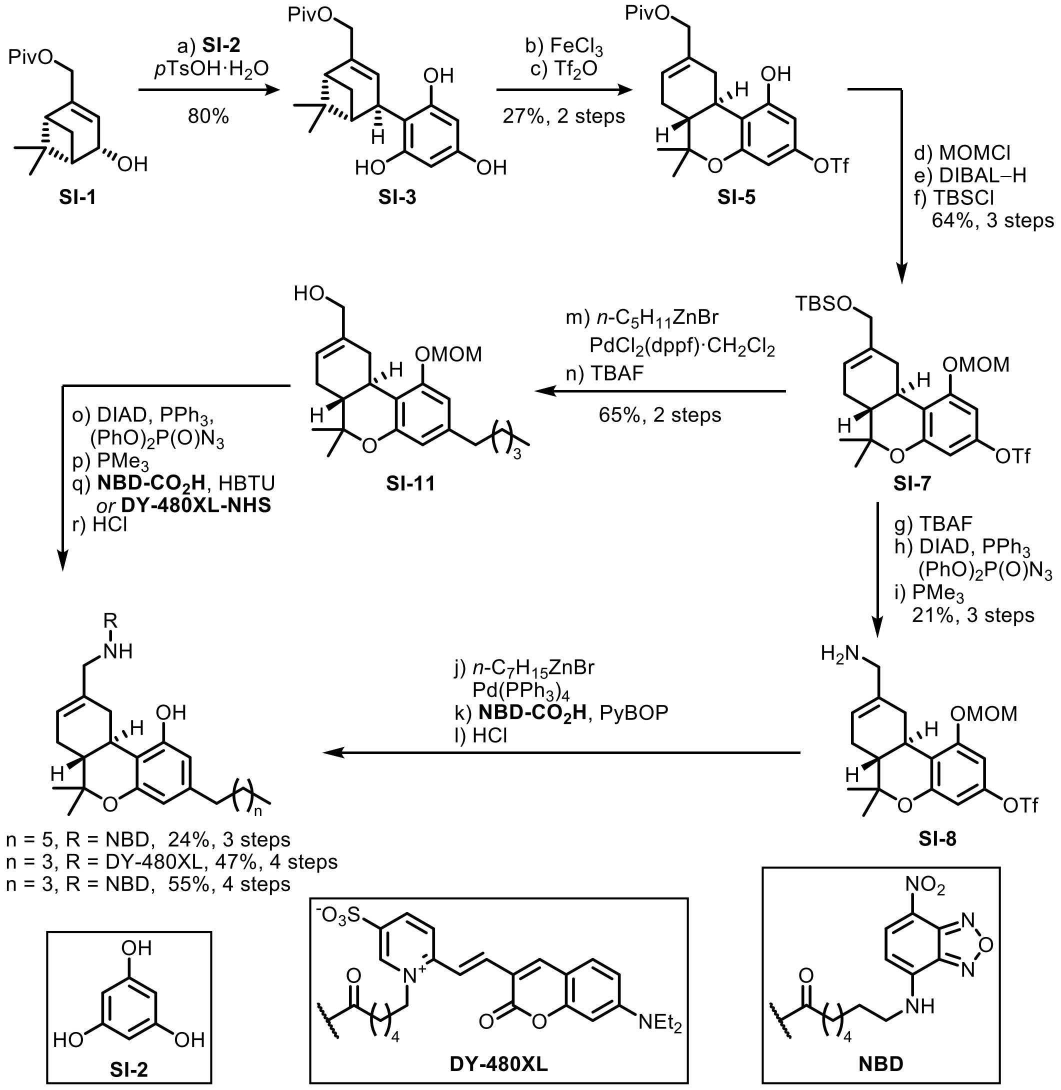


**Scheme S1.** Synthesis of $\Delta^8$-THC-derived fluorescent tracers via a heterobinfunctional derivative.[a]

[a]Reagents and conditions: a) phloroglucinol **SI-2**, *p*TsOH·$H_2O$, MeCN, rt, 80%; b) $FeCl_3$, $CH_2Cl_2$, 4 Å MS, 0 °C to rt; c) $Tf_2O$, 2,6-lutidine, $CH_2Cl_2$, –30 °C to rt, 27% over two steps; d) MOMCl, *i*-PrNEt$_2$, $CH_2Cl_2$, 0 °C, quant.; e) DIBAL–H, $CH_2Cl_2$, 0 °C; f) TBSCl, $NEt_3$, DMAP, $CH_2Cl_2$, rt, 64% over two steps; g) TBAF, THF, rt, 45%; h) DIAD, $PPh_3$, $(PhO)_2P(O)N_3$, THF, 0 °C to rt; i) $PMe_3$, THF, $H_2O$, rt, 48% over two steps; j) *n*-$C_7H_{15}$ZnBr, $Pd(PPh_3)_4$, THF, 70 °C, 39%; k) **NBD-CO₂H**Error! Reference source not found., PyBOP, *i*-PrNEt$_2$, $CH_2Cl_2$, rt, 69%; l) HCl, MeOH, rt, 89%; m) *n*-$C_5H_{11}$ZnBr, TBAI, $PdCl_2$(dppf)·$CH_2Cl_2$, THF, NMP, 60 °C, quant.; n) TBAF, THF, 0 °C, 65%; o) DIAD, $PPh_3$, $(PhO)_2P(O)N_3$, THF, 0 °C to rt, 75%; p) $PMe_3$, THF, $H_2O$, rt; q)Error! Reference source not found. **DY-480XL-NHS**, *i*-PrNEt$_2$, DMF, rt *or* **NBD-CO$_2$H**, HBTU, *i*-PrNEt$_2$, $CH_2Cl_2$, rt; r) HCl, MeOH, $Et_2O$, rt, 63–73% over three steps.

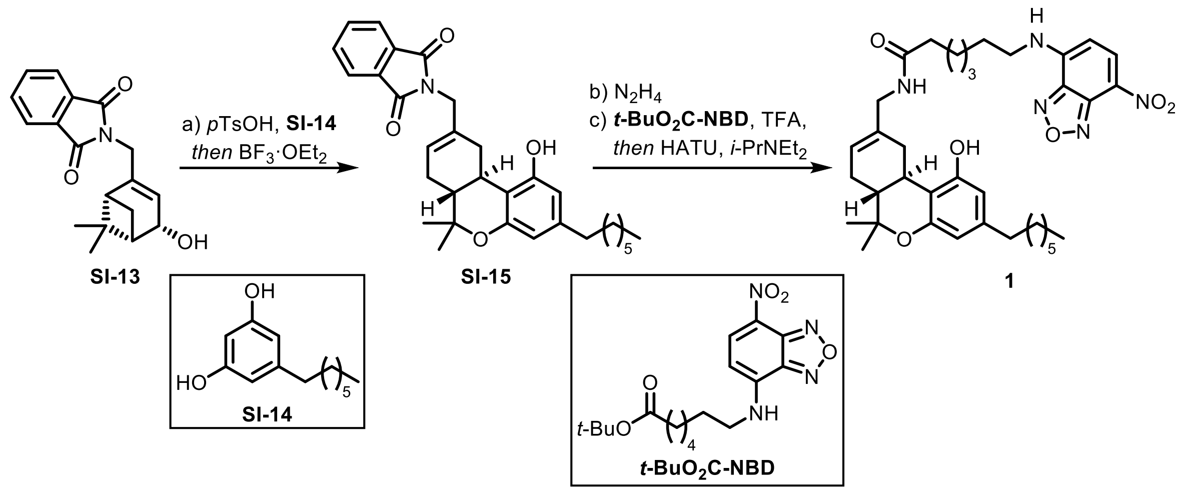


**Scheme S2.** Optimized synthetic strategy toward probe **1**.[a]

[a]Reagents and conditions: a) spherophorol **SI–14**, *p*TsOH·$H_2O$, $CH_2Cl_2$, rt *then* $BF_3$·$Et_2O$, –20 °C to 0 °C, 56%; b) $N_2H_4$·$H_2O$, (*E*/*Z*)-crotyl alcohol, EtOH, 75 °C, 75%; c) ***t*-BuO₂C-NBD**, TFA *then* HATU, *i*-Pr$NEt_2$, DMF, rt, 81%.

### COMPOUND SYNTHESIS AND CHARACTERIZATION

Synthesis of **SI-3**

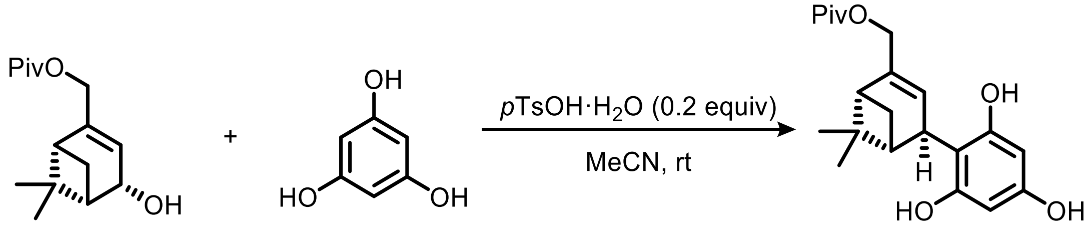


**SI-1** **SI-2** (10 equiv) **SI-3**

Allylic alcohol **SI-1** (1.9 g, 7.4 mmol, 1.0 equiv) and phloroglucinol (**SI-2**) (9.4 g, 0.074 mol, 10 equiv) were dissolved in MeCN (300 mL) and *p*TsOH·$H_2O$ (0.28 g, 1.5 mmol, 0.2 equiv) was added. The resulting solution was stirred at ambient temperature. After 30 min, TLC showed full conversion of limiting allylic alcohol reagent and the reaction was stopped by addition of brine (200 mL). After separation of the organic layer, the aqueous phase was extracted with EtOAc (6 × 75 mL). The combined organic extracts were washed with brine, dried over $Na_2SO_4$ and concentrated under reduced pressure. The unpurified solid was extracted with hot chloroform (5 × 50 mL, 55 °C), filtered, and the filtrate concentrated under reduced pressure. Purification by flash column chromatography ($SiO_2$; using 10 – 20% EtOAc in hexanes) afforded the product **SI-3** as off-white solid (2.1 g, 80%).

**$^1$H NMR** (400 MHz, $CD_3CN$) δ = 6.71 (br, 3H), 5.93 – 5.86 (m, 1H), 5.82 (s, 2H), 4.68 – 4.40 (m, 2H), 3.90 (t, $J$ = 2.5 Hz, 1H), 2.34 – 2.24 (m, 2H), 2.14 – 2.08 (m, 1H), 1.42 (d, $J$ = 9.1 Hz, 1H), 1.30 (s, 3H), 1.18 (s, 9H), 0.93 (s, 3H). **$^{13}$C NMR** (101 MHz, $CD_3CN$) δ = 179.1, 157.7, 157.4, 148.0, 122.3, 107.8, 96.1, 67.3, 48.3, 44.8, 41.40, 39.5, 38.3, 28.3, 27.4, 26.2, 21.1. **IR** (neat, $\nu_{max}$/cm$^{-1}$) 3420, 2976, 2936, 2872, 1704, 1628, 1606, 1516, 1465, 1399, 1384, 1367, 1284, 1232, 1162, 1143, 1074, 1030, 1009, 940, 881, 824, 773, 741, 723, 637, 527, 507. **HRMS (ESI)**: $m/z$ = 383.1830 [M+Na]$^+$ (calc. for $C_{21}H_{28}NaO_5$ $m/z$ = 383.1834). **$[\alpha]^{23}_D$** = –93.994 ± 0.198 (c = 1.0, MeOH).

Synthesis of **SI-5**

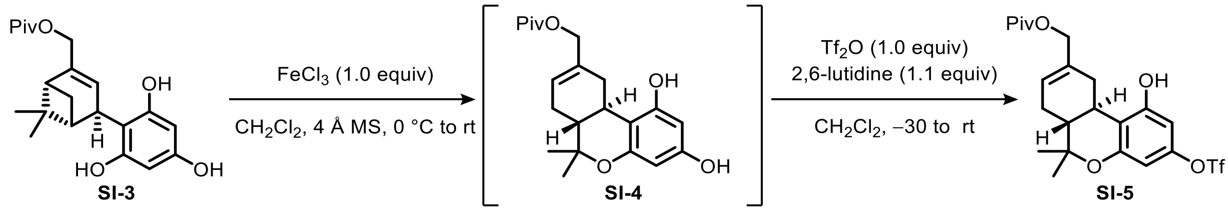


To a suspension of **SI-3** (1.4 g, 4.0 mmol, 1.0 equiv) and 1.5 g activated 4 Å MS in anhydrous $CH_2Cl_2$ (16 mL) at 0 °C was added anhydrous $FeCl_3$ (0.65 g, 4.0 mmol, 1.0 equiv) in three batches. The reaction mixture was then allowed to reach ambient temperature during 90 min, after which full conversion of starting material was observed. The reaction mixture was then filtered through a pad of Celite to remove the molecular sieves. The filtrate was washed with water and the aqueous layer was extracted with EtOAc (3 × 50 mL). The combined organic layers were washed with brine, dried over $Na_2SO_4$ and concentrated under reduced pressure. Purification of the crude material by flash column chromatography ($SiO_2$; using 20% EtOAc in hexanes) afforded bisphenol intermediate **SI-4** as light-brown foam (0.91 g, 80% pure, approx.50% yield). Intermediate **SI-4** was used for subsequent triflation without further purification and reagent equivalents were based on the estimated 50% yield in the cyclization step. Thus, to a solution of **SI-4** in anhydrous $CH_2Cl_2$ (40 mL) at –30 °C (acetone/dry ice) was added 2,6-lutidine (0.26 mL, 2.2 mmol, 1.1 equiv) and then a solution of triflic anhydride (0.34 mL, 2.0 mmol, 1.0 equiv) in anhydrous $CH_2Cl_2$ (0.5 mL). The resulting orange-yellow solution was allowed to reach ambient temperature in the course of 2 h, after which full conversion was observed. The reaction was stopped by addition of water (20 mL) and after separation of the aqueous layer the organic phase was washed with 1 M HCl, aq. sat. $NaHCO_3$ and brine (10 mL, each). The organic extracts were dried over $Na_2SO_4$ and concentrated under reduced pressure. Purification of the crude material by flash column chromatography ($SiO_2$; using 3 – 10% EtOAc in hexanes) afforded the product **SI-5** as white foam (0.54 g, 27% over 2 steps).

**$^1$H NMR** (400 MHz, $CDCl_3$) δ = 6.34 (d, *J* = 2.5 Hz, 1H), 6.28 (d, *J* = 2.5 Hz, 1H), 5.78 – 5.73 (m, 1H), 4.50 (s, 2H), 3.33 (dd, *J* = 16.5, 4.3 Hz, 1H), 2.75 – 2.67 (m, 1H), 2.29 – 2.21 (m, 1H), 1.95 – 1.76 (m, 3H), 1.39 (s, 3H), 1.23 (s, 9H), 1.09 (s, 3H). **$^{13}$C NMR** (101 MHz, $CDCl_3$) δ = 179.3, 156.3, 156.0, 148.4, 133.5, 123.1, 118.8 (q, *J* = 320.7 Hz), 113.2, 103.3, 100.8, 77.76, 68.1, 44.5, 39.1, 31.4, 31.3, 27.6, 27.4, 27.4, 18.6. **$^{19}$F NMR** (376 MHz, $CDCl_3$) δ = -73.0 (decoupled). **IR** (neat, $\nu_{max}$/cm$^{-1}$) 3374, 2978, 1700, 1598, 1421, 1209, 1180, 1141, 985, 865, 847, 759, 614.

**HRMS (ESI)**: *m/z* = 493.1507 $[M+H]^+$ (calc. for $C_{22}H_{28}F_3O_7S$ *m/z* = 493.1502). $[\alpha]^{23}_D$ = –145.512 ± 0.147 (c = 2.0, $CHCl_3$).

Synthesis of **SI-6**

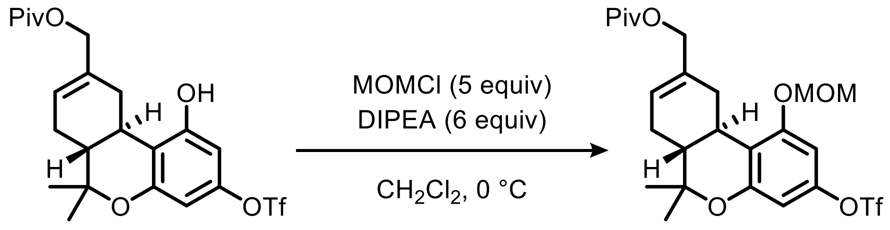


**SI-5** **SI-6**

To a solution of **SI-5** (0.052 g, 0.11 mmol, 1 equiv) in $CH_2Cl_2$ (1.1 mL) at 0 °C was added DIPEA (0.11 mL, 0.63 mmol, 6 equiv) and MOMCl (0.040 mL, 0.53 mmol, 5 equiv). The resulting solution was stirred at 0 °C for 60 min, after which TLC analysis showed consumption of starting material. The reaction was stopped by addition of aq. sat. $NaHCO_3$ cooled to ambient temperature and aq. sat. $NaHCO_3$ was added. The aqueous layer was extracted with $CH_2Cl_2$ and the combined organic extracts were washed with brine, dried over $Na_2SO_4$, and concentrated under reduced pressure to afford the product **SI-6** as a colorless wax (0.056 g, quantitative).

**$^1$H NMR** (500 MHz, $CDCl_3$) δ = 6.60 (d, *J* = 2.5 Hz, 1H), 6.45 (d, *J* = 2.5 Hz, 1H), 5.80 – 5.77 (m, 1H), 5.19 – 5.13 (m, 2H), 4.55 (d, *J* = 12.5 Hz, 1H), 4.44 (d, *J* = 12.4 Hz, 1H), 3.47 (s, 3H), 3.29 (dd, *J* = 17.0, 4.7 Hz, 1H), 2.71 (td, *J* = 11.0, 4.6 Hz, 1H), 2.29 – 2.19 (m, 1H), 1.95 – 1.76 (m, 2H), 1.40 (s, 3H), 1.22 (s, 9H), 1.10 (s, 3H). **$^{13}$C NMR** (126 MHz, $CDCl_3$) δ = 178.4, 157.5, 155.5, 148.8, 133.8, 123.8, 115.1, 104.7, 99.9, 94.7, 77.6, 68.1, 56.5, 44.7, 39.0, 31.8, 31.7, 27.7, 27.4, 27.4, 18.5, ($CF_3$ too weak). **$^{19}$F NMR** (471 MHz, $CDCl_3$) δ = -73.0. **IR** (neat, $\nu_{max}$/cm$^{-1}$) 2977, 1729, 1604, 1423, 1212, 1143, 1055, 992. **HRMS (ESI)**: *m/z* = 559.1580 $[M+Na]^+$ (calc. for $C_{24}H_{31}F_3NaO_8S$ *m/z* = 559.1584).

Synthesis of **SI-7**

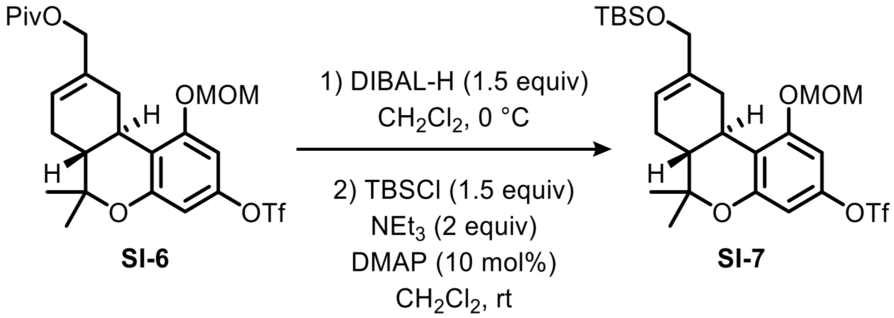


To a solution of **SI-6** (0.056 g, 0.10 mmol, 1.0 equiv) in $CH_2Cl_2$ (1.1 mL) at 0 °C was added DIBAL-H (1 M in hexanes, 0.16 mL, 0.16 mmol, 1.5 equiv) and the reaction mixture was allowed to reach ambient temperature. After 30 min, TLC (20% EtOAc in hexanes) showed full consumption of starting material. The reaction was stopped by addition of Rochelle's salt solution (1 mL) and the mixture was vigorously stirred until phase separation occurred. The aqueous phase was extracted with $CH_2Cl_2$ and the combined organic extracts were washed with brine, dried over $Na_2SO_4$ and concentrated under reduced pressure. The resulting yellowish oil was taken up in $CH_2Cl_2$ (1.0 mL) and TBSCl (0.024 g, 0.16 mmol, 1.5 equiv), $NEt_3$ (0.030 mL, 0.21 mmol, 2.0 equiv) as well as DMAP (1.3 mg, 0.010 mmol, 10 mol%) were added. The reaction solution was stirred at ambient temperature for 15 h. Water (1 mL) was added, the aqueous layer was extracted with $CH_2Cl_2$ and the combined organic extracts were washed with brine, dried over $Na_2SO_4$, and concentrated under reduced pressure. Purification by flash column chromatography ($SiO_2$; using 5% $Et_2O$ in hexanes) afforded the product **SI-7** as colorless oil (0.038 g, 64% over two steps).

**$^1$H NMR** (400 MHz, $CDCl_3$) δ = 6.60 (d, *J* = 2.5 Hz, 1H), 6.44 (d, *J* = 2.5 Hz, 1H), 5.71 (s, 1H), 5.17 (s, 2H), 4.04 (s, 2H), 3.47 (d, *J* = 0.8 Hz, 3H), 3.28 (dd, *J* = 17.0, 4.6 Hz, 1H), 2.69 (td, *J* = 10.8, 4.6 Hz, 1H), 2.22 (d, *J* = 13.4 Hz, 1H), 1.94 – 1.69 (m, 3H), 1.39 (s, 3H), 1.09 (s, 3H), 0.92 (s, 9H), 0.08 (d, *J* = 1.2 Hz, 6H). **$^{13}$C NMR** (101 MHz, $CDCl_3$) δ = 157.6, 155.5, 148.7, 138.0, 120.0, 115.5, 104.7, 99.9, 94.7, 77.8, 67.1, 56.4, 45.1, 31.8, 31.4, 27.7, 27.5, 26.1, 18.6, -5.1, -5.1, ($CF_3$ too weak). **$^{19}$F NMR** (471 MHz, $CDCl_3$) δ = -73.0. **IR** (neat, $\nu_{max}/cm^{-1}$) 2931, 2857, 1604, 1474, 1424, 1246, 1212, 1143, 1107, 1052, 992, 838. **HRMS (ESI)**: *m/z* = 589.1864 $[M+Na]^+$ (calc. for $C_{25}H_{37}F_3NaO_7SSi$ *m/z* = 589.1874). **$[\alpha]^{25}_D$** = –133.767 ± 0.080 (c = 2.0, $CHCl_3$)

Synthesis of **SI-8**

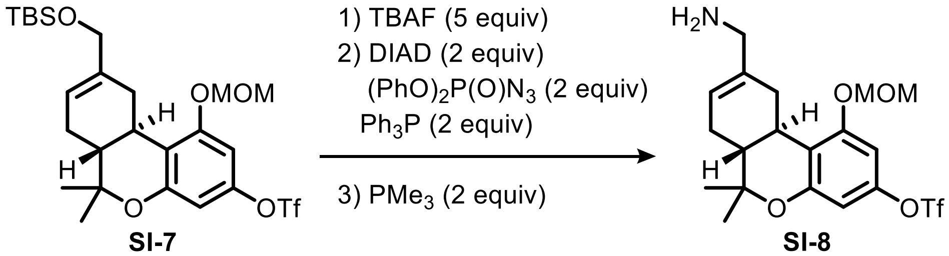


To a solution of **SI-7** (0.35 g, 0.61 mmol, 1.0 equiv) in THF (6.1 mL) was added TBAF (1 M in THF, 3.0 mL, 3.0 mmol, 5.0 equiv) and the residual solution was stirred at ambient temperature for 2 h. Aq. sat. $NaHCO_3$ was added, the aqueous layer was extracted with $CH_2Cl_2$ and the combined organic extracts were dried over $Na_2SO_4$ and concentrated under reduced pressure. Purification by flash column chromatography ($SiO_2$; using hexanes to 20% EtOAc in hexanes) afforded the intermediate allylic alcohol (0.12 g, 0.28 mmol).

To a solution of $PPh_3$ (0.15 g, 0.55 mmol, 2.0 equiv) in THF (1.8 mL) at 0 °C was added DIAD (0.11 mL, 0.55 mmol, 2.0 equiv) and the yellow suspension was stirred at 0 °C for 10 min. A solution of allyl alcohol intermediate (0.12 g, 0.28 mmol, 1.0 equiv) in THF (0.9 mL) was added, then the mixture was removed from the ice bath and $(PhO)_2P(O)N_3$ (0.12 mL, 0.55 mmol, 2.0 equiv) was added immediately. The resulting mixture was stirred at ambient temperature for 60 min. When TLC (10% EtOAc in hexanes) showed full consumption of starting material, the reaction mixture was concentrated under reduced pressure. Attempted purification by flash column chromatography ($SiO_2$; using hexanes to 5% $Et_2O$ in hexanes) failed to separate product from DIAD residues, and thus the mixture was taken up in 10:1 THF/water (2.5 mL, separately sparged with $N_2$). $PMe_3$ (1 M in THF, 0.55 mL, 0.55 mmol, 2.0 equiv) and the mixture was stirred at ambient temperature overnight. After concentration under reduced pressure, purification by flash column chromatography ($SiO_2$; using 1 to 15% MeOH in $CH_2Cl_2$, 1% 7 N $NH_3$ in MeOH) afforded the product **SI-8** together with $POMe_3$ impurity as yellowish oil (0.060 g, 48% over two steps).

**$^1$H NMR** (500 MHz, $CDCl_3$) δ = 6.57 (d, $J$ = 2.5 Hz, 1H), 6.44 (d, $J$ = 2.5 Hz, 1H), 5.66 (d, $J$ = 5.1 Hz, 1H), 5.18 (s, 2H), 3.48 (s, 3H), 3.27 – 3.23 (m, 2H), 3.23 – 3.15 (m, 1H), 2.70 (td, $J$ = 11.0, 4.6 Hz, 1H), 2.36 – 2.13 (m, 3H), 1.91 – 1.77 (m, 3H), 1.39 (s, 3H), 1.09 (s, 3H). **$^{19}$F NMR** (471 MHz, $CDCl_3$) δ = -73.0. **$^{13}$C NMR** (126 MHz, $CDCl_3$) δ = 157.4, 155.5, 148.7, 138.7, 119.4, 118.8 (q, $J$ = 231 Hz), 115.3, 104.7, 99.8, 94.8, 77.7, 56.5, 47.6, 45.0, 32.8, 31.9,

27.7, 27.5, 18.5. **IR** (neat, $\nu_{max}$/cm$^{-1}$) 3370, 2912, 1603, 1476, 1422, 1246, 1212, 1142, 1107, 991. **HRMS (ESI)**: *m/z* = 452.1346 $[M+H]^+$ (calc. for $C_{19}H_{25}F_3NO_6S$ *m/z* = 452.1349).

Synthesis of **SI-9**

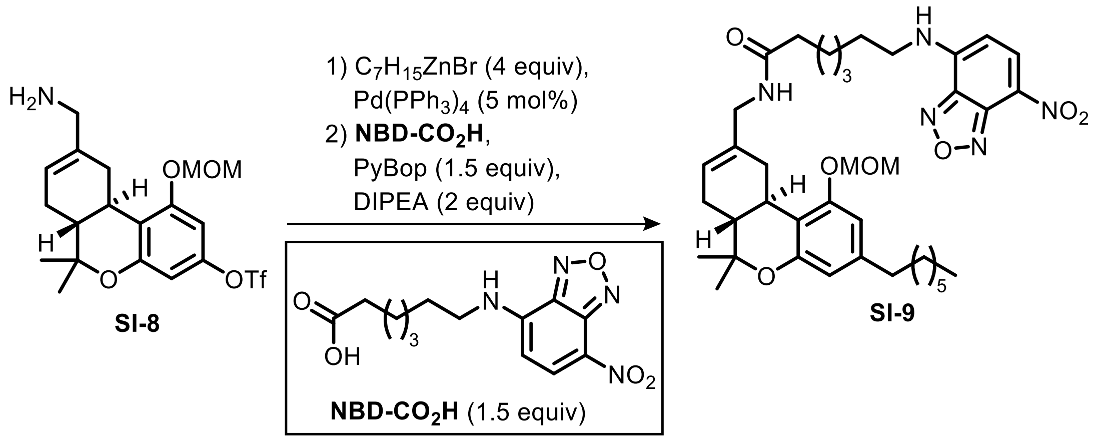


To a solution of **SI-8** (0.020 g, 0.044 mmol, 1.0 equiv) and $Pd(PPh_3)_4$ 2.6 mg, 2.2 μmol, 5 mol) in THF (0.4 mL) was added heptylzincbromide (0.5 M in THF, 0.35 mL, 0.18 mmol, 4 equiv) and the dark solution was stirred at 70 °C overnight. After cooling to ambient temperature, aq. sat. $NaHCO_3$ was added, the aqueous layer was extracted with $CH_2Cl_2$ and the combined organic extracts were dried over $Na_2SO_4$ and concentrated under reduced pressure. Purification by flash column chromatography ($SiO_2$; using 5% MeOH in $CH_2Cl_2$, 1% 7N $NH_3$ in MeOH) afforded the cross-coupling product in moderate purity (7.0 mg, 0.017 mmol, 39%). This residue was taken up in $CH_2Cl_2$ (0.2 mL) without further purification, and NBD acid **NBD-CO₂H** (8.4 mg, 0.026 mmol, 1.5 equiv), PyBop (7.0 mg, 0.026 mmol, 1.5 equiv) and DIPEA (6.0 μL, 0.035 mmol, 2.0 equiv) were added. The orange reaction mixture was stirred at ambient temperature for 72 h. Water was added and aqueous layer was extracted with $CH_2Cl_2$. The combined organic extracts were dried over $Na_2SO_4$ and concentrated under reduced pressure. Purification by preparative TLC ($SiO_2$; using 5% MeOH in $CH_2Cl_2$) afforded the product **SI-9** as an orange solid (6.0 mg, 69%).

**$^1$H NMR** (400 MHz, $CDCl_3$) δ = 8.49 (d, $J$ = 8.6 Hz, 1H), 6.44 (d, $J$ = 1.6 Hz, 1H), 6.34 (d, $J$ = 1.7 Hz, 1H), 6.16 (d, $J$ = 8.6 Hz, 1H), 5.61 (d, $J$ = 5.0 Hz, 1H), 5.47 – 5.39 (m, 1H), 5.18 (d, $J$ = 6.5 Hz, 1H), 5.14 (d, $J$ = 6.5 Hz, 1H), 3.84 (d, $J$ = 5.6 Hz, 2H), 3.48 (s, 3H), 3.47 – 3.44 (m, 2H), 3.25 (dd, $J$ = 15.9, 4.5 Hz, 1H), 2.69 (td, $J$ = 10.9, 4.6 Hz, 1H), 2.50 – 2.43 (m, 2H), 2.20 (q, $J$ = 7.7, 7.2 Hz, 3H), 1.85 – 1.75 (m, 5H), 1.69 (t, $J$ = 7.3 Hz, 2H), 1.56 (m, 2H), 1.51 – 1.43 (m, 4H), 1.37 (s, 3H), 1.31 – 1.25 (m, 8H), 1.09 (s, 3H), 0.90 – 0.84 (m, 3H). **$^{13}$C NMR** (101 MHz, $CDCl_3$) δ = 172.6, 156.7, 154.5, 144.4, 144.1, 143.2, 136.6, 135.5, 121.5, 112.2, 111.3, 106.4, 98.7, 94.6, 76.5, 56.5, 45.3, 44.0, 36.5, 36.1, 33.2, 32.0, 31.8, 31.2, 29.6, 29.3, 28.8, 28.3, 27.9, 27.7, 26.7, 25.4, 22.8, 18.5, 14.3. **IR** (neat, $\nu_{max}$/cm$^{-1}$) 3295, 2927, 2856, 1650,

1619, 1580, 1444, 1299, 1273, 1051. **HRMS (ESI)**: *m/z* = 714.3831 $[M+Na]^+$ (calc. for $C_{38}H_{53}N_5NaO_7$ *m/z* = 714.3837).

Synthesis of **1**

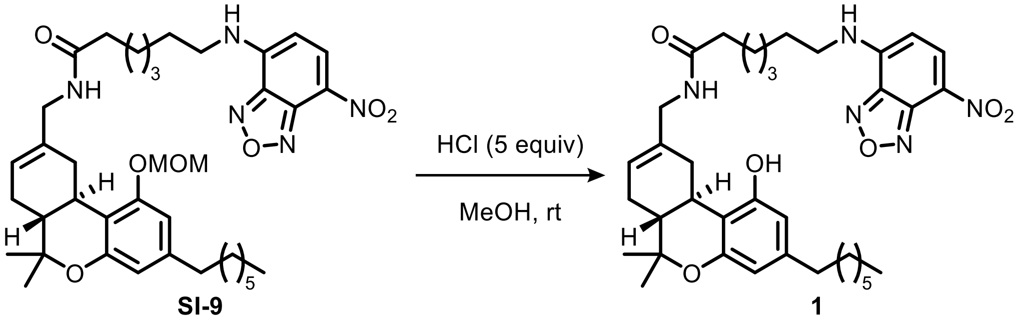

To a solution of **SI-9** (6.0 mg, 8.7 µmol, 1.0 equiv) in MeOH (0.1 mL) was added HCl (2 M in THF, 0.022 mL, 0.043 mmol, 5.0 equiv) and the orange solution was stirred at ambient temperature for 2 h. After TLC (10% MeOH in $CH_2Cl_2$) showed full consumption of starting material, aq. sat. $NaHCO_3$ was added, the aqueous layer was extracted with $CH_2Cl_2$ and the combined organic extracts were dried over $Na_2SO_4$ and concentrated under reduced pressure. Purification by pipette chromatography ($SiO_2$; using 1 to 5% MeOH in $CH_2Cl_2$,) afforded the product **1** as an orange wax (5.0 mg, 89%).

**$^1$H NMR** (500 MHz, $CDCl_3$) δ = 8.47 (d, *J* = 8.7 Hz, 1H), 6.57 (s, 1H), 6.29 – 6.21 (m, 1H), 6.18 – 6.12 (m, 2H), 5.63 – 5.50 (m, 2H), 3.81 (d, *J* = 20.8 Hz, 2H), 3.46 (m, 2H), 3.38 – 3.27 (m, 1H), 2.67 (td, *J* = 10.9, 4.6 Hz, 1H), 2.39 (td, *J* = 7.4, 3.6 Hz, 2H), 2.24 (t, *J* = 7.1 Hz, 2H), 2.21 – 2.14 (m, 1H), 1.85 – 1.75 (m, 5H), 1.73 – 1.66 (m, 2H), 1.56 – 1.40 (m, 6H), 1.36 (s, 3H), 1.30 – 1.21 (m, 8H), 1.08 (s, 3H), 0.88 – 0.84 (m, 3H). **$^{13}$C NMR** (126 MHz, $CDCl_3$) δ = 173.3, 155.3, 154.9, 144.4, 144.1, 143.1, 136.7, 135.3, 122.0, 110.1, 109.9, 107.9, 98.8, 76.5, 45.5, 45.0, 44.0, 36.6, 35.7, 32.7, 31.9, 31.6, 31.1, 29.5, 29.3, 28.7, 28.2, 27.8, 27.7, 26.6, 25.4, 22.8, 18.6, 14.3. **IR** (neat, $\nu_{max}$/cm$^{-1}$) 3320, 2926, 2855, 1622, 1582, 1530, 1444, 1299, 1264. **HRMS (ESI)**: *m/z* = 648.3747 $[M+H]^+$ (calc. for $C_{36}H_{50}N_5O_6$ *m/z* = 648.3756).

Synthesis of **SI-10**

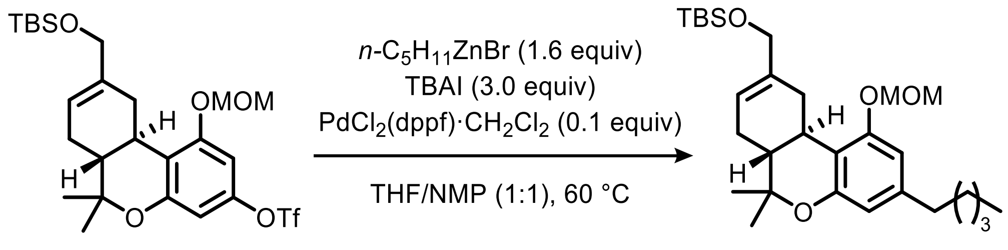

**SI-7** **SI-10**

A mixture of **SI-10** (0.10 g, 0.18 mmol, 1.0 equiv), TBAI (0.13 g, 0.35 mmol, 2.0 equiv), and $PdCl_2(dppf){\cdot}CH_2Cl_2$ (0.014 g, 0.018 mmol, 0.1 equiv) under $N_2$ was dissolved in 1:1 THF/NMP (0.6 mL). *n*-$C_5H_{11}ZnBr$ (0.5 M in THF, 0.42 mL, 0.21 mmol, 1.2 equiv) was added at ambient temperature and the resulting brown solution was heated to 60 °C and stirred for 4 h, after which TLC (5% $Et_2O$ in hexanes) showed full consumption of starting material. The reaction mixture was cooled to ambient temperature, and $CH_2Cl_2$ (1 mL) and aq. sat. $NH_4Cl$ (1 mL) were added. The aqueous phase was extracted with $CH_2Cl_2$ and the combined organic extracts were washed with brine, dried over $Na_2SO_4$, and concentrated under reduced pressure. Purification by flash column chromatography ($SiO_2$; using 2 to 5% $Et_2O$ in hexanes) afforded the product **SI-10** as a yellow oil (0.086 g, quantitative).

**$^1$H NMR** (400 MHz, $CDCl_3$) δ = 6.48 (d, *J* = 1.6 Hz, 1H), 6.36 (d, *J* = 1.6 Hz, 1H), 5.73 – 5.68 (m, 1H), 5.17 (s, 2H), 4.05 (s, 2H), 3.48 (s, 3H), 3.31 (dd, *J* = 17.1, 4.4 Hz, 1H), 2.70 (td, *J* = 10.8, 4.6 Hz, 1H), 2.52 – 2.47 (m, 2H), 2.21 (dd, *J* = 13.2, 4.8 Hz, 1H), 1.91 – 1.74 (m, 3H), 1.64 – 1.54 (m, 2H), 1.38 (s, 3H), 1.34 – 1.30 (m, 2H), 1.29 – 1.24 (m, 2H), 1.10 (s, 3H), 0.92 (s, 9H), 0.91 – 0.86 (m, 3H), 0.08 (s, 3H), 0.08 (s, 3H). **$^{13}$C NMR** (101 MHz, $CDCl_3$) δ = 156.9, 154.5, 142.9, 138.3, 119.9, 112.6, 111.3, 106.5, 94.6, 76.5, 67.2, 56.3, 45.5, 36.0, 31.9, 31.9, 31.8, 31.7, 30.9, 27.8, 27.7, 26.1, 22.7, 18.6, 14.2, -5.1, -5.1. **IR** (neat, $\nu_{max}$/cm$^{-1}$) 2954, 2928, 2894, 2856, 1617, 1575, 1403, 1254, 1154, 1112, 1066, 1050, 897. **HRMS (ESI)**: *m/z* = 489.3395 $[M+H]^+$ (calc. for $C_{29}H_{49}O_4Si$ *m/z* = 489.3395). **$[\alpha]^{25}_D$** = –99.240 ± 0.050 (c = 2.0, $CHCl_3$).

Synthesis of **SI-11**

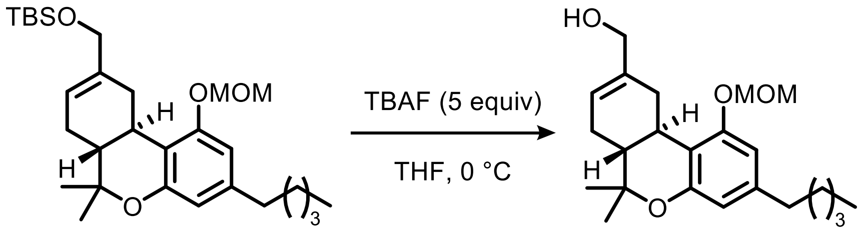


To a solution of **SI-10** (0.040 g, 0.082 mmol, 1.0 equiv) in THF (0.3 mL) at 0 °C was added TBAF (1 M in THF, 0.41 mL, 0.41 mmol, 5.0 equiv) and the solution was stirred at 0 °C for 1 h, after which TLC (30% EtOAc in hexanes) showed full consumption of starting material. The reaction was stopped by addition of aq. sat. $NaHCO_3$ and the aqueous phase was extracted with $CH_2Cl_2$. The combined organic extracts were washed with brine, dried over $Na_2SO_4$, and concentrated under reduced pressure. Purification by flash column chromatography ($SiO_2$; using 20 to 30% EtOAc in hexanes) afforded the product **SI-11** as yellow oil (0.020 g, 65%). **$^1$H NMR** (400 MHz, $CDCl_3$) δ = 6.44 (d, *J* = 1.6 Hz, 1H), 6.35 (d, *J* = 1.6 Hz, 1H), 5.77 – 5.71 (m, 1H), 5.17 (d, *J* = 1.3 Hz, 2H), 4.08 – 3.97 (m, 2H), 3.49 (s, 3H), 3.38 – 3.29 (m, 1H), 2.70 (td, *J* = 11.1, 4.8 Hz, 1H), 2.52 – 2.43 (m, 2H), 2.26 – 2.17 (m, 1H), 1.91 – 1.80 (m, 3H), 1.63 – 1.53 (m, 2H), 1.38 (s, 3H), 1.34 – 1.29 (m, 2H), 1.28 – 1.23 (m, 2H), 1.10 (s, 3H), 0.88 (m, 3H). **IR** (neat, $\nu_{max}$/cm$^{-1}$) 3416, 2954, 2928, 2856, 1617, 1574, 1429, 1404, 1182, 1151, 1049, 1010. **HRMS (ESI)**: *m/z* = 397.2349 [M+Na]$^+$ (calc. for $C_{23}H_{34}NaO_4$ *m/z* = 397.2349).

Synthesis of **SI-12**

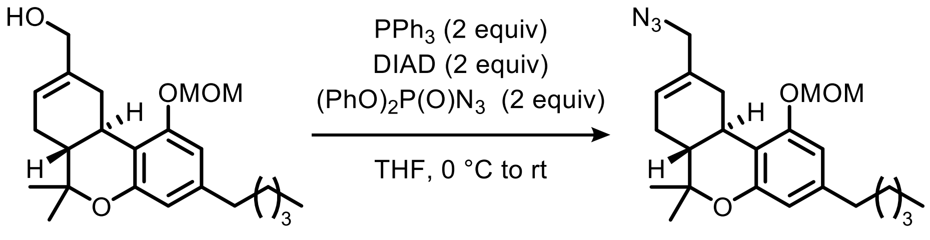


**SI-11** **SI-12**

To a solution of $PPh_3$ (0.028 g, 0.107 mmol, 2.0 equiv) in THF (0.35 mL) at 0 °C was added DIAD (0.021 mL, 0.107 mmol, 2.0 equiv) and the yellow suspension was stirred at 0 °C for 10 min. A solution of **SI-11** (0.020 g, 0.053 mmol) in THF (0.18 mL) was added, then the mixture was removed from the ice bath and $(PhO)_2P(O)N_3$ (0.023 mL, 0.107 mmol, 2.0 equiv) was added immediately. The resulting mixture was stirred at ambient temperature for 60 min. When TLC (10% EtOAc in hexanes) showed full consumption of starting material, the reaction mixture was concentrated under reduced pressure. Purification by flash column chromatography ($SiO_2$; using hexanes to 5% $Et_2O$ in hexanes) afforded the product **SI-12** as a yellow oil (0.016 g, 75%).

**$^1$H NMR** (400 MHz, $CDCl_3$) δ = 6.46 (d, *J* = 1.6 Hz, 1H), 6.36 (d, *J* = 1.6 Hz, 1H), 5.80 – 5.74 (m, 1H), 5.21 – 5.15 (m, 2H), 3.78 – 3.63 (m, 2H), 3.50 (s, 3H), 3.43 – 3.32 (m, 1H), 2.73 (td, *J* = 10.9, 4.6 Hz, 1H), 2.53 – 2.45 (m, 2H), 2.32 – 2.21 (m, 1H), 1.98 – 1.78 (m, 3H), 1.63 – 1.53 (m, 2H), 1.39 (s, 3H), 1.35 – 1.27 (m, 4H), 1.11 (s, 3H), 0.92 – 0.86 (m, 3H). **$^{13}$C NMR** (101 MHz, $CDCl_3$) δ = 156.7, 154.4, 143.2, 133.6, 125.1, 112.0, 111.3, 106.4, 94.6, 76.4, 57.4, 56.4, 45.0, 36.0, 33.1, 31.8, 31.8, 30.9, 27.9, 27.7, 22.7, 18.5, 14.2.**IR** (neat, $\nu_{max}$/cm$^{-1}$) 2956, 2929, 2857, 2096, 1617, 1575, 1237, 1151. **HRMS (ESI)**: *m/z* = 422.2413 [M+Na]$^+$ (calc. for $C_{23}H_{33}NaN_3O_3$ *m/z* = 422.2414).

Synthesis of **3**

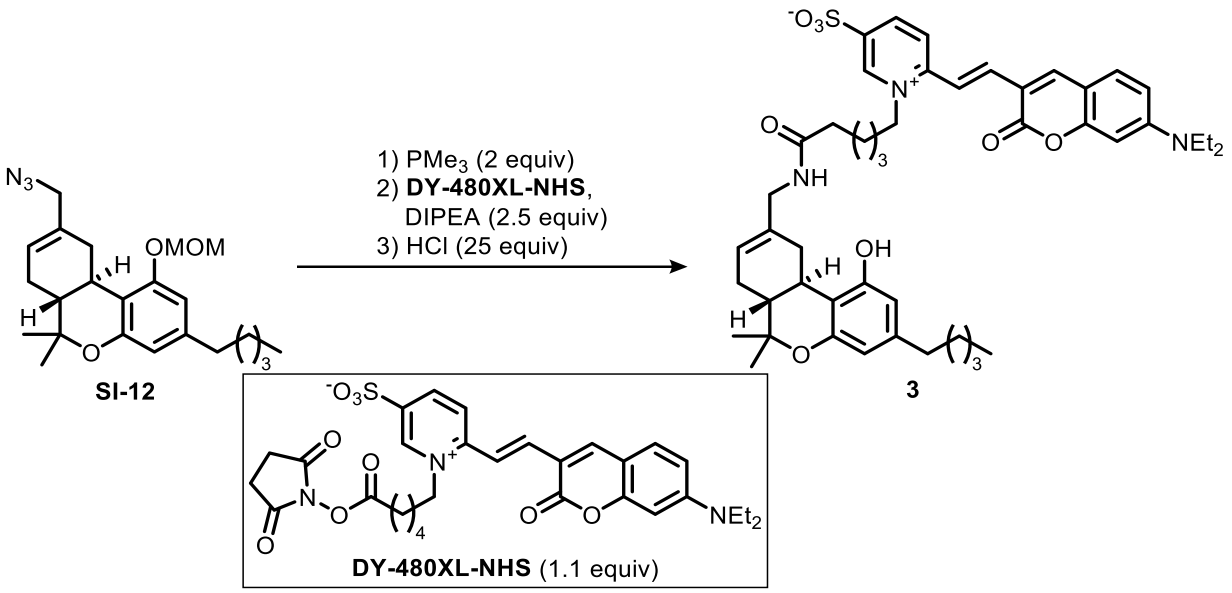

To a solution of **SI-12** (3.5 mg, 8.8 µmol, 1.0 equiv.) in 10:1 THF/water (0.1 mL) was added $PMe_3$ (1 M in THF, 0.018 mL, 0.018 mmol, 2.0 equiv.) and the reaction was stirred at ambient temperature for 4 h, after which TLC (10% EtOAc in hexanes) showed full consumption of starting material. The reaction mixture was concentrated under reduced pressure, and the residue was taken up in $CH_2Cl_2$ (0.1 mL). **DY-480XL-NHS** (6.0 mg, 9.7 µmol, 1.1 equiv.) and DIPEA (8.8 µL, 0.022 mmol, 2.5 equiv.) were added and the reaction was stirred overnight at ambient temperature. The mixture was concentrated under reduced pressure, the residue was dissolved in MeOH (0.1 mL) and HCl (2 M in $Et_2O$, 0.10 mL, 0.20 mmol, 25 equiv) was added. The mixture was stirred at ambient temperature for 2 h, then the reaction was concentrated *in vacuo* and directly purified by preparative TLC ($SiO_2$; using 5% MeOH in DCM) afforded the product **3** as a red solid (4.6 mg, 63% over three steps).

**$^1$H NMR** (500 MHz, $CD_2Cl_2$) δ 9.14 (s, 1H), 8.59 (d, $J$ = 8.5 Hz, 1H), 8.06 (d, $J$ = 8.9 Hz, 1H), 8.02 (d, $J$ = 15.4 Hz, 1H), 7.85 (s, 1H), 7.45 – 7.35 (m, 2H), 6.70 – 6.66 (m, 1H), 6.63 (bs, 1H), 6.54 – 6.45 (m, 1H), 6.22 (d, $J$ = 1.7 Hz, 1H), 6.07 (d, $J$ = 1.6 Hz, 1H), 5.62 (bs, 1H), 4.67 – 4.53 (m, 2H), 3.82 – 3.68 (m, 2H), 3.51 – 3.35 (m, 5H), 2.58 (td, $J$ = 10.7, 4.2 Hz, 1H), 2.37 – 1.96 (m, 10H), 1.85 – 1.39 (m, 6H), 1.37 – 1.14 (m, 13H), 0.99 (s, 3H), 0.82 (t, $J$ = 7.0 Hz, 3H). **$^{13}$C NMR** (126 MHz, $CD_2Cl_2$) δ 173.4, 160.9, 157.5, 156.9, 155.1, 153.5, 148.1, 143.1, 136.0, 131.5, 124.8, 122.6, 116.1, 113.9, 110.9, 110.8, 109.5, 109.2, 108.5, 97.2, 76.6, 59.4, 46.0, 45.8, 45.6, 36.2, 36.1, 32.7, 32.2, 32.1, 31.4, 29.4, 28.2, 27.9, 25.9, 25.1, 23.1, 18.7, 14.4, 12.8. **IR** (neat, $\nu_{max}$/cm$^{-1}$) 3331, 2927, 2856, 1713, 1618, 1579, 1504, 1042.

Synthesis of **4**

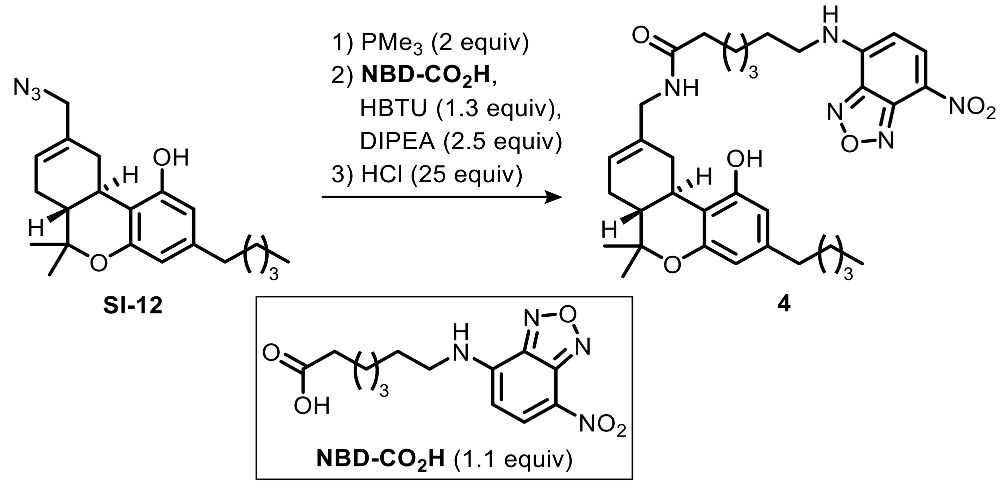


To a solution of **SI-12** (3.5 mg, 8.8 µmol, 1.0 equiv) in 10:1 THF/water (0.1 mL) was added $PMe_3$ (1 M in THF, 0.018 mL, 0.018 mmol, 2.0 equiv.) and the reaction was stirred at ambient temperature for 4 h, after which TLC (10% EtOAc in hexanes) showed full consumption of starting material. The reaction mixture was concentrated under reduced pressure, and the residue was taken up in $CH_2Cl_2$ (0.1 mL). Acid **NBD-CO₂H** (3.1 mg, 9.1 µmol, 1.1 equiv.), HBTU (5.4 mg, 0.010 mmol, 1.3 equiv.), DIPEA (3.6 µL, 0.021 mmol, 2.5 equiv.) were added and after stirring for 1 h full conversion was observed by TLC (5% MeOH in DCM, 1% 7 N $NH_3$ in MeOH). Aq. sat. $NaHCO_3$ was added, and the aqueous layer was extracted with $CH_2Cl_2$. The combined organic extracts were washed with brine, dried over $Na_2SO_4$ and concentrated under reduced pressure. Finally, the residue was dissolved in MeOH (0.1 mL) and HCl (2 M in $Et_2O$, 0.10 mL, 0.20 mmol, 25 equiv) was added. The mixture was stirred at ambient temperature for 2h, then the reaction was stopped by addition of aq. sat. $NaHCO_3$. The aqueous layer was extracted with $CH_2Cl_2$ and the combined organic extracts were washed with brine, dried over $Na_2SO_4$ and concentrated under reduced pressure. Purification by preparative TLC ($SiO_2$; using 60% EtOAc in hexanes) afforded the product as an orange solid (4.0 mg, 73% over three steps).

**$^1$H NMR** (500 MHz, $CD_2Cl_2$) δ = 8.47 (dd, *J* = 8.7, 0.5 Hz, 1H), 6.61 (br, 1H), 6.26 – 6.13 (m, 3H), 6.02 (br, 1H), 5.69 – 5.56 (m, 2H), 3.87 – 3.72 (m, 2H), 3.53 – 3.41 (m, 2H), 3.31 (dd, *J* = 16.0, 4.6 Hz, 1H), 2.66 (td, *J* = 11.1, 4.8 Hz, 1H), 2.39 (td, *J* = 7.4, 1.9 Hz, 2H), 2.27 – 2.13 (m, 3H), 1.85 – 1.63 (m, 5H), 1.59 – 1.24 (m, 12H), 1.33 (s, 3H), 1.06 (s, 3H), 0.87 (t, *J* = 7.0 Hz, 3H). **$^{13}$C NMR** (126 MHz, $CD_2Cl_2$) δ = 173.6, 156.0, 155.3, 145.0, 143.4, 137.3, 135.9, 121.8, 110.6, 110.1, 108.2, 76.8, 45.6, 44.5, 37.0, 36.0, 33.2, 32.1, 32.0, 31.4, 29.2, 28.7, 28.2, 27.9, 27.1, 25.9, 23.1, 18.7, 14.4. **IR** (neat, $\nu_{max}$/cm$^{-1}$) 3314, 2926, 2855, 1621, 1582, 1530,

1443, 1426, 1351, 1298, 1264, 1185. **HRMS (ESI)**: *m/z* = 620.3440 $[M+H]^+$ (calc. for $C_{34}H_{46}N_5O_6$ *m/z* = 620.3443). $\mathbf{[\alpha]^{25}_D}$ = –86.767 ± 0.577 (c = 0.20, $CHCl_3$).

Synthesis of **2**

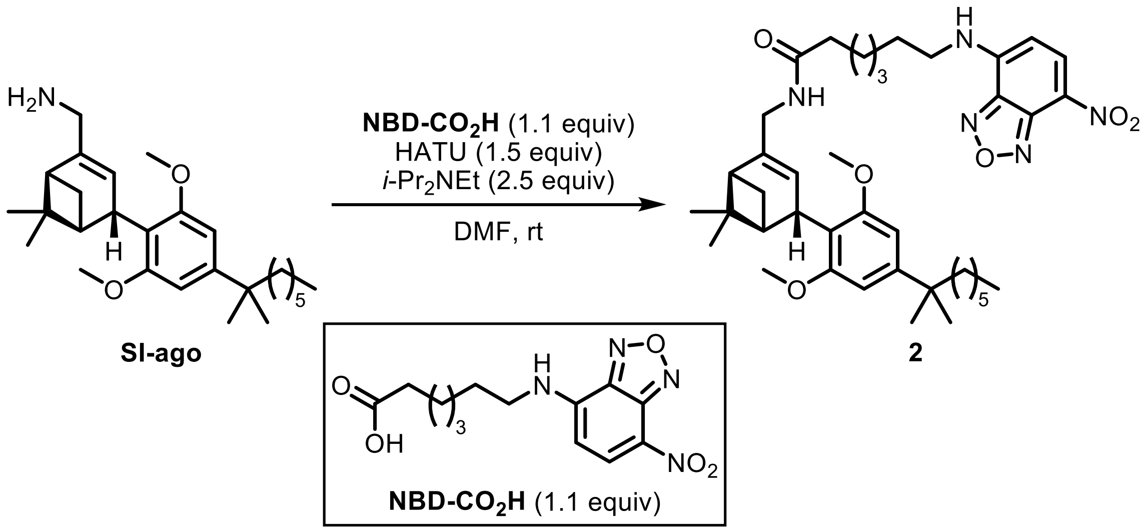


Synthesis of **SI-ago** can be found in *Chem. Eur. J.* **2020**, *26*, 1380–1387.

To a solution of **NBD-CO₂H** (4.2 mg, 13.5 µmol, 1.1 equiv) in DMF (0.1 mL) was added HATU (7.0 mg, 18.4 µmol, 1.5 equiv) and *i*-$Pr_2NEt$ (5 µL, 30.8 µmol, 2.5 equiv) and the reaction mixture was stirred at 0 °C for 5 min before being added to a mixture of **SI-ago** (5.1 mg, 12.3 µmol, 1.0 equiv) in DMF (50 µL). The reaction mixture was stirred at rt for 1 h and concentrated *in vacuo*. The crude product was purified by preparative TLC ($SiO_2$, 50% EtOAc in hexanes) to afford the title product **2** as an orange solid (7.6 mg, 88%).

**$^1$H NMR** (400 MHz, $CD_2Cl_2$) δ 8.47 (d, *J* = 8.7 Hz, 1H), 6.49 (s, 2H), 6.19 (d, *J* = 8.7 Hz, 1H), 5.59 (dt, *J* = 2.9, 1.5 Hz, 1H), 5.39 (t, *J* = 5.7 Hz, 1H), 3.98 – 3.93 (m, 1H), 3.90 – 3.76 (m, 2H), 3.72 (s, 6H), 3.53 – 3.44 (m, 2H), 2.23 – 2.12 (m, 3H), 2.08 (td, *J* = 5.7, 1.4 Hz, 1H), 2.04 – 1.97 (m, 1H), 1.87 – 1.74 (m, 2H), 1.71 – 1.62 (m, 3H), 1.61 – 1.39 (m, 6H), 1.32 – 1.20 (m, 8H), 1.27 (s, 3H), 1.25 (s, 6H), 1.15 – 1.03 (m, 2H), 0.95 (s, 3H), 0.85 (t, *J* = 6.7 Hz, 2H). **$^{13}$C NMR** (101 MHz, $CD_2Cl_2$) δ 172.29, 158.49, 149.71, 144.53, 144.16, 138.68, 136.77, 123.48, 117.41, 102.81, 55.74, 47.47, 44.46, 44.37, 44.25, 43.94, 40.72, 38.00, 37.52, 36.52, 31.85, 30.09, 29.76, 28.81, 28.67, 28.21, 27.60, 26.55, 26.08, 25.43, 24.74, 22.73, 20.79, 13.90. **IR** (neat, $\nu_{max}$/cm$^{-1}$) 3312, 2926, 2856, 1649, 1621, 1580, 1530, 1502, 1448, 1410, 1350, 1299, 1272, 1239, 1186, 1162, 1123, 1040. **HRMS (ESI)**: *m/z* = 704.4374 $[M+H]^+$ (calc. for $C_{40}H_{58}N_5O_6$ *m/z* = 704.4382). **$[\alpha]^{25}_D$** = +38.271 ± 0.471 (c = 0.31, $CH_2Cl_2$).

Synthesis of **SI-15**

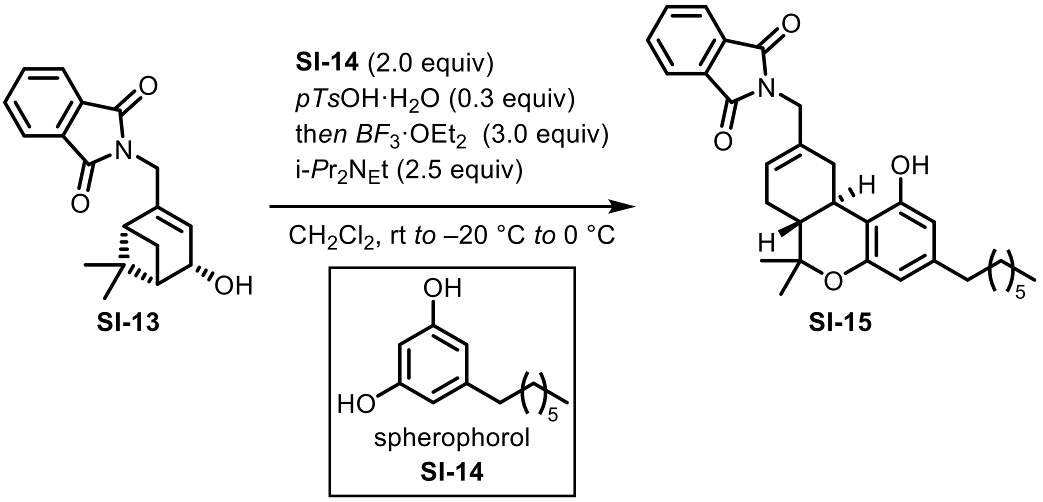


Verbenol **SI-13** (149 mg, 500 µmol, 1.0 equiv) was added to a solution of spherophorol **SI-14** (208 mg, 1.00 mmol, 2.0 equiv) and TsOH·$H_2O$ (28.5 mg, 150 µmol, 0.3 equiv) in $CH_2Cl_2$ (10 mL), and the mixture was stirred for 1 h at rt. The reaction mixture was cooled to –20 °C. $BF_3$·$OEt_2$ (190 µmol, 1.54 mmol, 3.0 equiv) was added dropwise, and the resulting pale green solution was allowed to warm to 0 °C over the course of 2 h while stirring. The reaction was quenched by addition of brine (25 mL). The phases were separated, and the aqueous layer was extracted with $Et_2O$ (4 × 20 mL). Combined organic extracts were dried over $MgSO_4$, filtered and concentrated *in vacuo*. Purification by flash column chromatography ($SiO_2$; 5 – 15% Acetone in hexanes) afforded the product **SI-15** as an off-white foam (136 mg, 56%).

**$^1$H NMR** (400 MHz, $CDCl_3$) δ 7.91 – 7.82 (m, 2H), 7.77 – 7.67 (m, 2H), 6.24 (s, 2H), 5.98 – 5.92 (m, 1H), 4.44 (ddd, *J* = 15.5, 3.0, 1.9 Hz, 1H), 4.27 – 4.17 (m, 1H), 4.04 – 3.97 (m, 1H), 2.44 – 2.36 (m, 2H), 2.34 – 2.19 (m, 3H), 1.57 – 1.48 (m, 2H), 1.47 (d, *J* = 9.1 Hz, 1H), 1.29 (s, 3H), 1.29 – 1.21 (m, 8H), 0.97 (s, 3H), 0.86 (t, *J* = 6.9 Hz, 3H). **$^{13}$C NMR** (101 MHz, $CDCl_3$) δ 168.5, 155.4, 148.0, 143.3, 134.3, 132.0, 123.7, 120.6, 111.9, 108.7, 47.4, 44.3, 43.2, 41.2, 37.8, 35.6, 31.9, 31.0, 29.4, 29.3, 28.1, 26.0, 22.8, 20.7, 14.2. **IR** (neat, $\nu_{max}$/cm$^{-1}$): 3449, 2925, 2855, 1770, 1708, 1624, 1578, 1467, 1430, 1392, 1341, 1282, 1111, 1019, 947, 756, 728. **HRMS (ESI)**: *m/z* = 488.279 [M+H]$^+$ (calc. for $C_{31}H_{38}NO_4$ *m/z* = 488.2795). **[α]$^{25}_D$** = –98.221 ± 0.098 (c = 1.0, $CHCl_3$).

Synthesis of **SI-16**

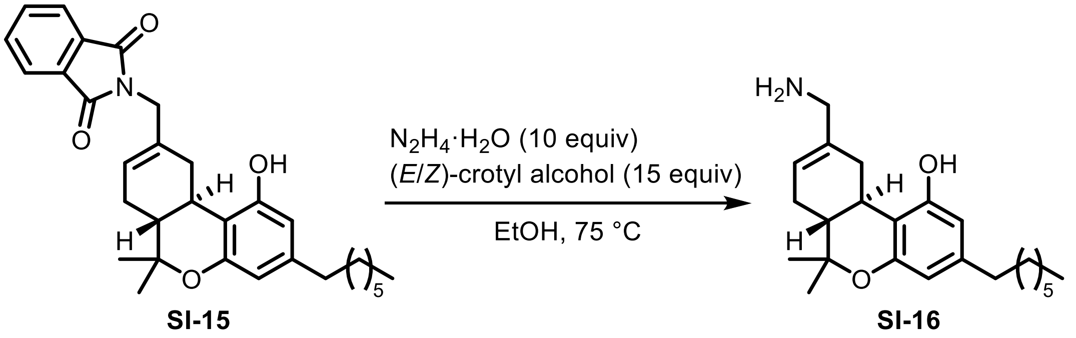

To a solution of **SI-15** (150 mg, 307 µmol, 1.0 equiv) in EtOH (6 mL) were added $N_2H_4 \cdot H_2O$ (150 µL, 3.08 mmol, 10 equiv) and (*E*/*Z*)-crotyl alcohol (400 µL, 4.69 mmol, 15 equiv), and the resulting mixture was stirred at 75 °C for 2 h. The mixture was cooled to rt, filtered through a pad of Celite® and washed with EtOH (3 × 30 mL). Purification by flash column chromatography ($SiO_2$; 1% 6M $NH_3$ in MeOH, 9% MeOH in $CH_2Cl_2$) afforded the product **SI-16** as a pink oil (82.7 mg, 75%).

**$^1$H NMR** (500 MHz, $CD_3OD$) δ 6.17 (d, *J* = 1.6 Hz, 1H), 6.09 (d, *J* = 1.6 Hz, 1H), 5.71 – 5.67 (m, 1H), 3.41 (ddd, *J* = 17.3, 4.8, 2.0 Hz, 1H), 3.25 – 3.23 (m, 2H), 2.69 – 2.62 (m, 1H), 2.43 – 2.38 (m, 2H), 2.29 – 2.20 (m, 1H), 1.95 – 1.85 (m, 1H), 1.83 – 1.76 (m, 1H), 1.75 (td, *J* = 11.7, 4.5 Hz, 1H), 1.59 – 1.51 (m, 2H), 1.35 (s, 3H), 1.34 – 1.21 (m, 8H), 1.08 (s, 3H), 0.90 (t, *J* = 7.0 Hz, 3H). **$^{13}$C NMR** (126 MHz, $CD_3OD$) δ 157.8, 155.8, 143.6, 138.2, 122.2, 111.4, 109.8, 108.5, 77.2, 47.7, 46.8, 36.6, 34.0, 33.1, 33.0, 32.3, 30.3, 30.3, 28.8, 28.0, 23.7, 18.6, 14.4. **IR** (neat, $\nu_{max}/cm^{-1}$): 2924, 2853, 1619, 1574, 1424, 1382, 1264, 1184, 1109, 1084, 1049, 839. **HRMS (ESI)**: *m/z* = 358.2736 $[M+H]^+$ (calc. for $C_{23}H_{36}NO_2$ *m/z* = 358.2741). **$[\alpha]^{25}_D$** = –143.774 ± 0.377 (c = 1.0, $CHCl_3$).

Synthesis of **1**

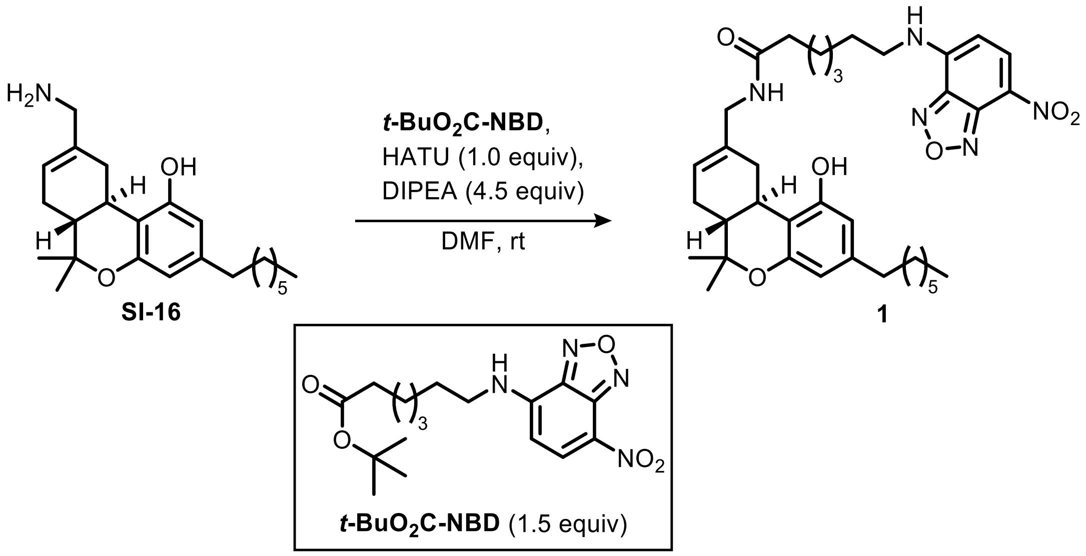

***t*-BuO$_2$C-NBD** (56.3 mg, 155 μmol, 1.5 equiv) was dissolved in TFA (1.0 mL) and the solution was stirred for 2 h at rt. The mixture was concentrated *in vacuo* and used directly in the next step without further purification. To the solution of this crude, amine **SI-16** (36.5 mg, 102 μmol, 1.0 equiv) and DIPEA (80 μL, 459 μmol, 4.5 equiv) in DMF (0.30 mL), was added HATU (39.0 mg, 103 μmol, 1.0 equiv). The reaction mixture was stirred at rt for 1 h and subsequently quenched with sat. aq. $NH_4Cl$ (20 mL). The mixture was diluted with $CH_2Cl_2$ (10 mL). The layers were separated, and the aqueous phase was extracted with EtOAc (3 × 15 mL). Combined organic extracts were dried over $MgSO_4$, filtered and concentrated *in vacuo*. Purification by flash column chromatography ($SiO_2$; 1 – 3%, MeOH in $CH_2Cl_2$) afforded the product **1** as a red wax (53.8 mg, 81%).

**$^1$H NMR** (500 MHz, $CDCl_3$) δ 8.43 (d, *J* = 8.7 Hz, 1H), 7.11 (m, 1H), 6.96 (m, 1H), 6.19 (s, 2H), 6.13 (d, *J* = 8.7 Hz, 1H), 5.86 (t, *J* = 5.9 Hz, 1H), 5.56 (d, *J* = 4.9 Hz, 1H), 3.80 (d, *J* = 5.8 Hz, 2H), 3.53 – 3.40 (m, 2H), 3.34 (dd, *J* = 15.3, 3.6 Hz, 1H), 2.63 (td, *J* = 10.9, 4.7 Hz, 1H), 2.35 (td, *J* = 7.5, 3.6 Hz, 2H), 2.27 (t, *J* = 7.2 Hz, 2H), 2.18 – 2.08 (m, 1H), 1.85 – 1.64 (m, 5H), 1.54 – 1.43 (m, 6H), 1.32 (s, 3H), 1.30 – 1.16 (m, 10H), 1.04 (s, 3H), 0.84 (t, *J* = 6.9 Hz, 3H). **$^{13}$C NMR** (126 MHz, $CDCl_3$) δ 173.8, 155.8, 154.8, 144.5, 144.4, 144.1, 142.9, 136.9, 135.2, 123.4, 121.7, 110.1, 109.5, 108.0, 98.8, 76.4, 45.5, 45.1, 44.1, 36.6, 35.7, 32.7, 31.9, 31.6, 31.1, 29.5, 29.3, 28.8, 28.2, 27.8, 27.6, 26.6, 25.5, 22.8, 18.5, 14.2. **IR** (neat, $\nu_{max}$/cm$^{-1}$): 3317, 2924, 2853, 1620, 1580, 1529, 1442, 1350, 1296, 1262, 1184. **HRMS (ESI)**: *m/z* = 670.3576 [M+Na]$^+$ (calc. for $C_{36}H_{49}N_5NaO_6$ *m/z* = 670.3575). **[α]$^{25}_{D}$** = –96.006 ± 0.321 (c = 1.0, $CHCl_3$).

### NMR SPECTRA

$^{1}$H NMR (400 MHz, $CD_3CN$) of **SI-3**

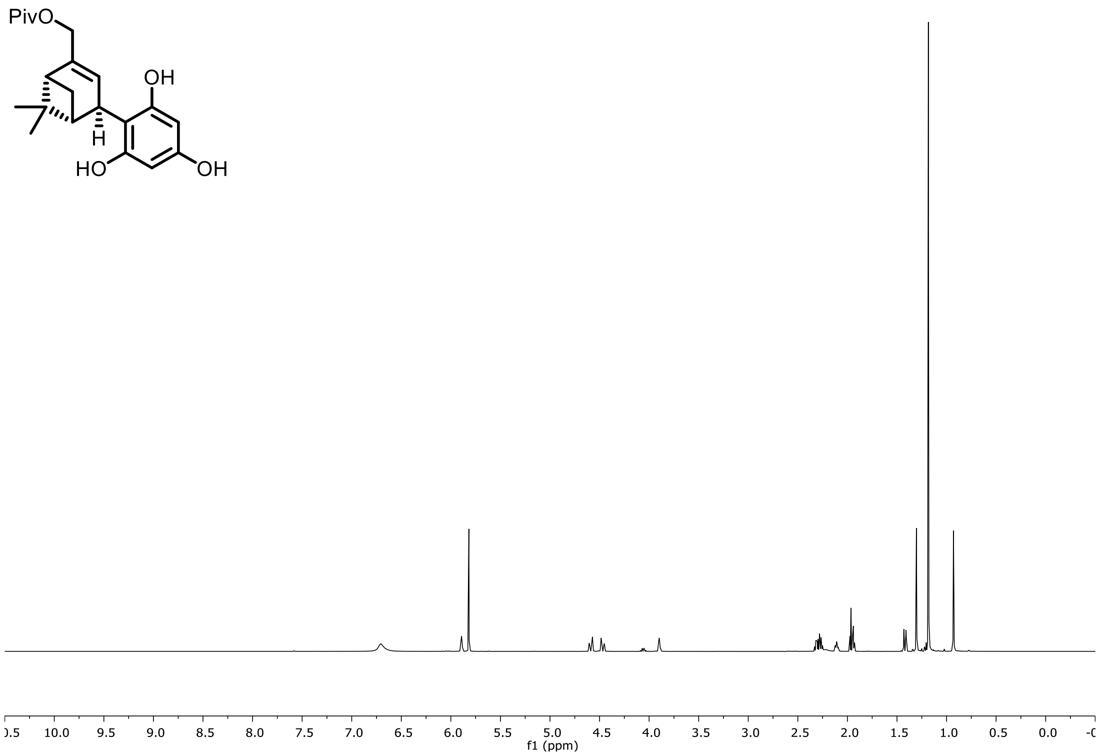


$^{13}$C NMR (101 MHz, $CD_3CN$) of **SI-3**

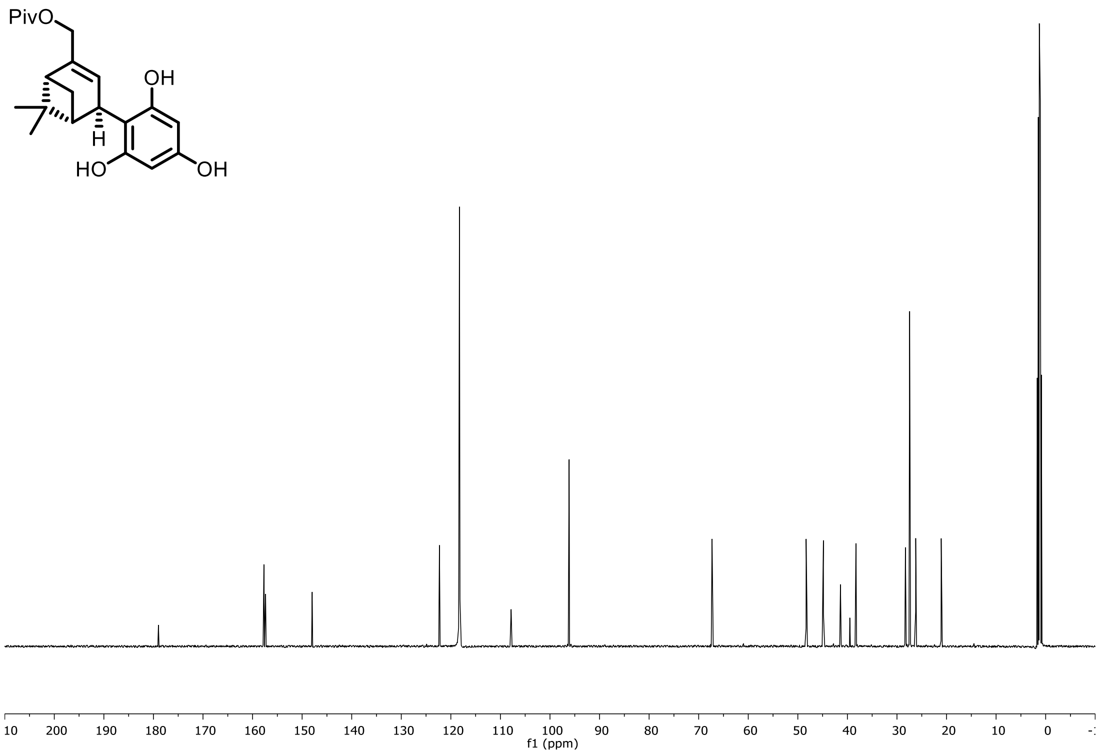

$^{1}H$ NMR (400 MHz, $CDCl_3$) of **SI-5**

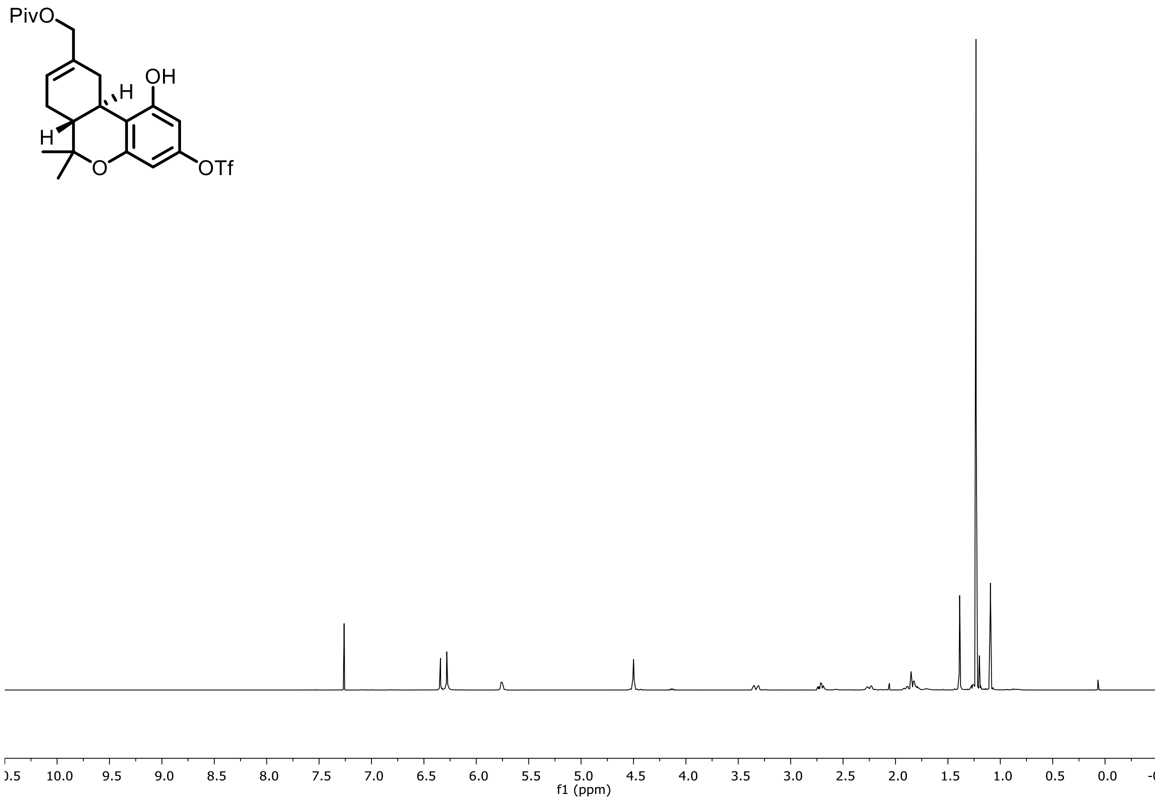


$^{13}C$ NMR (101 MHz, $CDCl_3$) of **SI-3**

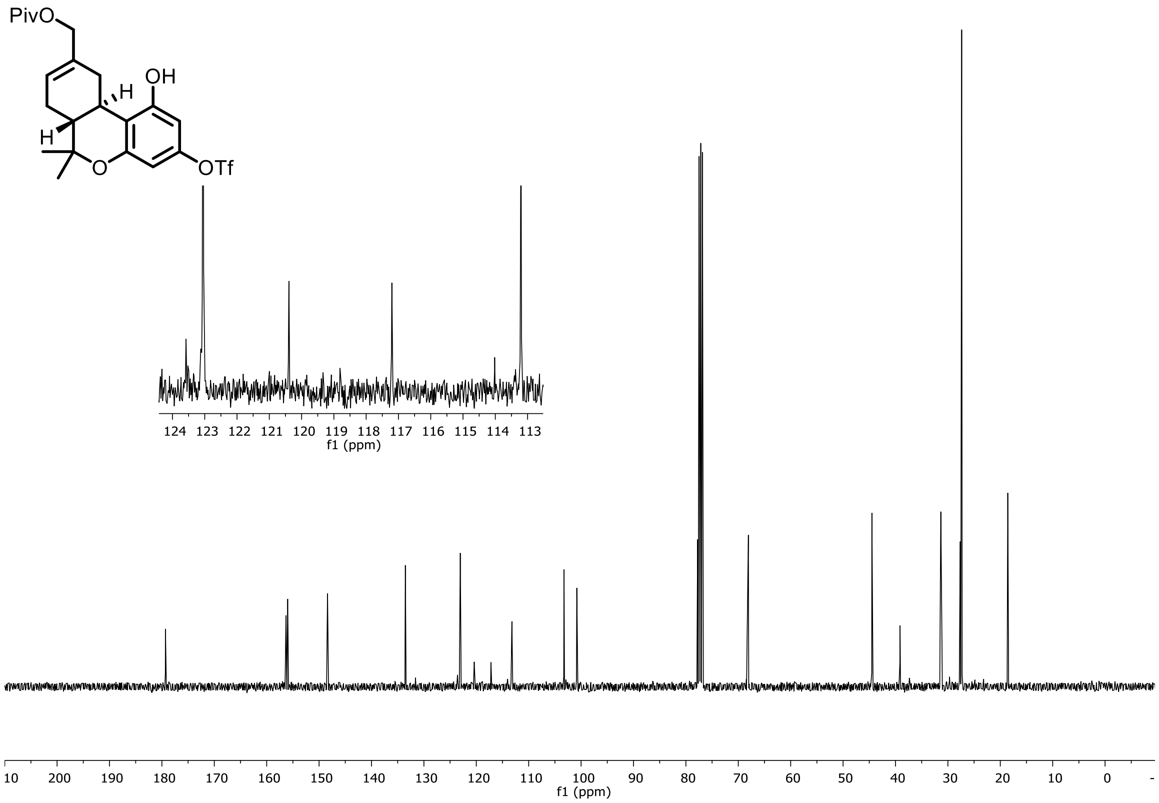

$^{19}$F NMR (376 MHz, $CDCl_3$) of **SI-5**

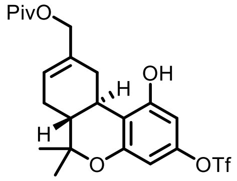


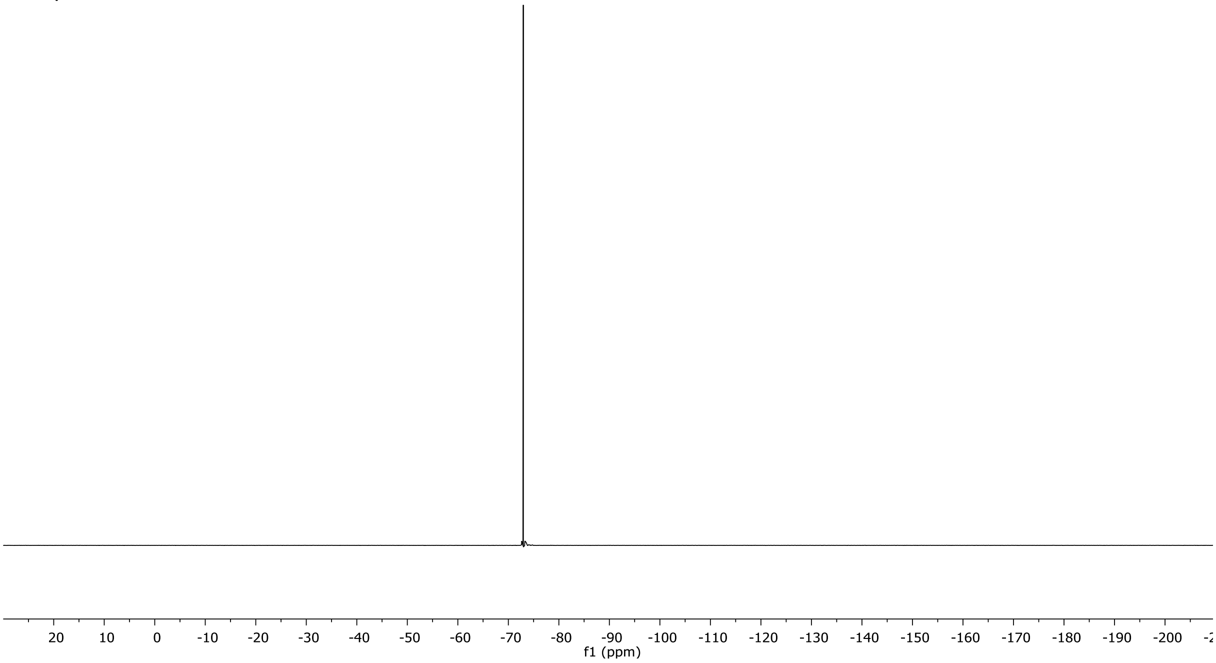

[1]H NMR (500 MHz, $CDCl_3$) of **SI-6**

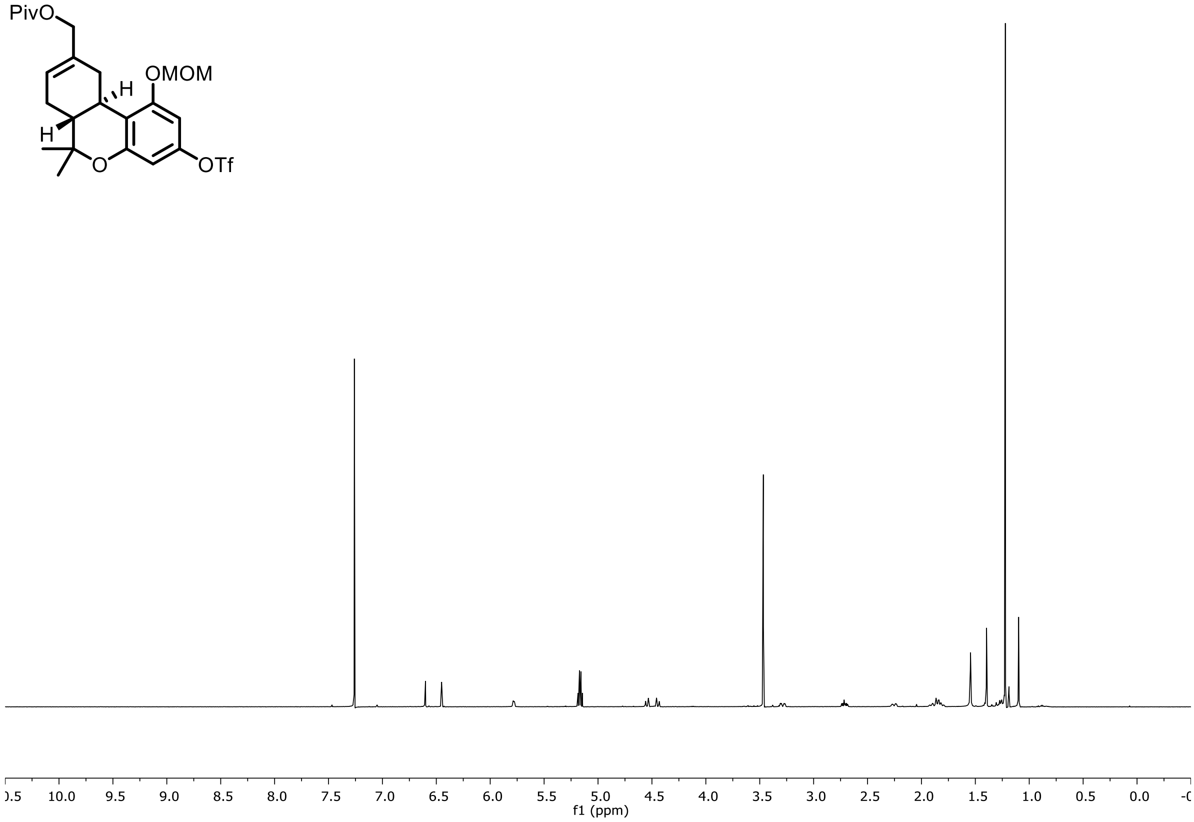


[13]C NMR (126 MHz, $CDCl_3$) of **SI-6**

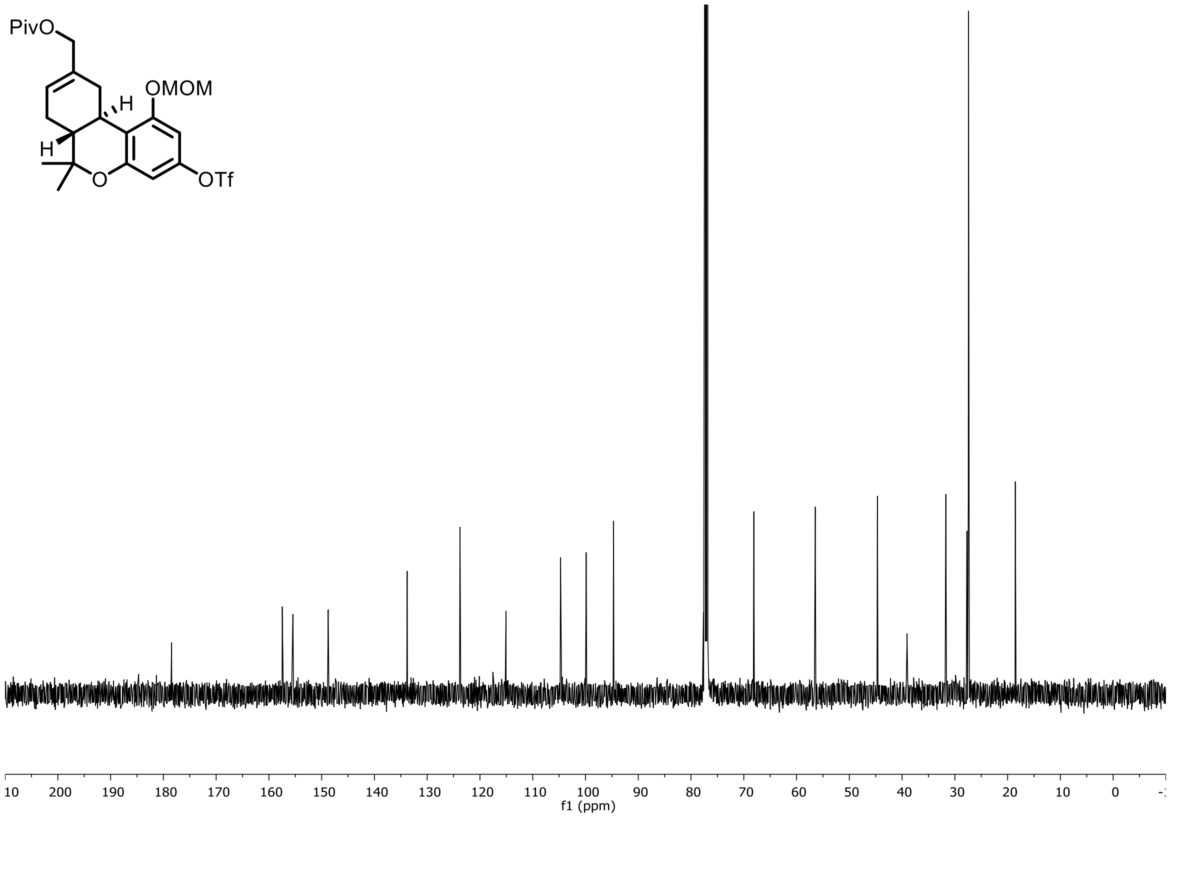

<sup></sup>

$^{19}$F NMR (471 MHz, $CDCl_3$) of **SI-6**

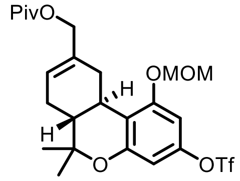


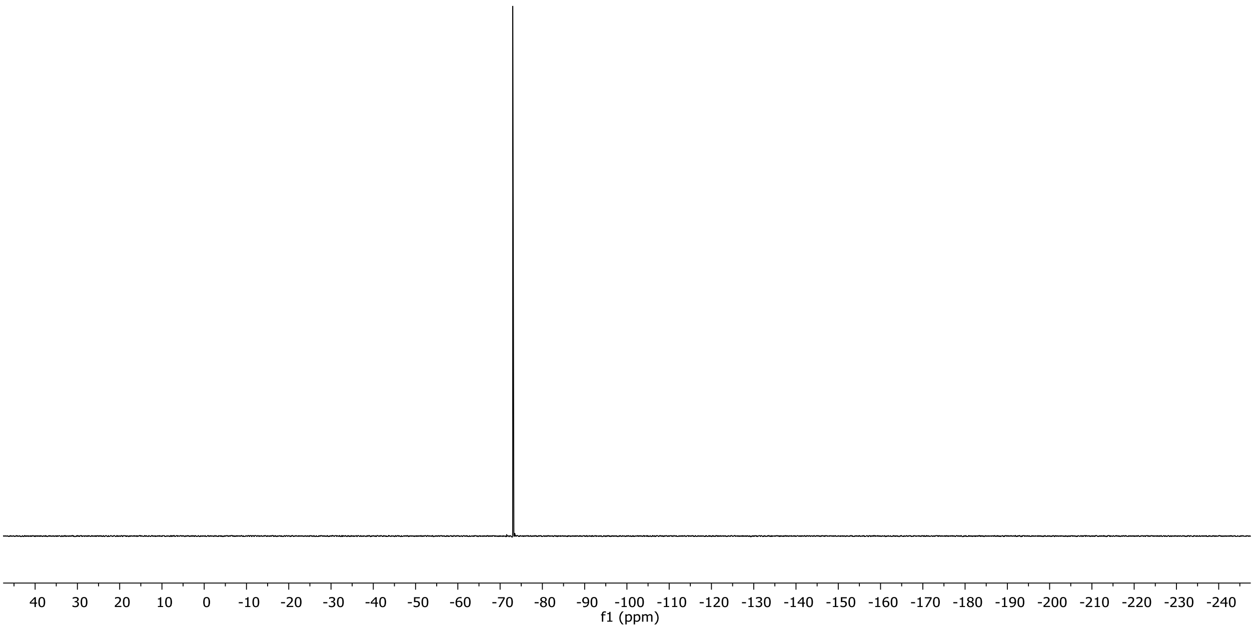

$^{1}H$ NMR (400 MHz, $CDCl_3$) of **SI-7**

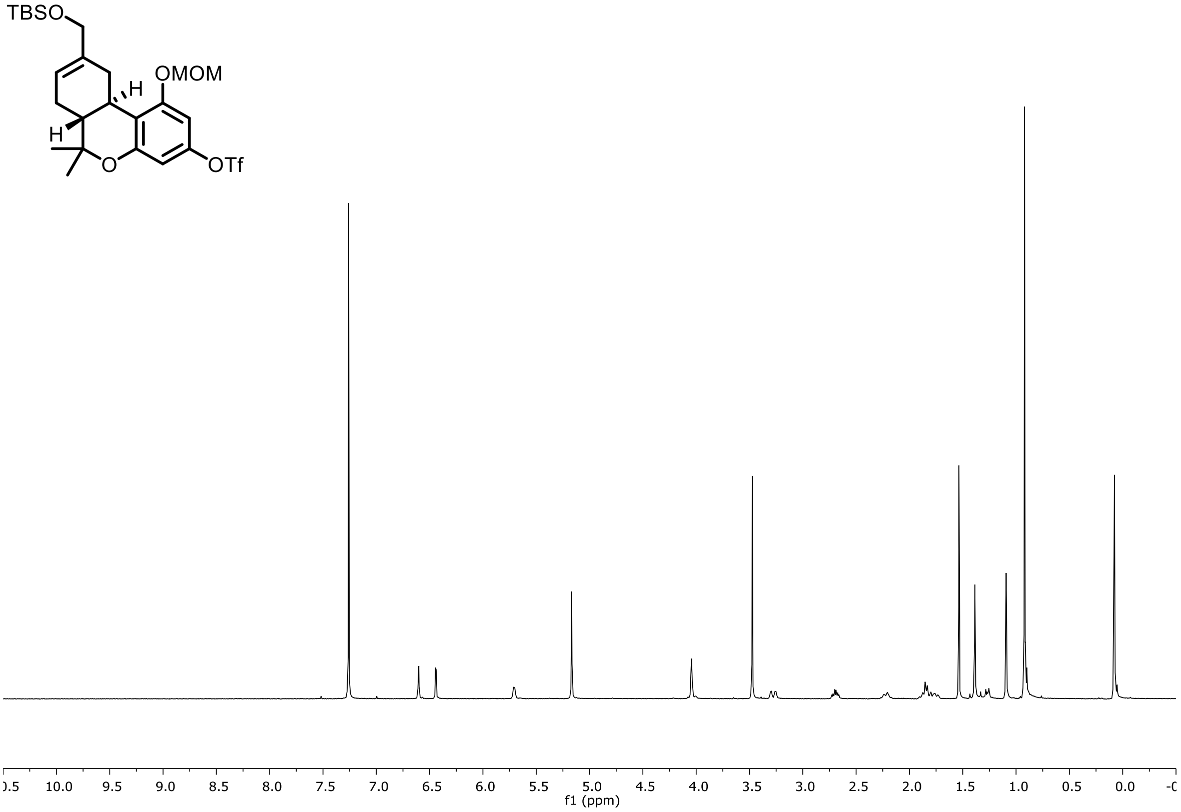


$^{13}C$ NMR (101 MHz, $CDCl_3$) of **SI-7**

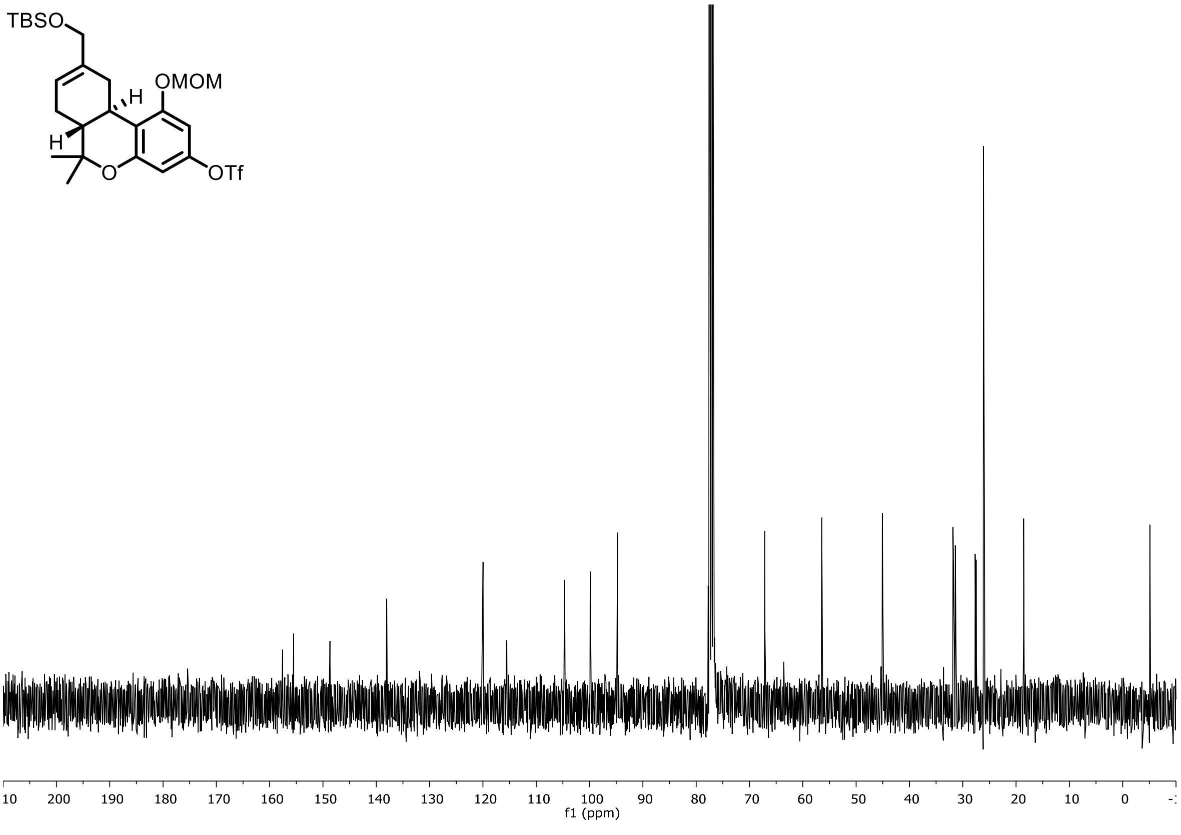

$^{19}$F NMR (376 MHz, $CDCl_3$) of **SI-7**

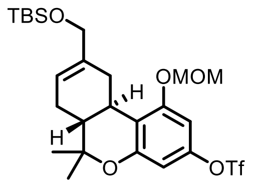


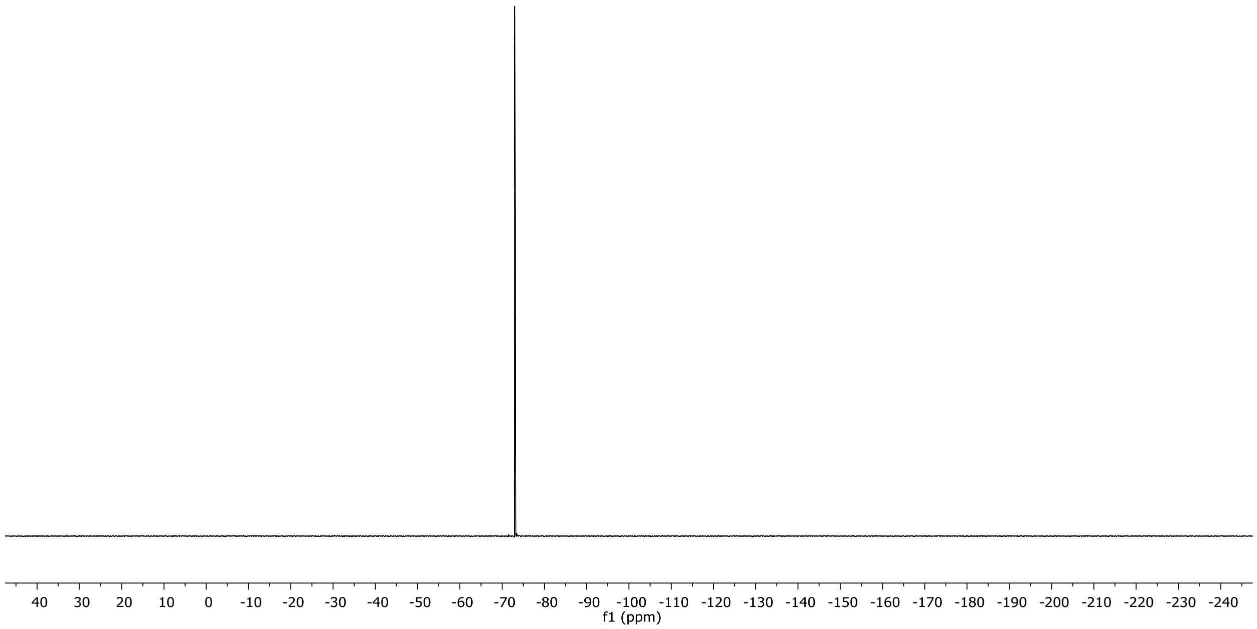

$^{1}H$ NMR (500 MHz, $CDCl_3$) of **SI-8**

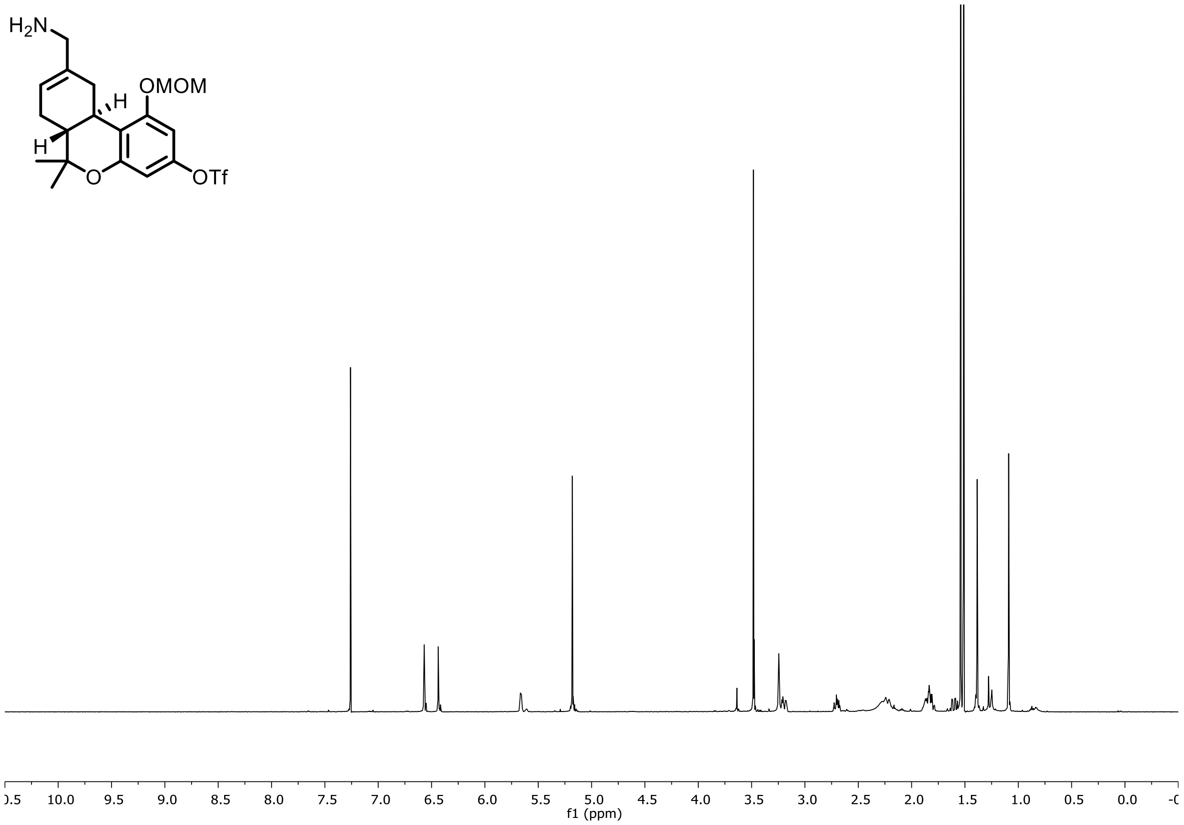


$^{13}C$ NMR (126 MHz, $CDCl_3$) of **SI-8**

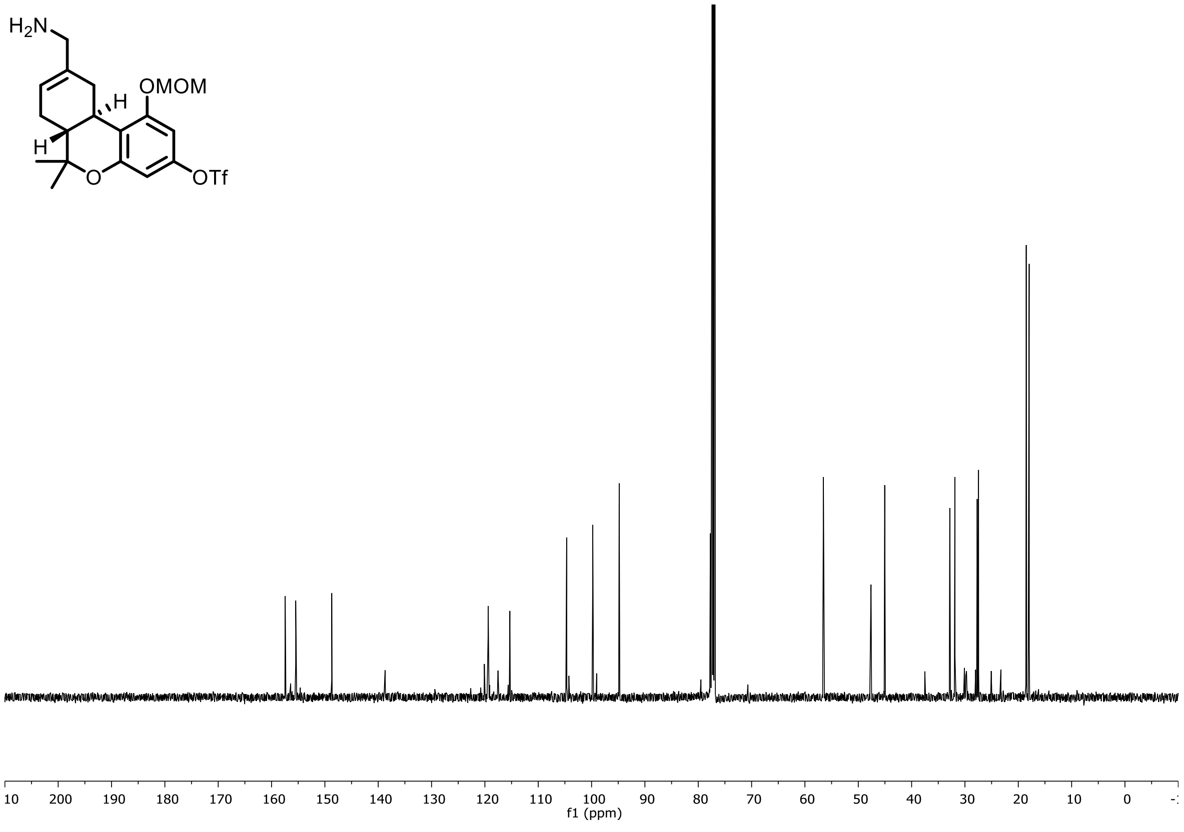

$^{19}F$ NMR (471 MHz, $CDCl_3$) of **SI-8**

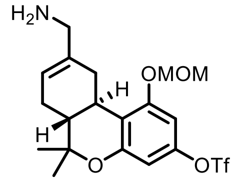


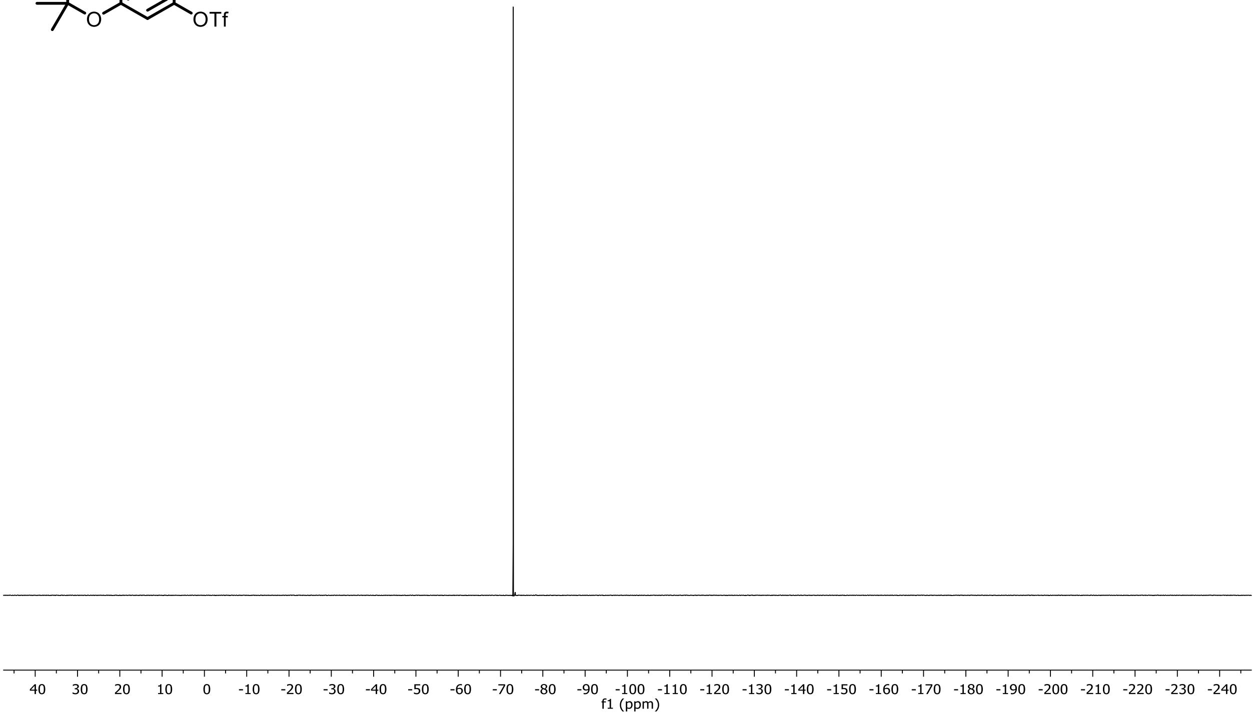

$^1$H NMR (400 MHz, $CDCl_3$) of **SI-9**

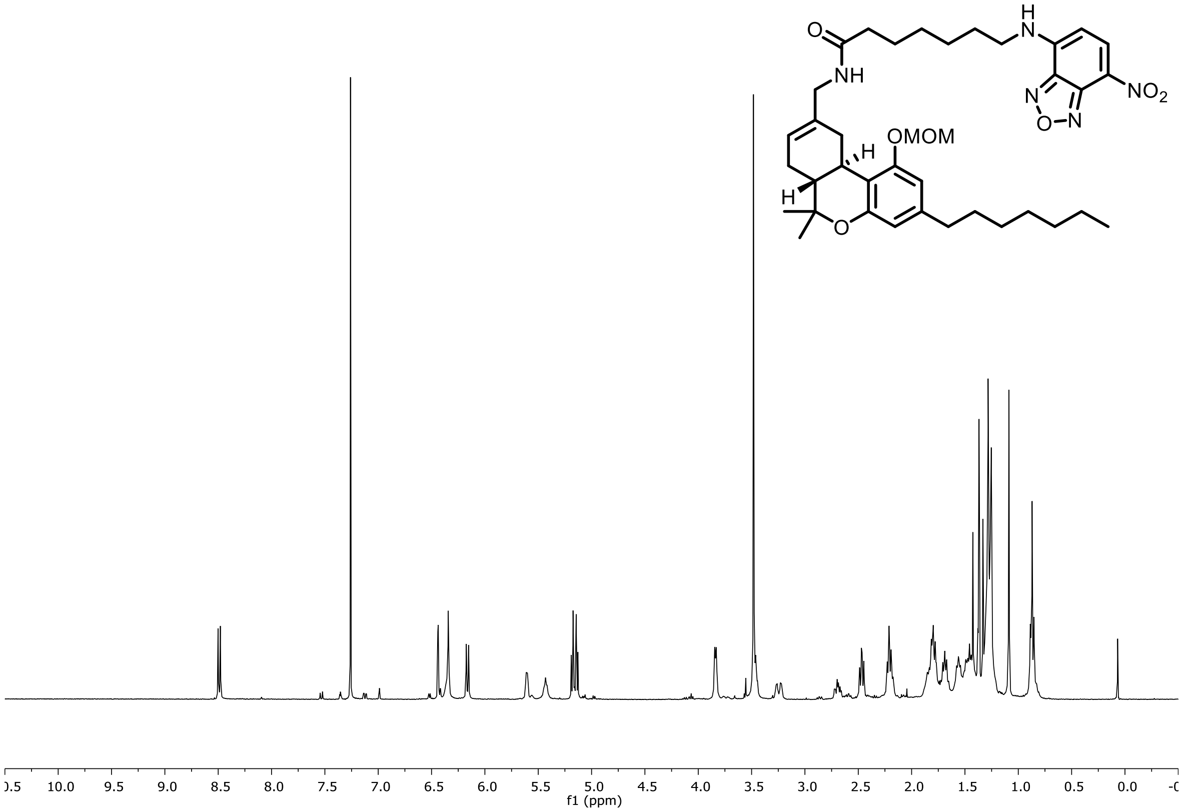


$^{13}$C NMR (101 MHz, $CDCl_3$) of **SI-9**

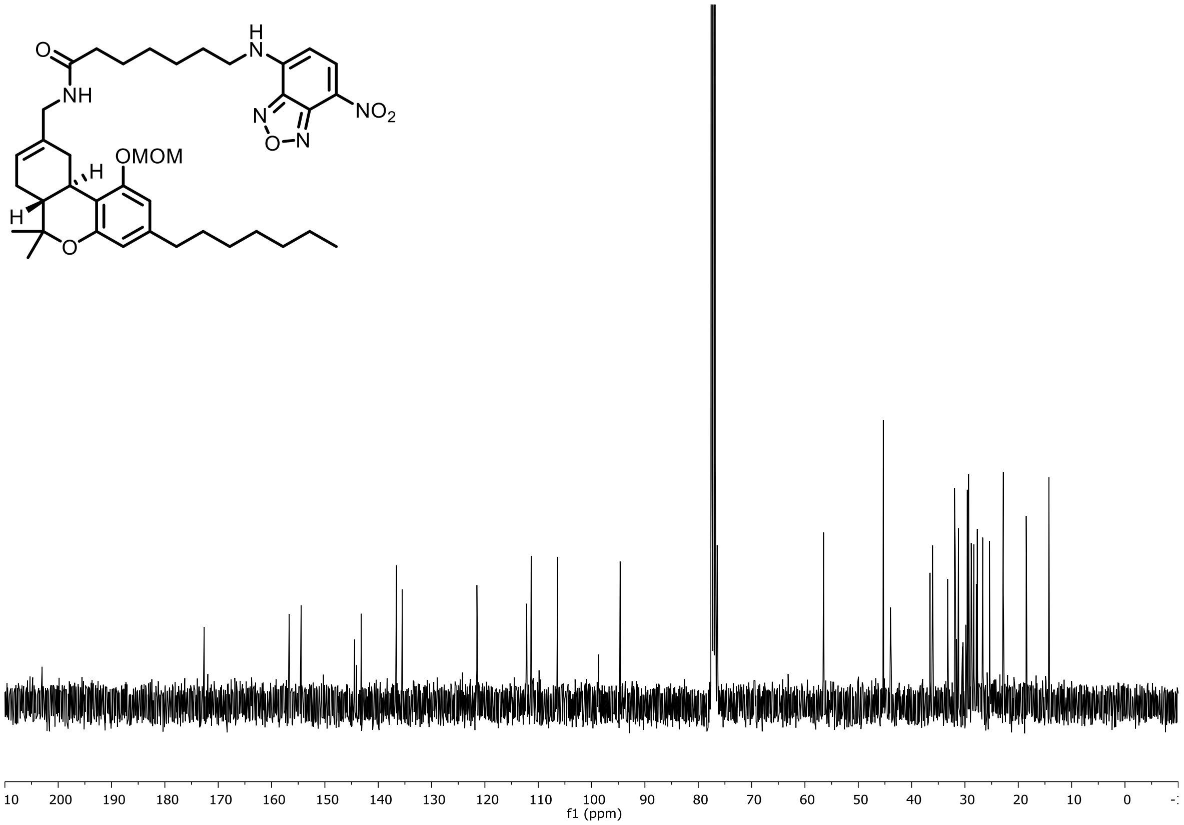

$^{1}H$ NMR (500 MHz, $CDCl_3$) of **1**

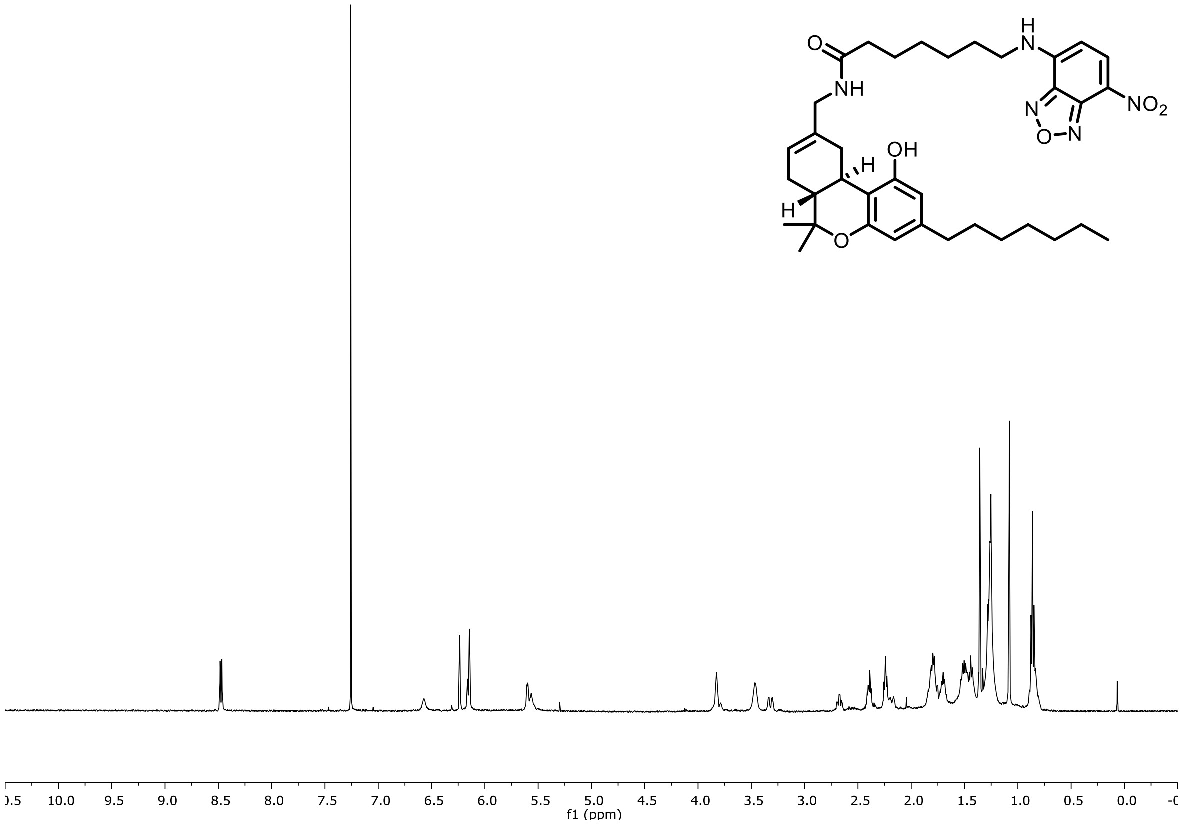


$^{13}C$ NMR (126 MHz, $CDCl_3$) of **1**

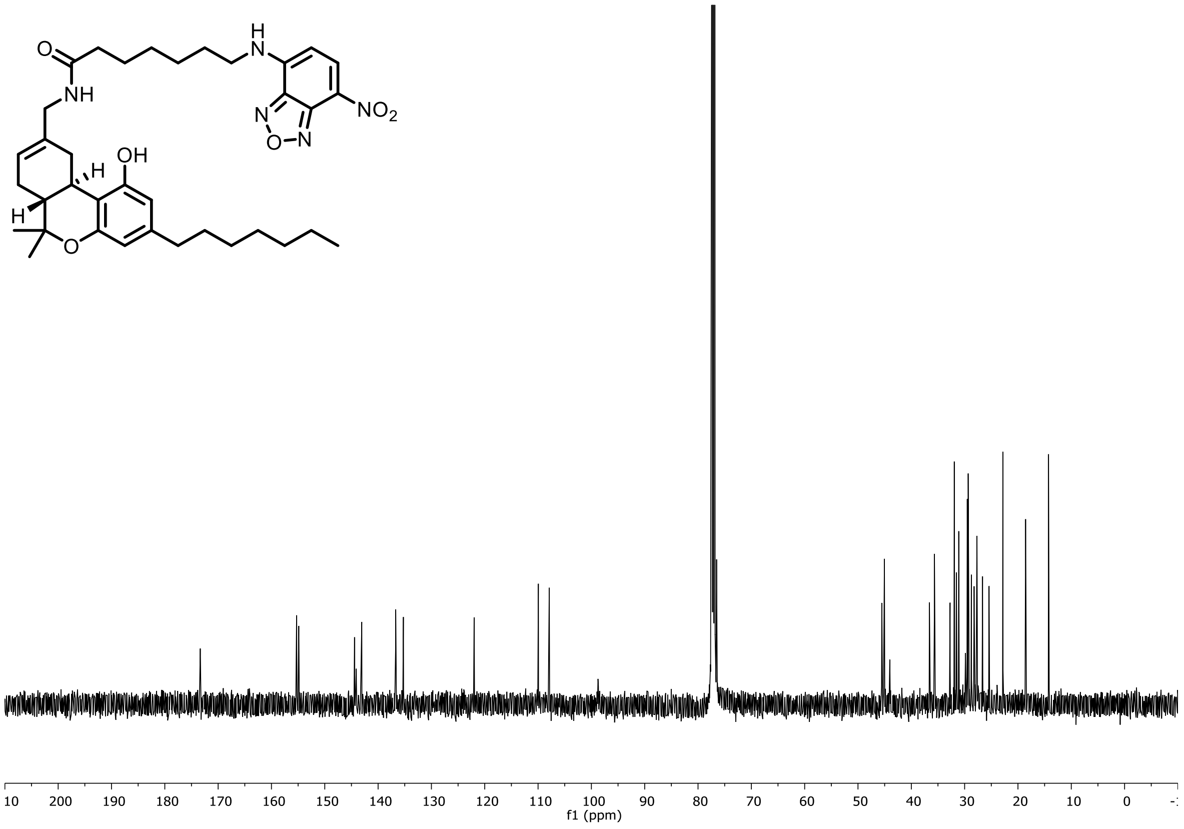

$^{1}$H NMR (400 MHz, $CDCl_3$) of **SI-10**

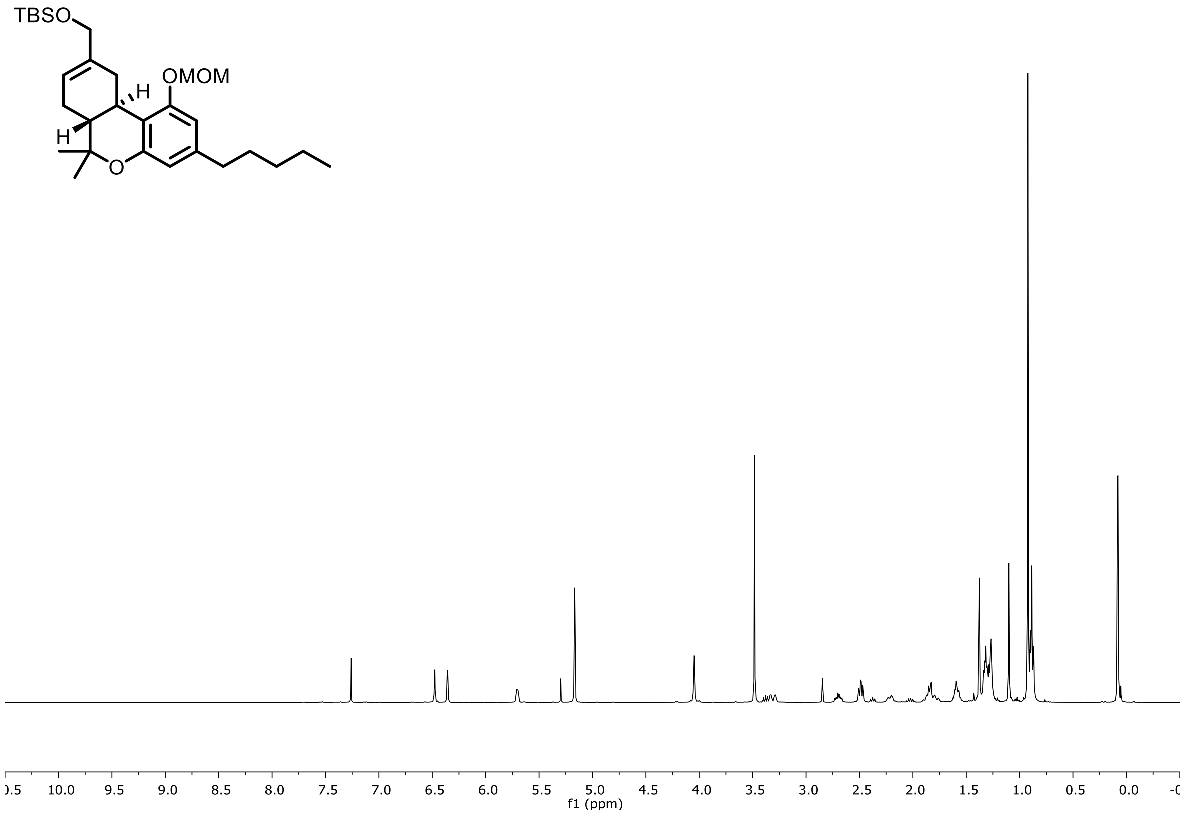


$^{13}$C NMR (101 MHz, $CDCl_3$) of **SI-10**

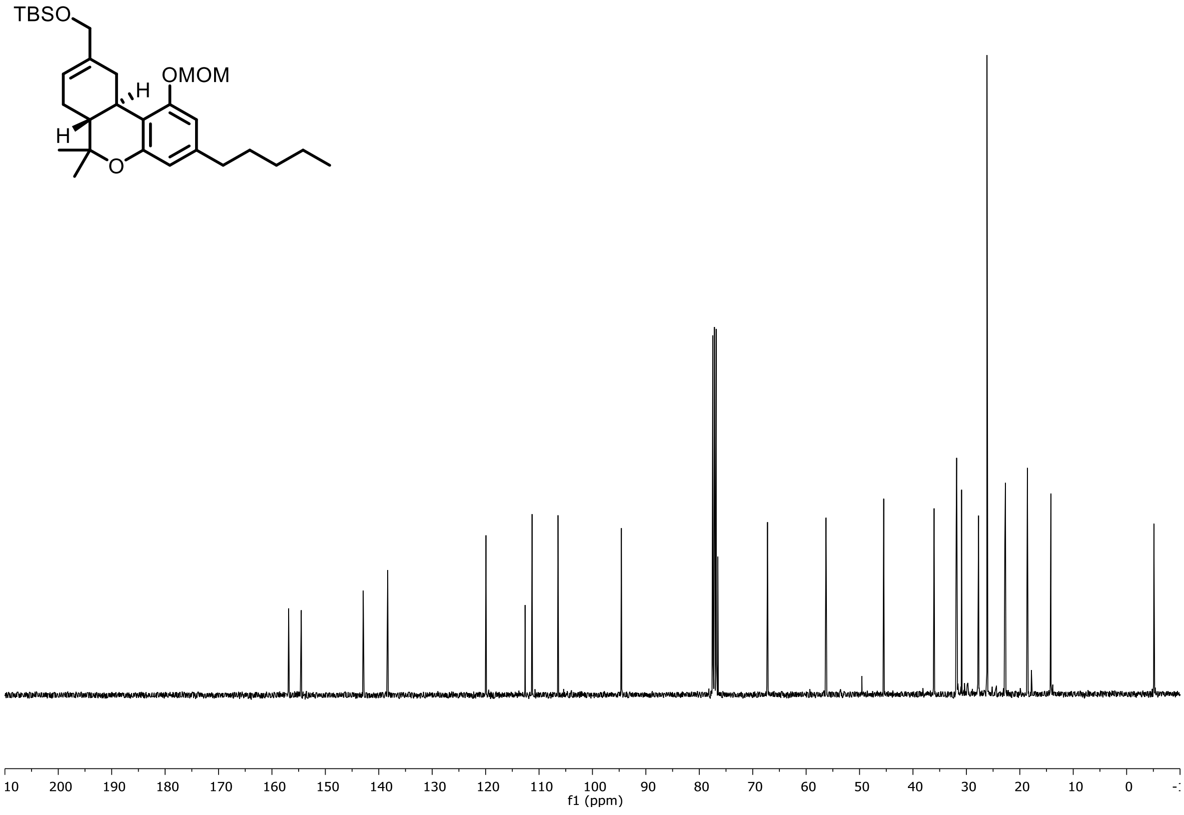

$^{1}H$ NMR (400 MHz, $CDCl_3$) of **SI-11**

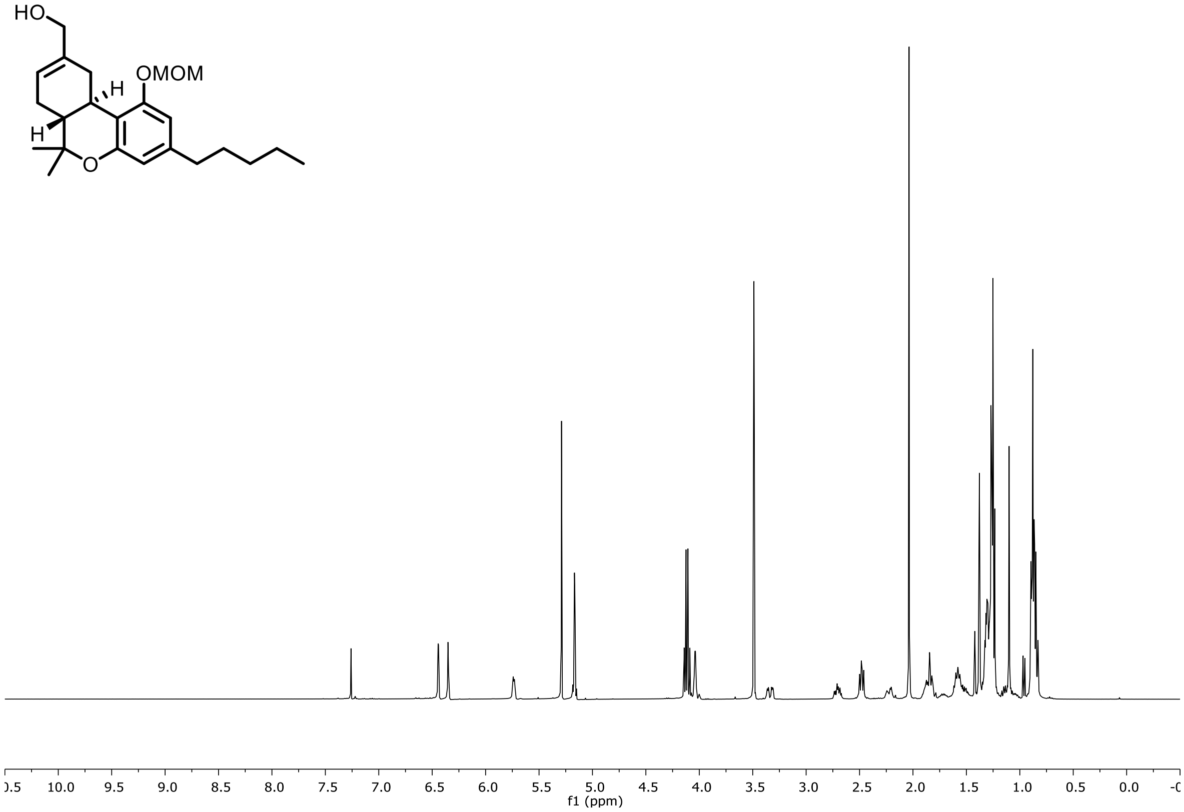


$^{13}C$ NMR (101 MHz, $CDCl_3$) of **SI-11**

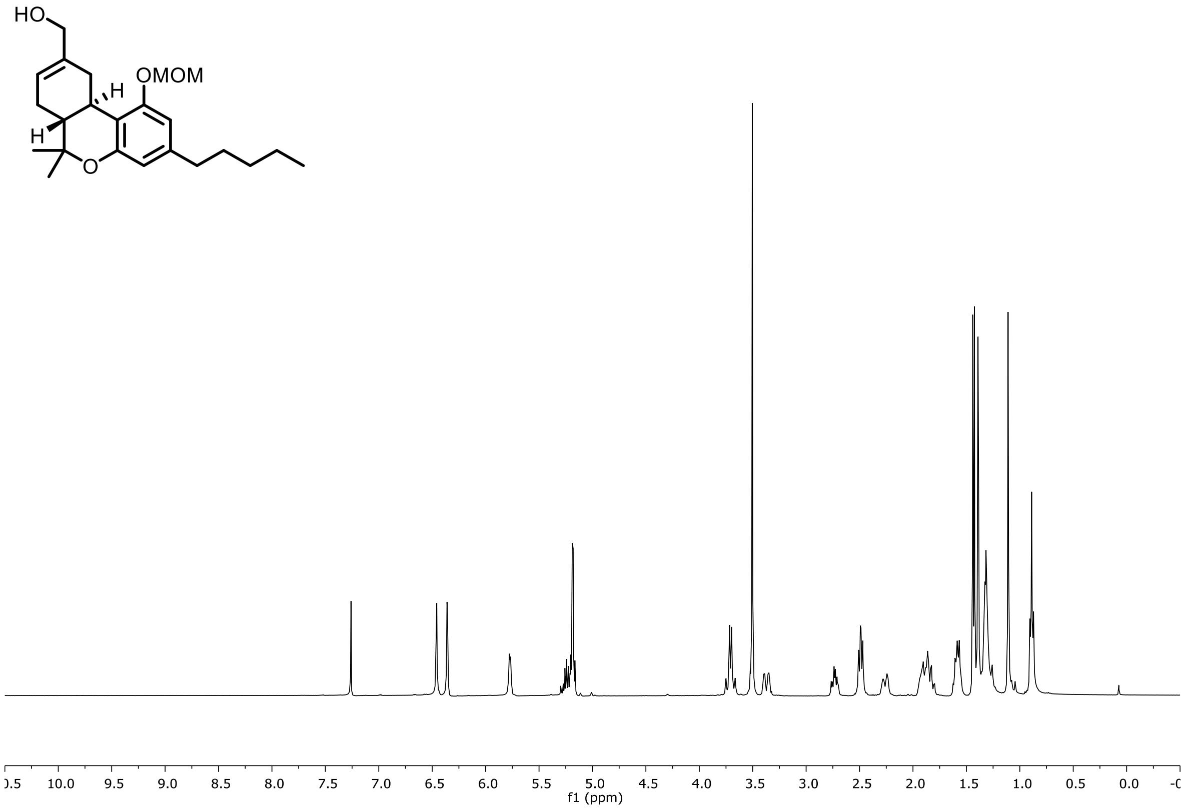

$^{1}H$ NMR (400 MHz, $CDCl_3$) of **SI-12**

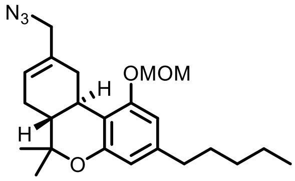


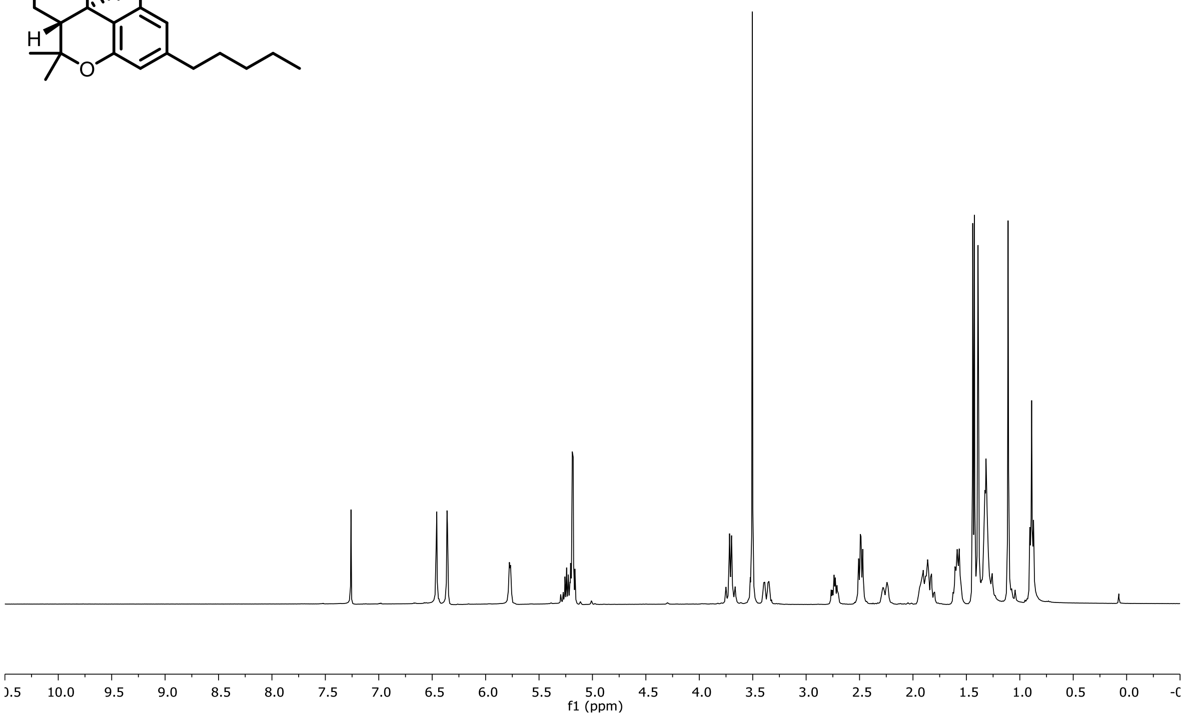


$^{13}C$ NMR (101 MHz, $CDCl_3$) of **SI-12**

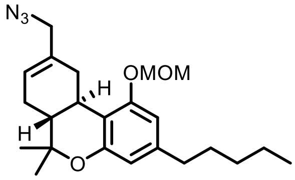


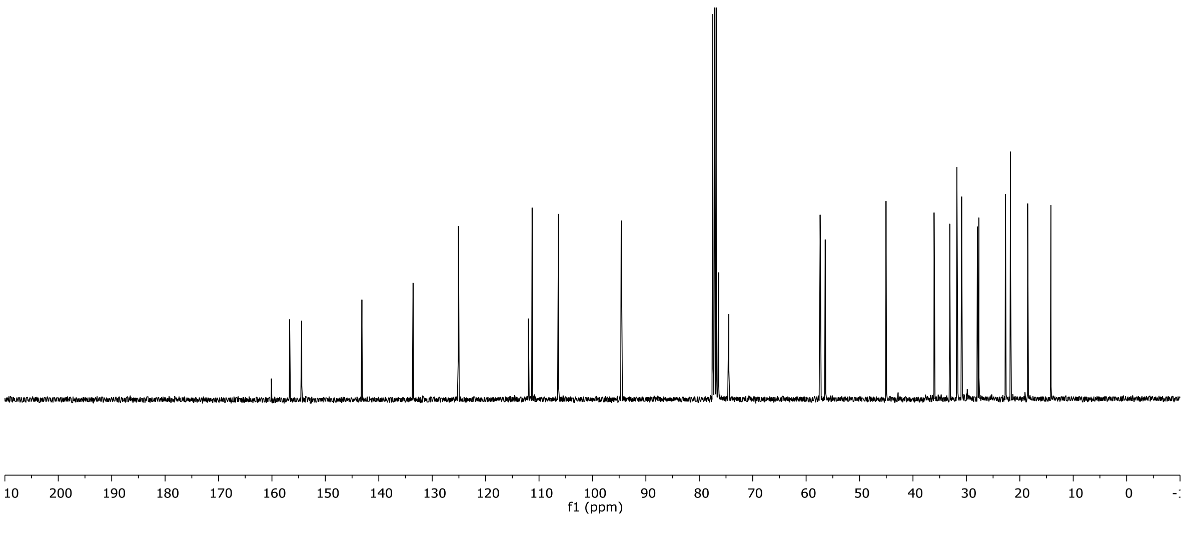

$^{1}H$ NMR (500 MHz, $CD_2Cl_2$) of **3**

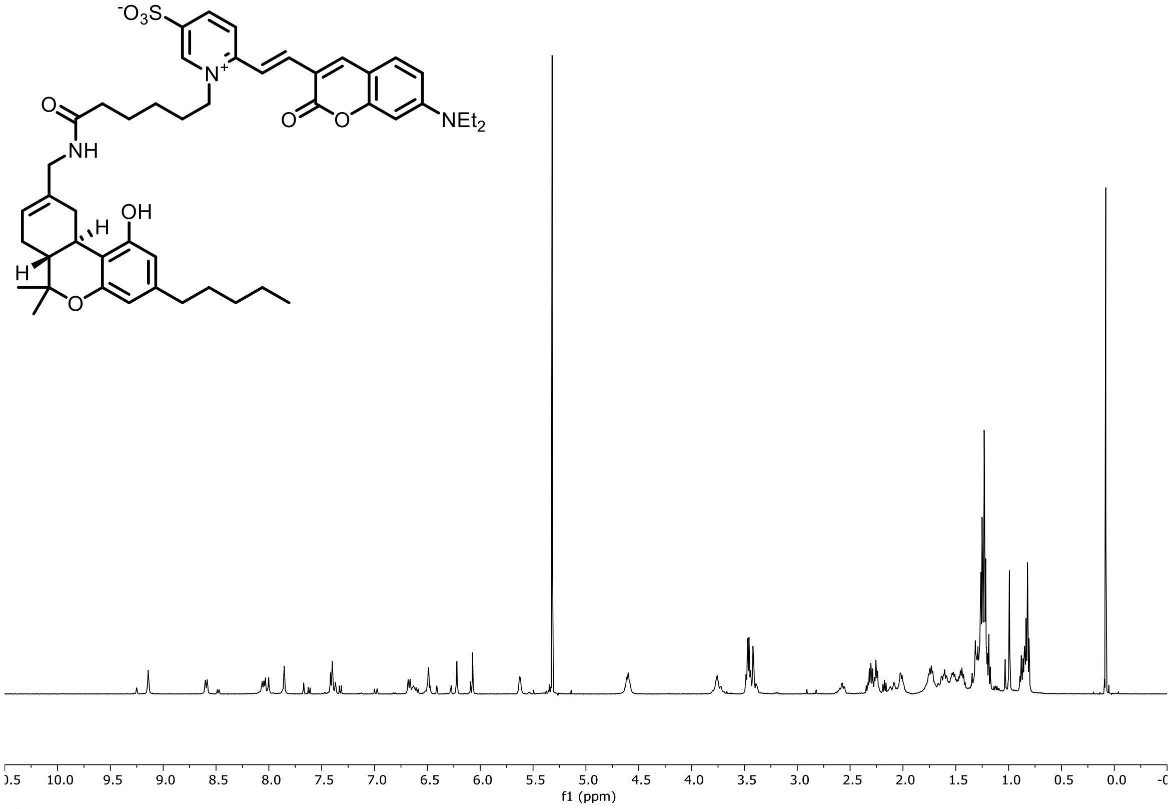


$^{13}C$ NMR (126 MHz, $CD_2Cl_2$) of **3**

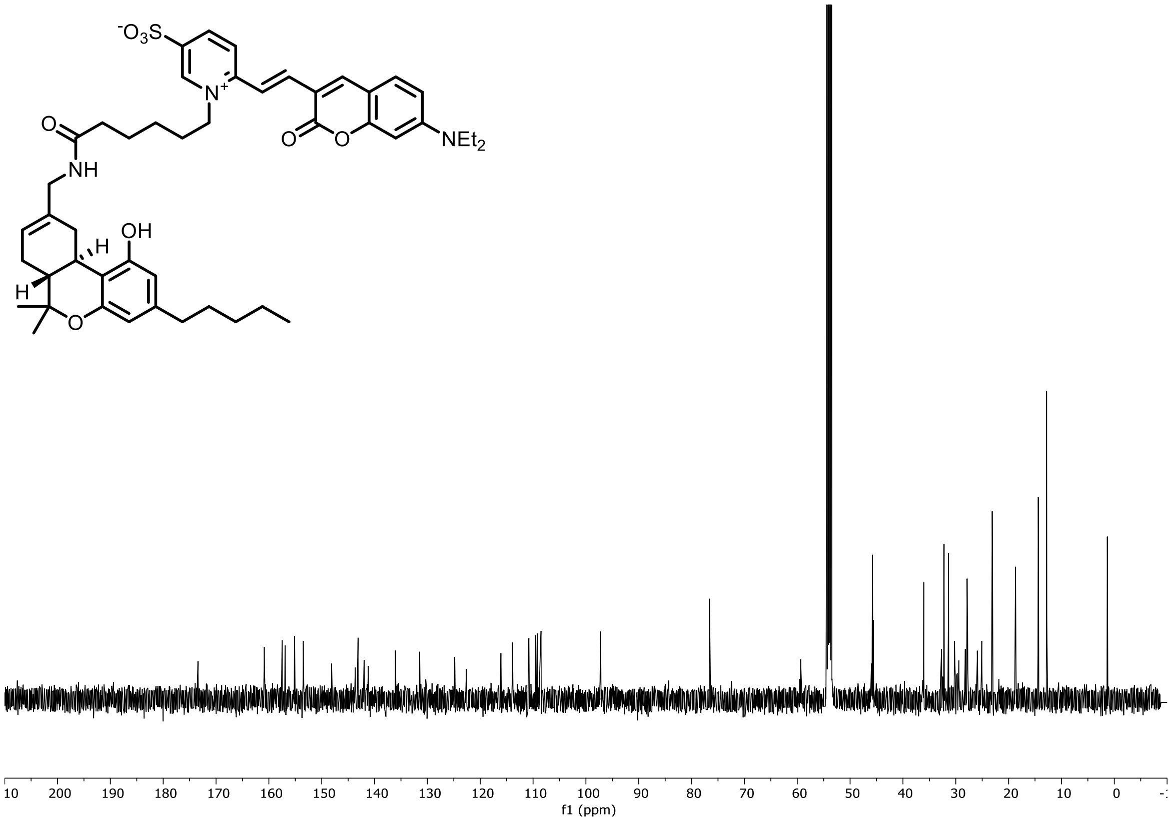

[1]H NMR (500 MHz, $CD_2Cl_2$) of **4**

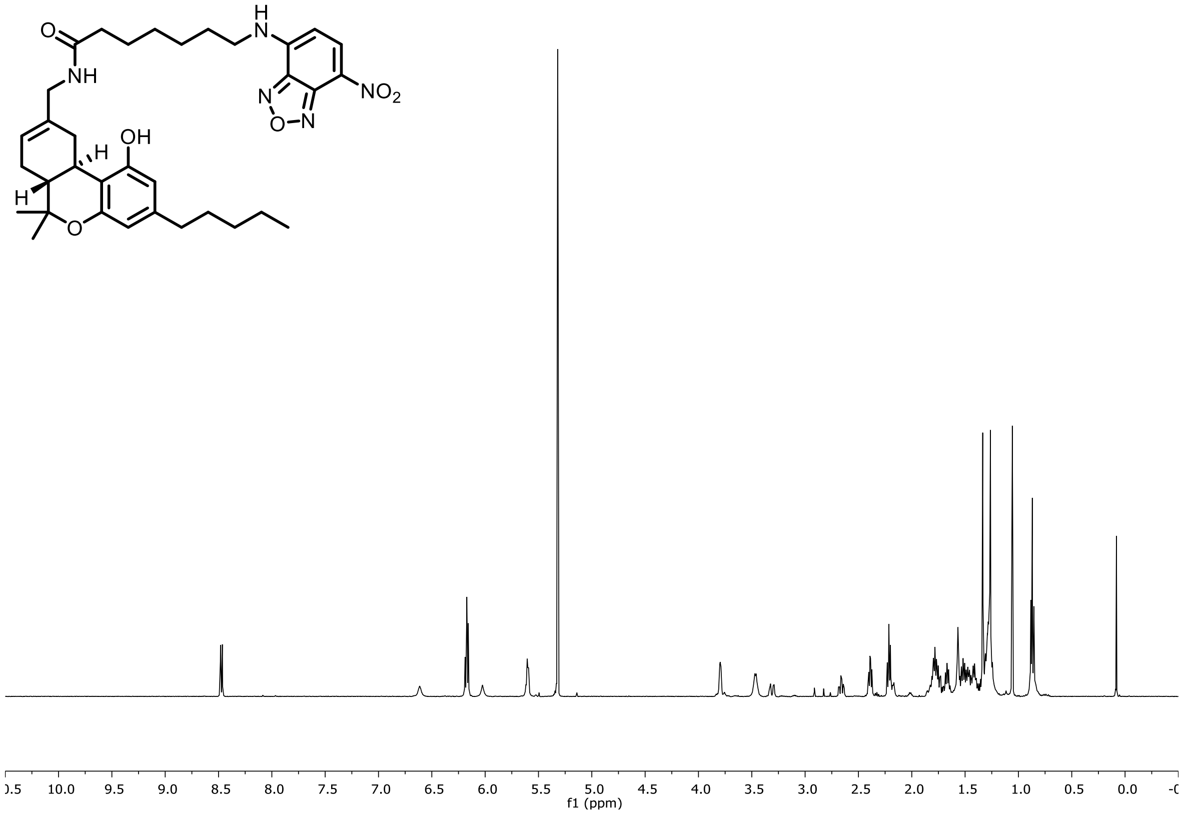


[13]C NMR (126 MHz, $CD_2Cl_2$) of **4**

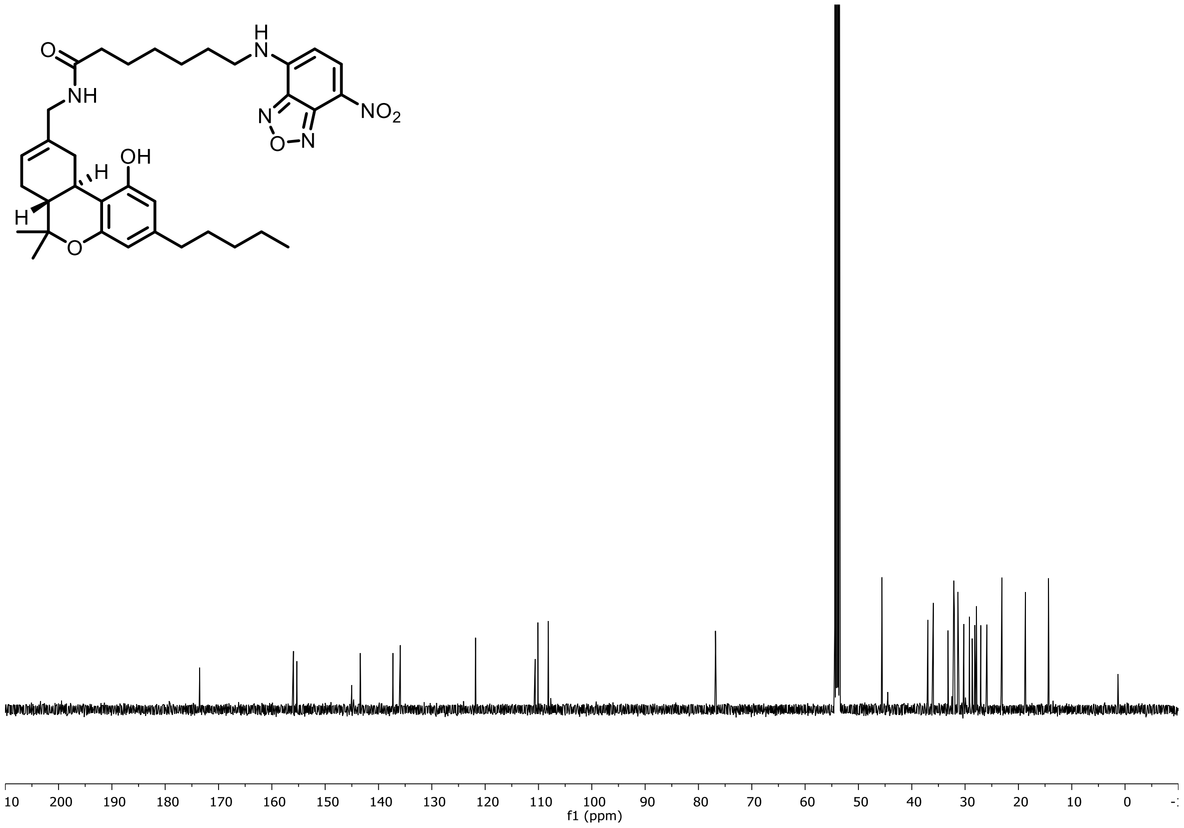

$^{1}H$ NMR (400 MHz, $CD_2Cl_2$) of **2**

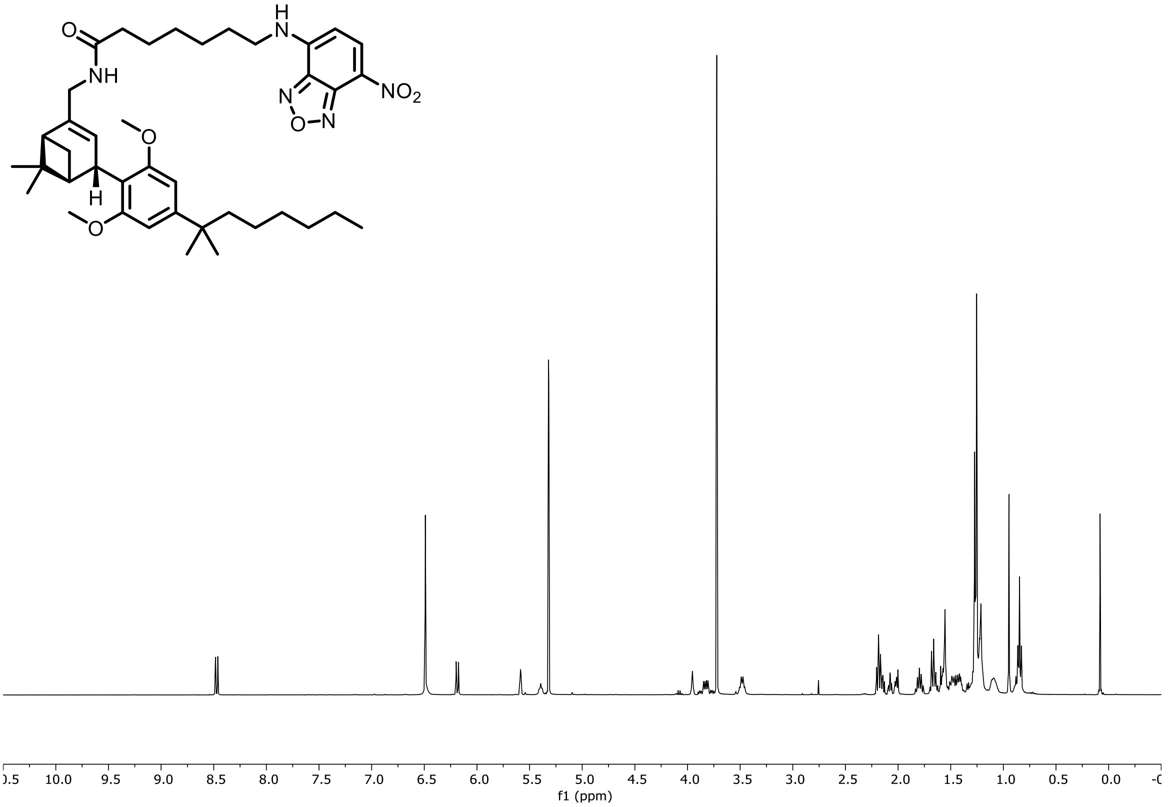


$^{13}C$ NMR (101 MHz, $CD_2Cl_2$) of **2**

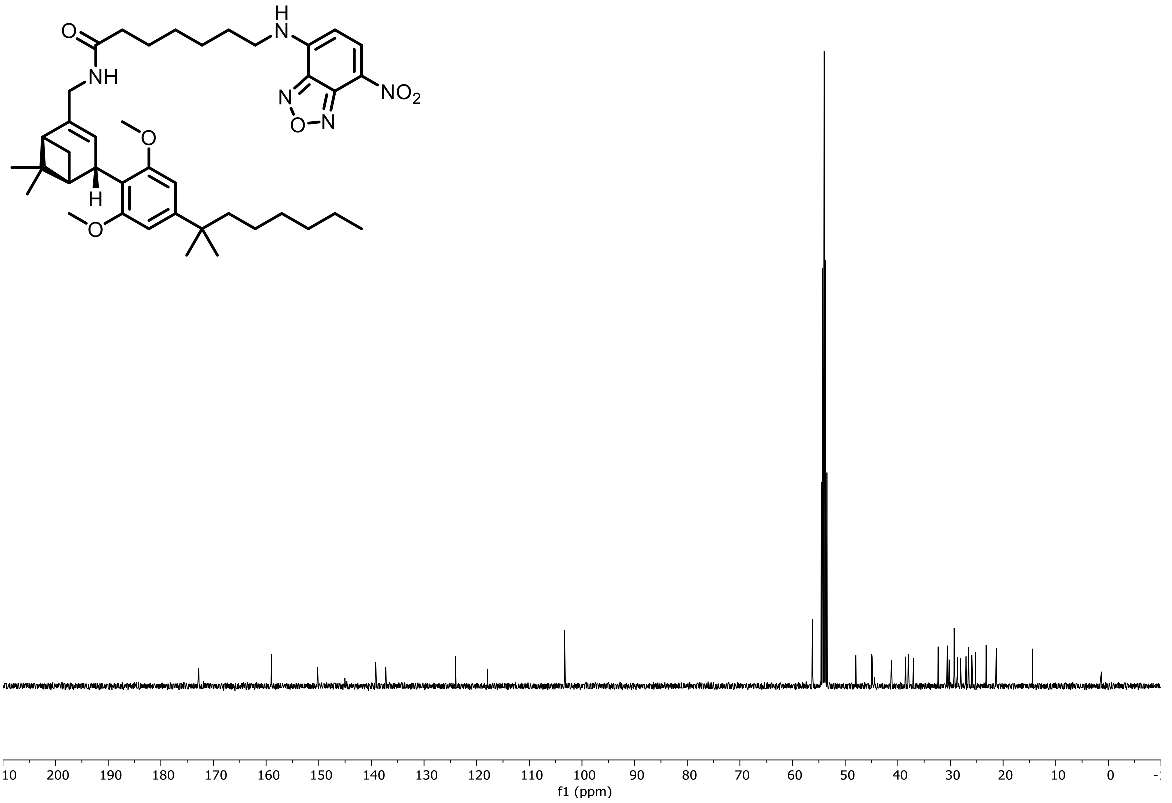

[1]H NMR (400 MHz, $CDCl_3$) of **SI-15**

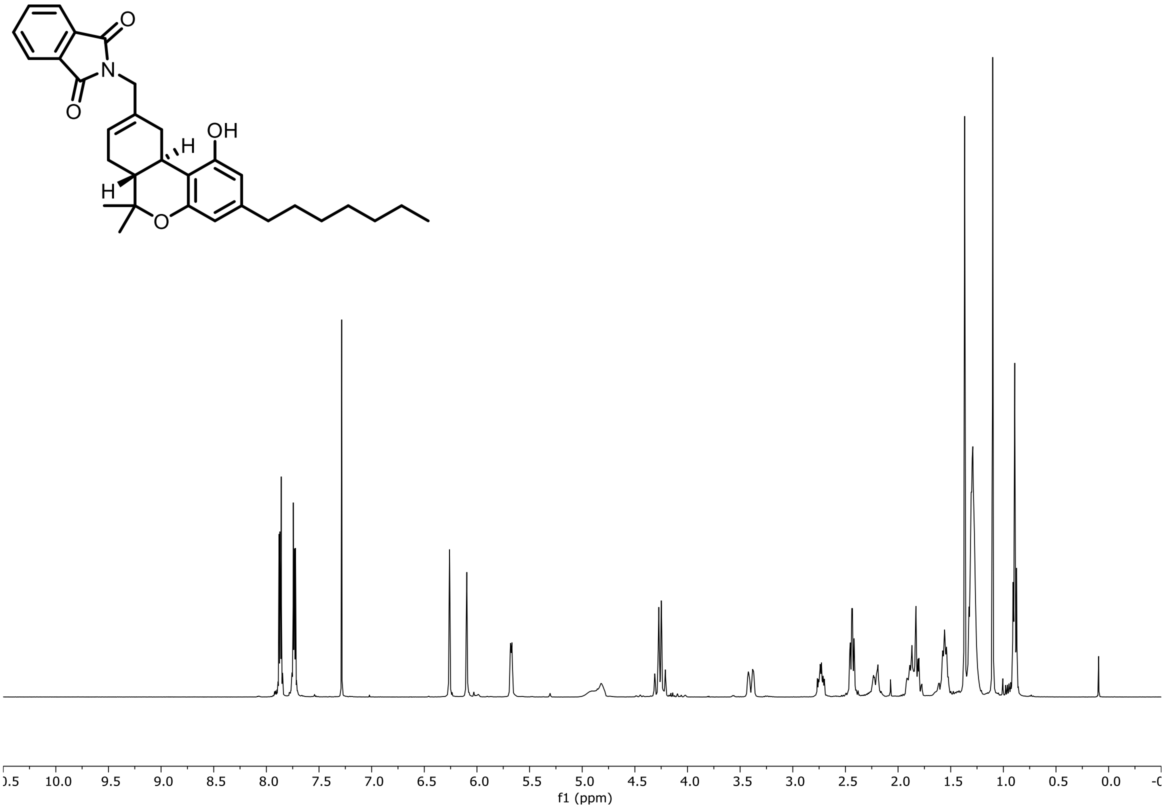


[13]C NMR (101 MHz, $CDCl_3$) of **SI-15**

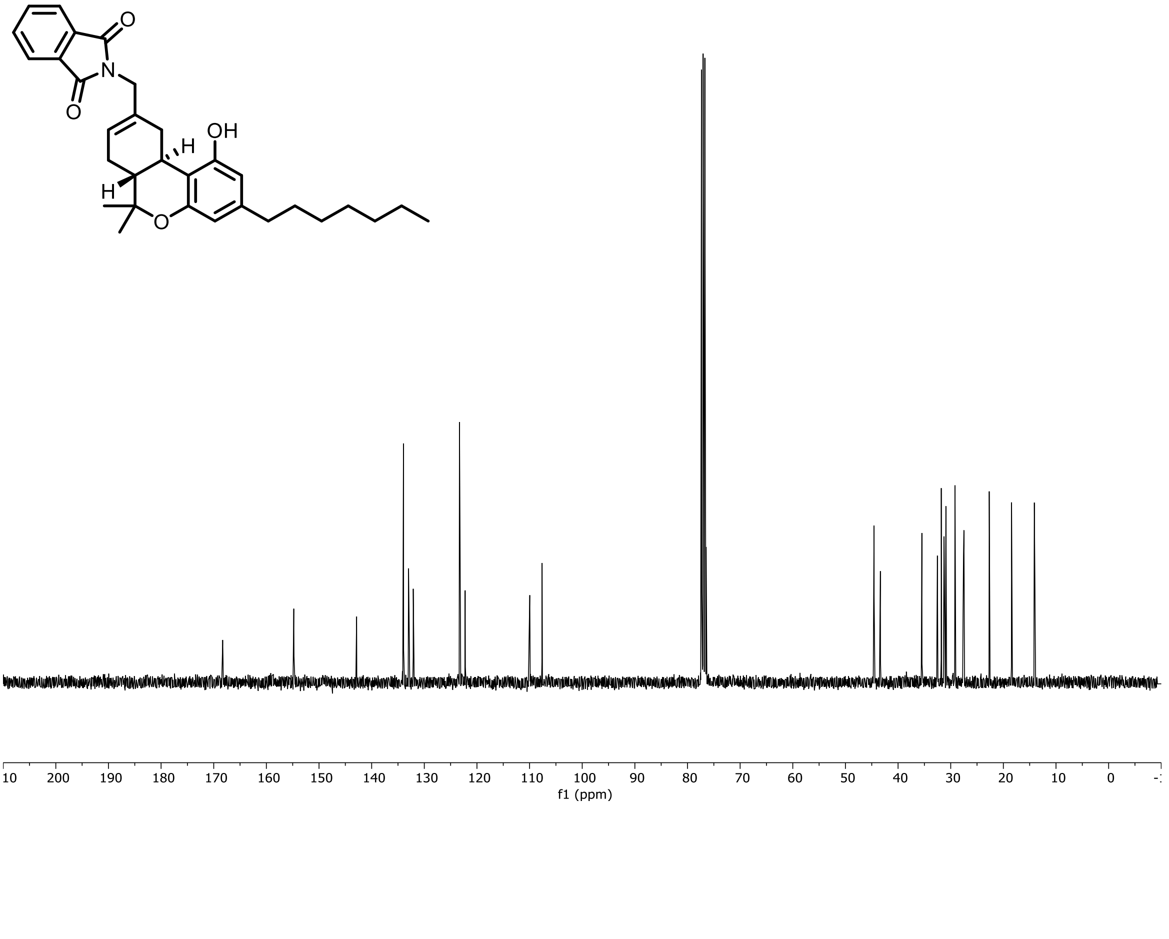

$^{1}H$ NMR (500 MHz, $CD_3OD$) of **SI-16**

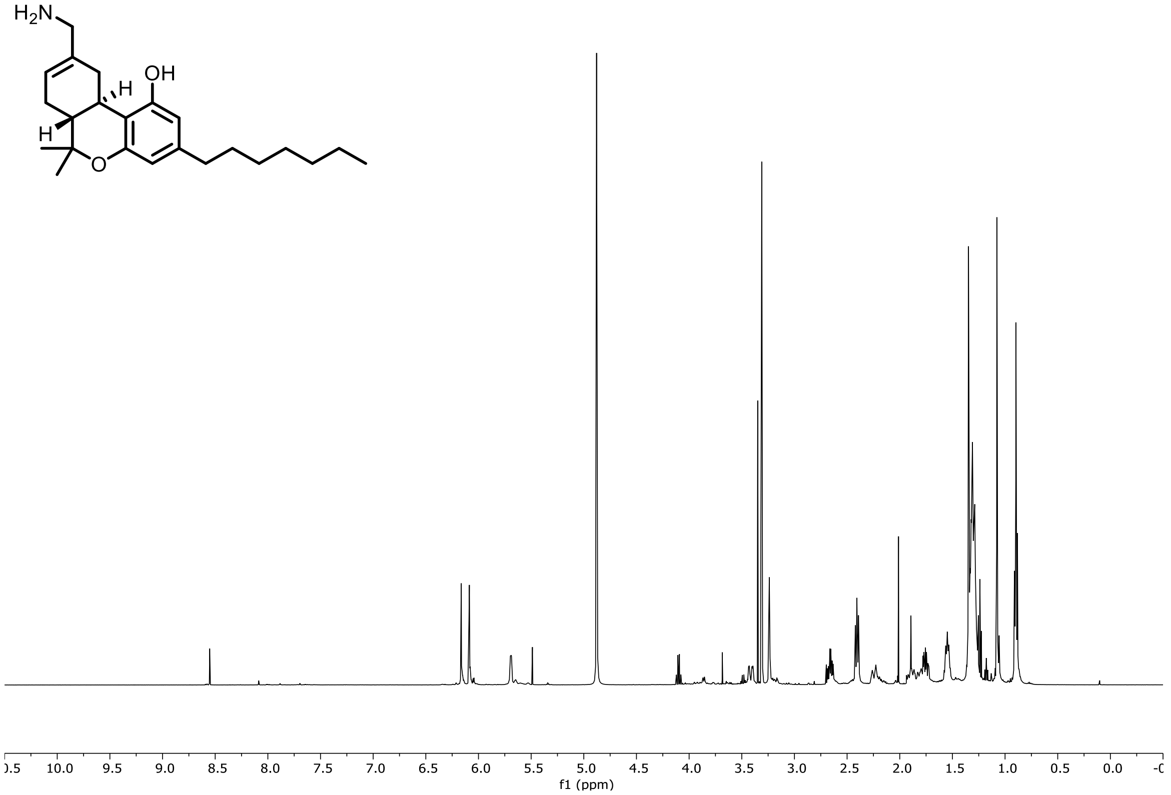


$^{13}C$ NMR (126 MHz, $CD_3OD$) of **SI-16**

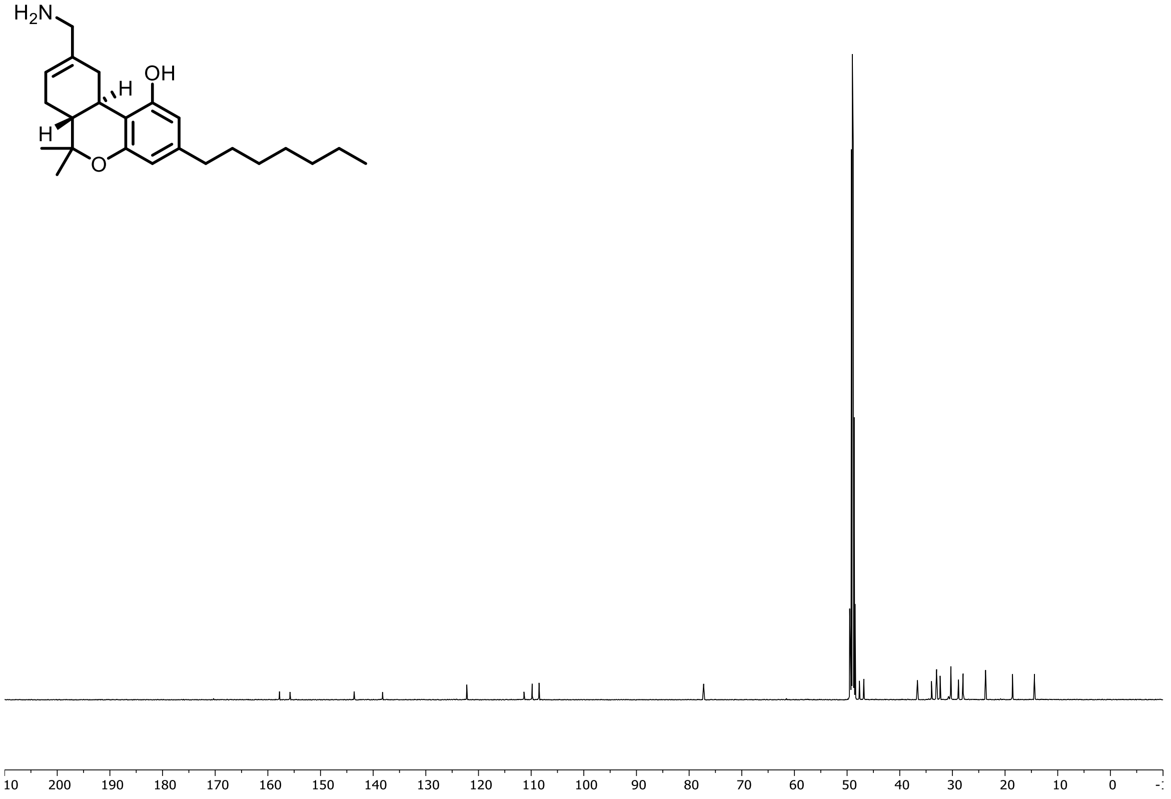
